# Non-Canonical Odor Coding in the Mosquito

**DOI:** 10.1101/2020.11.07.368720

**Authors:** Meg A. Younger, Margaret Herre, Olivia V. Goldman, Tzu-Chiao Lu, Gabriela Caballero-Vidal, Yanyan Qi, Zachary N. Gilbert, Zhongyan Gong, Takeshi Morita, Saher Rahiel, Majid Ghaninia, Rickard Ignell, Benjamin J. Matthews, Hongjie Li, Leslie B. Vosshall

## Abstract

Female *Aedes aegypti* mosquitoes are a persistent human foe, transmitting arboviruses including dengue and yellow fever when they bite us to obtain a blood meal. Mosquitoes are intensely attracted to human-emitted body odor, heat, and carbon dioxide, which they detect using three different large multi-gene families encoding odor-gated ion channels. Genetic mutations that cause profound disruptions to the olfactory system have modest effects on human attraction, suggesting significant redundancy in odor coding. The canonical view is that olfactory sensory neurons each express a single chemosensory receptor that defines its ligand selectivity. Using immunostaining, RNA *in situ* hybridization, and single nucleus RNA sequencing, we discovered that *Aedes aegypti* uses an entirely different organizational principle, with many neurons co-expressing multiple chemosensory receptor genes. *In vivo* electrophysiology demonstrates that the broad ligand-sensitivity of mosquito olfactory neurons is due to this non-canonical co-expression. The redundancy afforded by an olfactory system in which many neurons co-express multiple receptors with different chemical sensitivity may greatly increase the robustness of the mosquito olfactory system and explain our longstanding inability to engineer new compounds that disrupt the detection of human body odor by mosquitoes.

## INTRODUCTION

Increased global travel, a growing world population, and rising temperatures increase the emergence and transmission of novel disease-causing pathogens spread by “vector” organisms such as mosquitoes, ticks, sandflies, and fleas. Diseases spread by these arthropods collectively account for more than 700,000 deaths every year (WHO, 2020). Female *Aedes aegypti* mosquitoes spread arboviruses including dengue, Zika, yellow fever, and chikungunya. Only female mosquitoes bite, and they do so because they require a blood-meal for reproduction (Allan et al., 1987). *Aedes aegypti* prefer to bite human hosts, which contributes to their effectiveness as a disease vector (Brown et al., 2014; Gouck, 1972; McBride et al., 2014). To identify human hosts, mosquitoes rely heavily on chemosensory cues, including carbon dioxide (CO_2_) emitted from breath, and human body odor, which is a mixture of hundreds of different individual odorants including alcohols such as 1-octen-3-ol and volatile amines such as ammonia (Acree et al., 1968; Bernier et al., 2000; Cook et al., 2011; Davis, 1984; Gallagher et al., 2008; Geier et al., 1999; Kline, 1994; Smallegange et al., 2005; Smith et al., 1970). Insects detect such chemosensory cues using receptors encoded by three large multi-gene families, Odorant Receptors (ORs), Ionotropic Receptors (IRs), and Gustatory Receptors (GRs). All three gene families encode ionotropic ligand-gated ion channels, in contrast to the metabotropic seven transmembrane domain G protein-coupled odorant receptors utilized by vertebrates and *Caenorhabditis elegans* nematodes (Ihara et al., 2013). ORs are odorant-gated ion channels (Butterwick et al., 2018; Del Mármol et al., 2021; Sato et al., 2008; Wicher et al., 2008) that are formed by a heteromultimeric complex of the conserved co-receptor Orco and a ligand-selective OR (Benton et al., 2006; Larsson et al., 2004; Neuhaus et al., 2005; Sato et al., 2008). IRs are variant ionotropic glutamate receptors that are formed by one or more of three conserved co-receptors, Ir25a, Ir8a, and Ir76b, and ligand-selective subunits that determine the range of odorants detected by the receptor complex (Abuin et al., 2011; Benton et al., 2009; Silbering et al., 2011). Although GRs are primarily taste receptors (Clyne et al., 2000; Montell, 2009; Scott et al., 2001), some GRs detect temperature (Ni et al., 2016), and several GRs form a complex that detects carbon dioxide (CO_2_) in a variety of insects (Jones et al., 2007; Kwon et al., 2007). CO_2_ is an important volatile human host cue that activates and attracts mosquitoes (Gillies, 1980). In *Aedes aegypti*, *Gustatory Receptor 3* (*Gr3*) encodes an essential subunit of the CO_2_ receptor, and *Gr3* mutant mosquitoes lose all sensitivity to CO_2_ (McMeniman et al., 2014).

Because mosquitoes specialize on humans and require blood to reproduce, the drive to find humans is strong and innate. Indeed, even mosquitoes genetically engineered to eliminate genes critical for peripheral detection of host sensory cues can find and bite people. Animals lacking the Odorant receptor co-receptor (*Orco*), the obligate co-receptor required for the function of the entire family of ORs, show strong attraction to humans (DeGennaro et al., 2013). Deleting *Ir8a, Ir76b,* or *Ir25a* co-receptors reduces but does not eliminate attraction to humans (De Obaldia et al., 2022; Raji et al., 2019). Similarly, while mosquitoes lacking the obligate CO_2_ receptor subunit *Gr3* do not respond to CO_2_ and show impaired behavioral responses in laboratory assays, they are highly effective in finding humans in a more naturalistic semi-field setting (McMeniman et al., 2014). Although the exact odor profile of people varies considerably, *Aedes aegypti* are highly effective in finding humans to bite, despite widespread efforts by humans to mask our odor with chemical repellents (Tawatsin et al., 2006; Travis et al., 1949). We have yet to identify long-lasting interventions to prevent this deadly biting behavior, and it is not known how the mosquito olfactory system is seemingly infallible in its ability to detect humans.

The cloning of the first odorant receptors in 1991 (Buck and Axel, 1991) led to the subsequent discovery that each vertebrate olfactory sensory neuron expresses a single odorant receptor that specifies its functional properties. With few exceptions, the well-studied olfactory system of *Mus musculus* mice features olfactory sensory neurons that are thought to express a single olfactory receptor (Bashkirova and Lomvardas, 2019; Chess et al., 1994). The same organization was reported in *Drosophila melanogaster* flies (Clyne et al., 1999; Gao and Chess, 1999; Vosshall et al., 1999), although recent work challenges this model (McLaughlin et al., 2021; Task et al., 2021). In both species, decades of evidence has supported the model that neurons expressing a given receptor project axons to dedicated olfactory glomeruli in the first sensory processing center in the brain, the antennal lobe in insects (Couto et al., 2005; Fishilevich and Vosshall, 2005; Vosshall et al., 2000) and the olfactory bulb in vertebrates (Mombaerts et al., 1996; Ressler et al., 1994; Vassar et al., 1994). This “one-receptor-to-one-neuron-to-one-glomerulus” organization is believed to be a widespread motif in insect olfactory systems, and the convergence onto discrete glomeruli is hypothesized to permit the brain to utilize combinatorial coding and parse which subpopulation of olfactory neurons is activated by a given odorant (Bisch-Knaden et al., 2018; Semmelhack and Wang, 2009; Wang et al., 2003).

Consistent with this “one-receptor-to-one-neuron-to-one-glomerulus” organization in insects, the number of expressed chemosensory receptors in the OR and IR gene families in many insects roughly correlates to the number of olfactory glomeruli. This holds true in the honey bee *Apis mellifera* (∼180 receptors/∼160 glomeruli) (Flanagan and Mercer, 1989; Robertson et al., 2010), the tobacco hornworm *Manduca sexta* (∼60 receptors/∼70 glomeruli) (Grosse-Wilde et al., 2011), and *Drosophila melanogaster* flies (∼60 receptors/∼55 glomeruli) (Benton et al., 2009; Laissue et al., 1999; Robertson et al., 2003). Based on these studies, it is widely thought that merely counting the number of antennal lobe glomeruli in a new species would be reasonably predictive of the number of chemosensory receptors found in its genome. In *Aedes aegypti*, however, there is a striking mismatch between the number of expressed chemosensory receptors and the number of antennal lobe glomeruli, with at least twice as many receptors as available glomeruli (Bohbot et al., 2007; Ignell et al., 2005; Matthews et al., 2018; Shankar and McMeniman, 2020; Zhao et al., 2020). This leads to the question of how the mosquito olfactory system is organized to accommodate so many receptors and whether this deviation from rules established in other species explains their exquisite ability to locate human hosts.

In this study, we developed a CRISPR-Cas9-based genetic knock-in strategy in *Aedes aegypti* to generate genetically-modified mosquito strains that label molecularly distinct populations of olfactory sensory neurons. We used these strains to understand how the mosquito olfactory system is organized and discovered that OR- and IR-expressing olfactory sensory neurons frequently innervated the same antennal lobe glomeruli. To ask if this was a feature of individual olfactory neurons expressing multiple chemosensory receptors, we profiled receptor expression in peripheral sensory organs using RNA *in situ* hybridization and by immunostaining with antibodies that recognize endogenous OR and IR co-receptors. To complement these studies, we carried out single nucleus RNA sequencing to profile gene expression in the antennae and maxillary palps. Through these experiments, we found that the olfactory system of *Aedes aegypti* does not have the expected “one-receptor-to-one-neuron-to-one-glomerulus” organization seen in other organisms. We frequently observed co-expression of multiple chemosensory receptors from at least two of the three receptor gene superfamilies within individual olfactory sensory neurons. We also saw expression of multiple receptors from a single family within the same olfactory sensory neuron. To test if multiple receptors function to detect different ligands within the same olfactory sensory neuron, we used *in vivo* electrophysiology to examine odorant responses in the maxillary palp. We discovered a class of neurons that expresses members of both the OR and IR gene family and that responds to odorants that activate either OR or IR pathways. When we mutated either the OR or IR pathway by deleting the major co-receptors, neurons retained responsivity to the odorant sensed by the pathway that was still intact. Therefore, both ORs and IRs are required to detect different classes of odorants in the same sensory neuron. This sensory organization, in which multiple receptors responding to different chemosensory stimuli are co-expressed, suggests a redundancy in the code for human odor. We speculate that this unconventional organization underlies the robust, seemingly unbreakable properties of the *Aedes aegypti* olfactory system in detecting human odor and driving human host-seeking in this olfactory specialist.

## RESULTS

### Mismatch in chemosensory receptor and olfactory glomerulus number suggests a novel olfactory organization

In the mosquito, olfactory cues are sensed by olfactory sensory neurons in the antenna and the maxillary palp, whose axons project to the ipsilateral antennal lobe of the brain (Distler and Boeckh, 1997; Ignell et al., 2005) (Figure 1A-D, Figure S1A-C). The antennal lobe, the insect equivalent of the vertebrate olfactory bulb, is organized into discrete olfactory glomeruli in which axons from peripheral olfactory sensory neurons terminate and synapse with local interneurons and projection neurons that relay olfactory information to the higher brain (Stocker, 1994). Previous studies used morphological criteria to define 50 (Ignell et al., 2005), 60 (Zhao et al., 2020), or 81 (Shankar and McMeniman, 2020) discrete olfactory glomeruli in the female *Aedes aegypti* antennal lobe. In this study, we define approximately 65 olfactory glomeruli (64.9 ± 0.9, mean±SEM), obtained by counting antennal lobe glomeruli in the left hemisphere of 12 female *Aedes aegypti* brains stained to reveal synaptic neuropil (Figure 1B,I,K, Figure S2-5). The glomerulus count ranged from 60-72 glomeruli per antennal lobe, indicating a high level of variability in the organization of the antennal lobe. We generated 3-D reconstructions of complete antennal lobes and saw considerable variability in the size and shape of the glomeruli (Figure S1). We were able to consistently identify certain landmark glomeruli, most notably the three glomeruli that are innervated by the maxillary palp (Ignell et al., 2005; Shankar and McMeniman, 2020) (Figure S1D-K).

**Figure 1:**
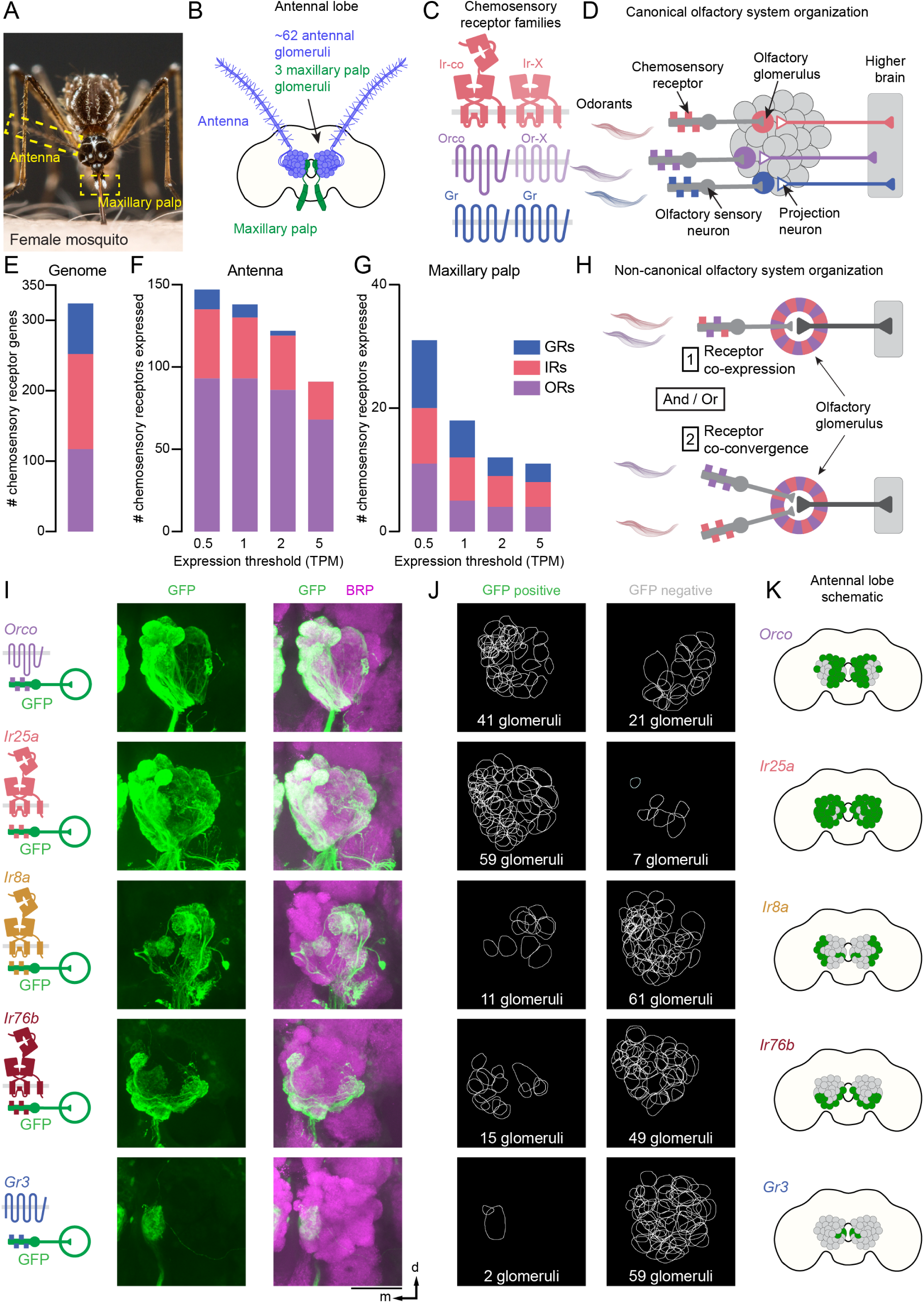
Mismatch in chemosensory receptor and olfactory glomerulus number. (**A**) *Aedes aegypti* female with sensory structures (yellow boxes). (**B**) Approximate number of antennal lobe glomeruli per brain hemisphere innervated by the indicated sensory structure, derived from quantification of the left antennal lobe in 12 brains presented in (I-J) and Figure S2-S5. See also Figure S1. (**C**) Cartoons of insect chemosensory gene families. (**D**) Cartoon of canonical olfactory system organization. (**E-G**) Stacked bar plots of the number of chemosensory genes in the *Aedes aegypti* genome (E), and the number expressed above the indicated TPM thresholds in the antenna (F) and maxillary palp (G). (**H**) Two models of non-canonical olfactory system organization. (**I**) Maximum-intensity projections of confocal Z-stacks of antennal lobes in the left brain hemisphere of the indicated genotype with immunofluorescent labeling of GFP (green) and the nc82 monoclonal antibody, which recognizes Brp (magenta). Brp is used throughout this paper as a synaptic marker (Wagh et al., 2006). Scale bar: 50 µm. Orientation: d=dorsal, m=medial. (**J**) 2-D representation of the boundary of each glomerulus in (I) that is GFP positive or GFP negative. See also Figure S2-S6. (**K**) Cartoon schematic of the antennal lobe regions receiving projections from olfactory sensory neurons expressing the indicated chemosensory receptor.

The canonical “one-receptor-to-one-neuron-to-one glomerulus” organization posits that the number of chemosensory receptors should roughly match the number of glomeruli in the antennal lobe (Figure 1D). While there is not yet a clear consensus on the number of olfactory glomeruli in *Aedes aegypti*, it ranges from 50 to 81. How does this relate to the number of chemosensory receptors expressed? In the updated *Aedes aegypti* L5 genome (Matthews et al., 2018), there are 117 OR, 135 IR, and 72 GR genes for a total of 324 structural genes that could function in the olfactory system (Figure 1E). Reanalysis of previously published antennal and maxillary palp RNA-sequencing data (Matthews et al., 2016) using multiple expression thresholds demonstrates that even at the conservative threshold of 5 transcripts per million (TPM), the mosquito olfactory system expresses 102 chemosensory receptors, and adjusting the threshold to 2, 1, or 0.5 TPM increases the number of receptors plausibly expressed to 134, 156, and 178, respectively (Figure 1F,G). Thus, there are many more chemosensory receptors expressed in the olfactory system than available antennal lobe glomeruli, suggesting that the organization of the *Aedes aegypti* olfactory system must differ from the canonical scheme. We speculate that the mismatch can be resolved by expressing multiple receptors per neuron or having multiple molecularly distinct neurons co-converge on a single glomerulus or both (Figure 1H).

To begin to distinguish between these two organizational principles, we generated a collection of CRISPR-Cas9 gene-targeted strains that label subpopulations of olfactory neurons using the Q-system, a binary expression system similar to Gal4/UAS (Brand and Perrimon, 1993) that uses cell type-specific expression of the QF2 transcription factor to induce expression of an effector from the QF2 binding *QUAS* enhancer (Potter et al., 2010; Riabinina et al., 2015; Riabinina et al., 2016). We introduced an in-frame insertion that replaced the stop codon of each of the co-receptors *Orco*, *Ir25a*, *Ir8a*, and *Ir76b*, as well as the CO_2_ receptor subunit *Gr3* with the transcription factor *QF2* (Figure 1I, Figure S2-S6. See Data File 1 for a full description of all genotypes by figure) (Matthews et al., 2019; Potter et al., 2010; Riabinina et al., 2016). These gene-sparing knock-in strains were designed to cause minimal disruption to the locus to increase the likelihood that they would faithfully report expression of the endogenous gene. We crossed these *QF2* driver lines individually to a *QUAS-CD8:GFP* reporter to label neuronal membranes and visualized axonal projection patterns in the antennal lobe.

*Orco*, *Ir25a*, *Ir8a*, and *Ir76b* co-receptor driver lines were expressed in olfactory sensory neurons with distinct projection patterns in the antennal lobe (Figure 1I-K). Unexpectedly, neurons that expressed *Ir25a* projected to almost all glomeruli in the antennal lobe (89.9 ± 1.4%, mean±SEM, n=3) (Figure 1I-K, Figure S3), and expression overlapped extensively with glomeruli labeled by *Orco* (Figure 1I-K, Figure S2, Figure S3). While these co-receptor driver lines labeled glomeruli in the same regions from brain to brain, the interindividual expression patterns were not identical, consistent with the variability in glomerular arrangement that we have observed. Neurons that detect CO_2_ are located in the maxillary palp (Grant et al., 1995; Lu et al., 2007; Omer and Gillies, 1971) and we saw that *Gr3*-expressing neurons projected to a large glomerulus in the posterior antennal lobe, Glomerulus 1 (Figure 1I-K) which is also innervated by *Ir25a*-expressing neurons. We also noted the presence of a second small glomerulus that was often innervated by *Gr3*-expressing neurons in the antenna (Figure S6B). These initial findings point to the overlap of *OR-*, *IR-*, and *GR*-expressing neurons in the antennal lobe of *Aedes aegypti*, consistent with recent observations in *Drosophila melanogaster* (Task et al., 2021).

### Co-expression of *Orco* and *Ir25a* in the mosquito olfactory system

The high degree of overlap between glomeruli labeled by *Orco*- and *Ir25a*-expressing olfactory sensory neurons suggests that there is either widespread *Orco* and *Ir25a* co-expression within individual sensory neurons or that *Orco* and *Ir25a* are expressed in different neurons whose axons co-converge onto individual antennal lobe glomeruli or both (Figure 1H). To determine if *Orco* and *Ir25a* are co-expressed, we adapted the Split-QF2 system (Riabinina et al., 2019) for use in the mosquito. This system “splits” the transcription factor QF2 into two components, the DNA binding domain (QF2-DBD) and the activation domain (QF2-AD) each tagged with a synthetic leucine zipper (Figure 2A,B). When both the QF2-DBD and QF2-AD are co-expressed in the same cell, the two domains associate via the leucine zipper, reconstitute a functional QF2 protein, initiate transcription at the *QUAS* enhancer, and drive expression of a reporter gene (Figure 2C).

**Figure 2.**
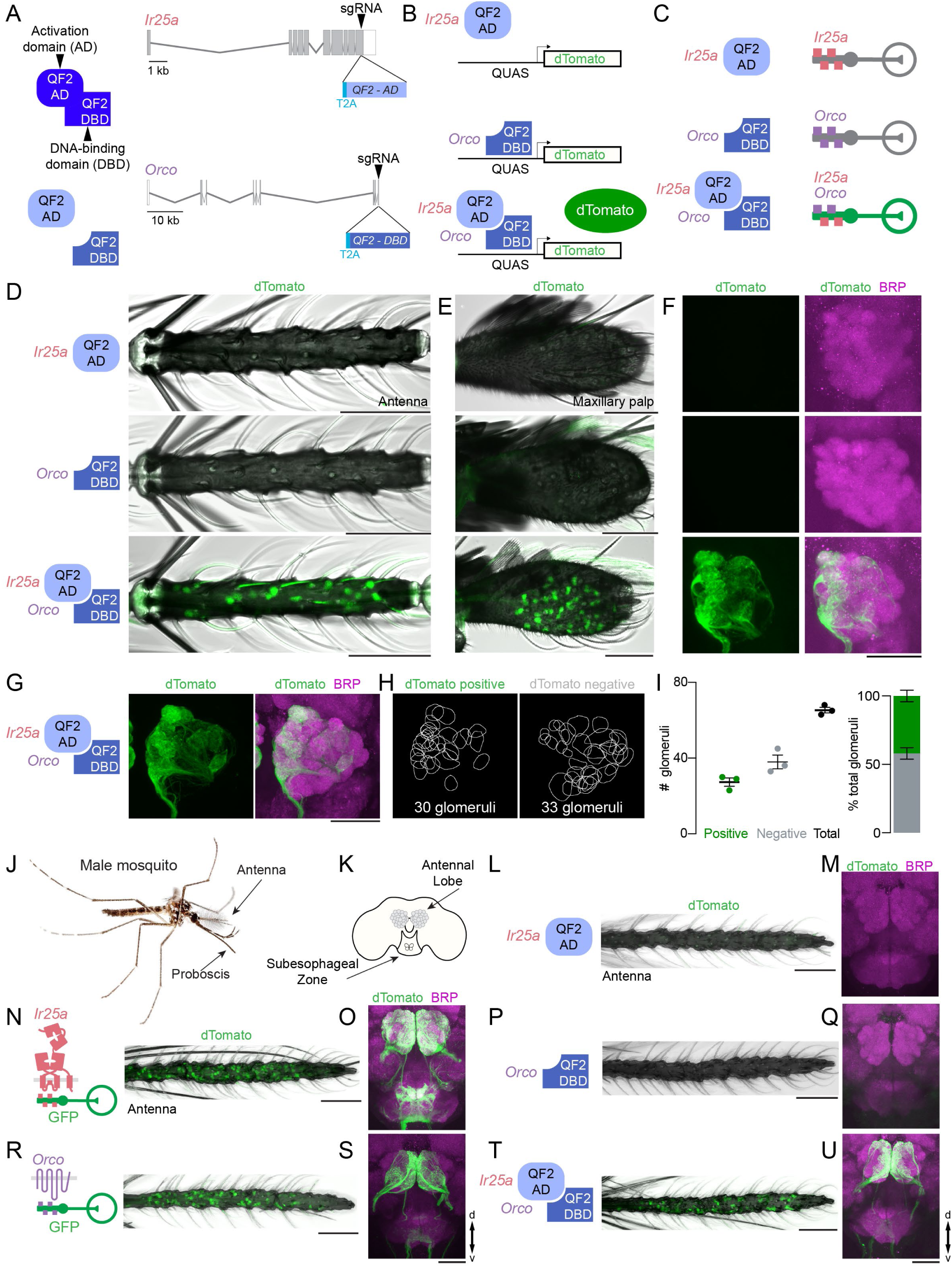
Genetic evidence for widespread *Orco* and *Ir25a* co-expression. (**A**) Schematic of the Split-QF2 system (left) and diagrams of *Orco* and *Ir25a* gene loci with exons (grey boxes), introns (grey lines) and CRISPR-Cas9 gRNA site (arrowhead) used to insert *T2A-QF2-AD* (light blue) and *T2A-QF2-DBD* (medium blue). AD and DBD gene maps are not to scale. (**B-C**) Schematic of the Split-QF2 system (B) and outcome of gene expression in olfactory sensory neurons of the indicated genotypes (C). (**D-E**) Maximum-intensity projections of confocal Z-stacks of female antennae (D) and female maxillary palps (E) of the indicated genotypes showing intrinsic dTomato fluorescence, with transmitted light overlay. See also Figure S7A,B. (**F-G**) Maximum-intensity projections of confocal Z-stacks of antennal lobes from the left brain hemisphere of the indicated genotype with immunofluorescent labeling of dTomato (green) and Brp (synaptic marker, magenta). See also Figure S8. (**H-I**) 2-D representation of the boundary of each glomerulus in (G) that is GFP positive or GFP negative (H) and quantification (I). Data are presented as mean±SEM, n=3. See also Figure S7C. (**J**) *Aedes aegypti* male with sensory structures (arrows). (**K**) Cartoon schematic of the brain including the antennal lobe glomeruli and the suboesophageal zone. (**L-U**) Maximum-intensity projections of confocal Z-stacks of male antennae (L,N,P,R,T) and male brains (M,O,Q,S,U) of the indicated genotype with immunofluorescent labeling of dTomato (green) and Brp (synaptic marker, magenta). Scale bars: 50 µm. Orientation: proximal left (D, E, L, N, P, R, T), medial left (F,G); d=dorsal, v=ventral, m=medial.

Using the same stop-codon replacement approach that we used to generate the *QF2*-lines, we inserted the *QF2-AD* into the *Ir25a* locus to generate *IR25a-QF2-AD* and the *QF2-DBD* into the *Orco* locus to generate *Orco-QF2-DBD*. When either *IR25a-QF2-AD* or *Orco-QF2-DBD* was used to drive expression of *dTomato* (Shaner et al., 2004), we did not see fluorescence in the female antenna, maxillary palp, or the antennal lobe (Figure 2D-F, Figure S7). Therefore, neither QF2-DBD nor QF2-AD alone can activate expression from the *QUAS* enhancer. However, when *Orco-QF2-DBD* and *IR25a-QF2-AD* were crossed into the same animal, we saw expression of *dTomato* in antennal and maxillary palp neurons of female mosquitoes, as well as axonal projections in the antennal lobe (Figure 2D-F, Figure S7, Figure S8). Nearly half of the glomeruli in the antennal lobe were labelled with dTomato (Figure 2G-I, Figure S7, Figure S8). This points to widespread *Orco* and *Ir25a* co-expression within *Aedes aegypti* olfactory sensory neurons, although we note that these findings do not rule out the possibility that co-convergence may be present as well.

We then examined the olfactory system of male mosquitoes (Figure 2J-U). We observed extensive expression of *dTomato* throughout the antenna and in axons that terminate in the antennal lobe when we drove expression with either the *Orco-QF2* or *IR25a-QF2* driver lines (Figure 2N,O,R,S). We again saw no expression of *dTomato* when driven by either the *IR25a-QF2-AD* or *Orco-QF2-DBD* control lines (Figure 2L,M,P,Q). However, when *Orco-QF2-DBD* and *IR25a-QF2-AD* were crossed into the same animal, we observed widespread *dTomato* expression in the male antenna and antennal lobe, indicating co-expression of *Ir25a* and *Orco* in male olfactory sensory neurons (Figure 2T-U). It is not currently possible to compare male and female glomerular position due to differences in antennal lobe volume and shape and lack of glomerulus-specific driver lines, however, the general expression pattern was similar between males and females, with innervation predominant in the anterior-medial antennal lobes (Figure 2F,G,U).

### Co-expression of *Orco* and *Ir25a* in the mosquito taste system

Another source of olfactory information in *Aedes aegypti* may derive from olfactory neurons on the proboscis, the mouthpart of the mosquito that engages in taste and food ingestion. *Drosophila melanogaster* flies express IRs and GRs in the proboscis, but not ORs (Larsson et al., 2004). In contrast, *Orco* neurons are widespread in both the proboscises of *Anopheles gambiae* and *Aedes aegypti* mosquitoes, and RNA-sequencing data from *Aedes aegypti* has shown there are many ligand-selective ORs expressed in this taste tissue (Matthews et al., 2016; Riabinina et al., 2016). Cells in the proboscis have been shown to respond to volatile odorants in *Anopheles gambiae*. However, projections from these neurons in *Anopheles gambiae* and *Aedes aegypti* extend to the subesophageal zone, the taste processing center of the brain, and not to the antennal lobe (Ghaninia et al., 2007; Jové et al., 2020; Kwon et al., 2006; Riabinina et al., 2016). It remains unknown if the brain interprets these cues as olfactory or gustatory.

We used our QF2 and Split-QF2 reagents to reveal the expression of *Orco* and *Ir25a* in proboscis neurons and examined how these cells innervate the subesophageal zone. There was extensive expression of both *Orco* and *Ir25a* alone in the proboscis (Figure 3A) as well as co-expression of *Orco* and *Ir25a* as defined by dTomato expression in the Split-QF2 animals (Figure 3C). *Ir25a*-expressing neurons send extensive projections to the subesophageal zone, with axons terminating in the anterior and posterior regions of the subesophageal zone. There is a small cluster of glomeruli in the central subesophageal zone that receives dense innervation as well (Figure 3B). *Orco*-expressing neurons do not project to the anterior region and send sparse projections to the posterior subesophageal zone and subesophageal zone glomeruli (Figure 3B). Innervation by the neurons that co-express both *Orco* and *Ir25a* send projections only to the posterior-ventral subesophageal zone, with the densest innervation in the medial region and sparser lateral arborizations (Figure 3D).

**Figure 3.**
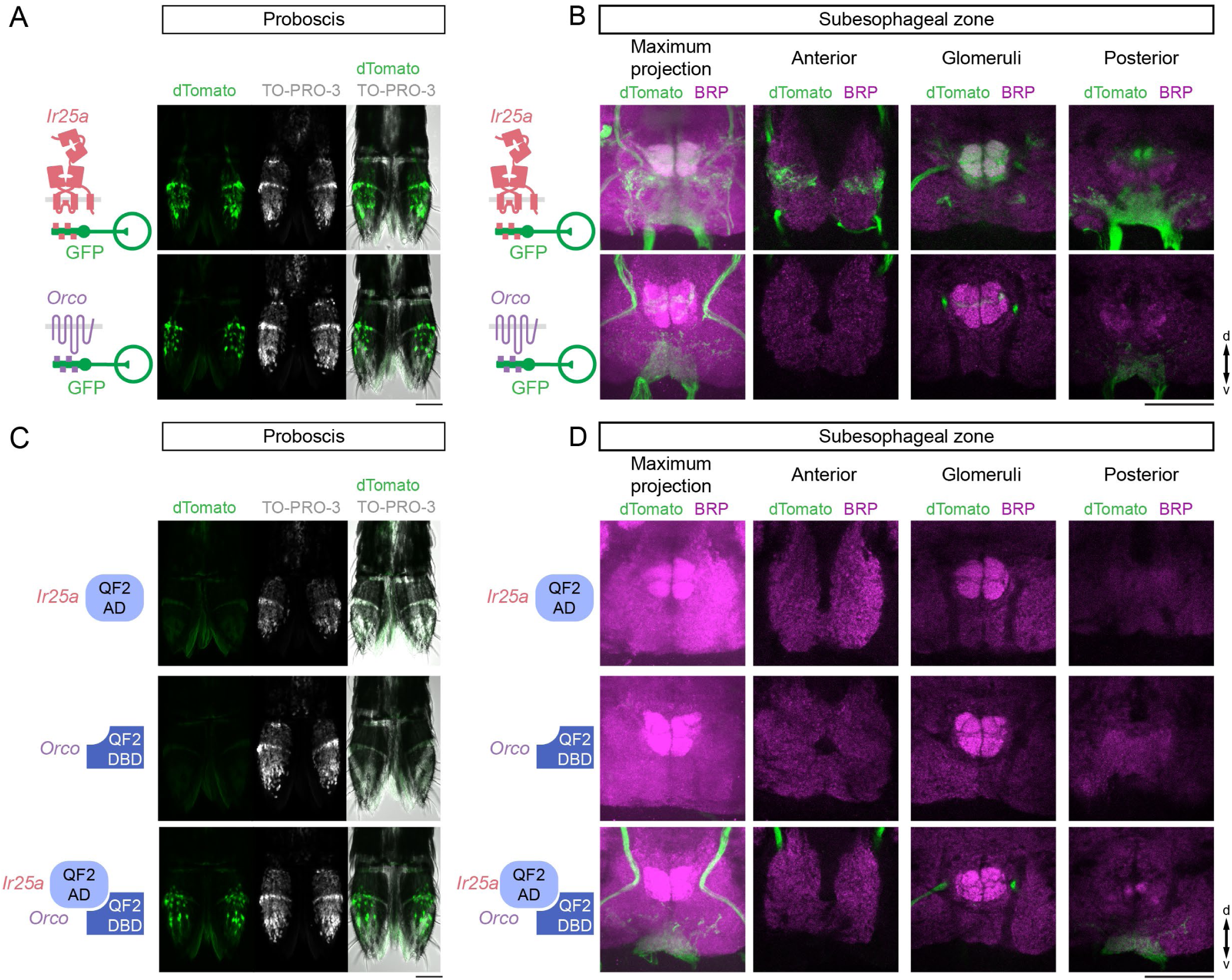
*Orco* and *Ir25a* co-expression in the mosquito proboscis. (**A, C**) Maximum-intensity projection of whole-mount dTomato (green) expression and TO-PRO-3 nuclear stain (white) in female proboscises of the indicated genotypes. (**B, D**) Left panel, maximum-intensity projections of confocal Z-stacks of suboesophageal zone from the indicated genotypes with immunofluorescent labeling of dTomato (green) and Brp (synaptic marker, magenta). Right three panels, single confocal sections through indicated areas of the subesophageal zone. Orientation: proximal up (A,C); d=dorsal, v=ventral (B,D). Scale bars: 50 µm.

Since *Ir25a* complexes mediate not only detection of volatile odorants but also gustatory cues, it is possible that sensory afferents in the *Aedes aegypti* proboscis are able to detect olfactory as well as gustatory information within the same neurons. Alternatively, IRs and ORs in these neurons may function as olfactory receptors that relay olfactory information to the taste center of the mosquito brain.

### Extensive co-expression of chemosensory co-receptors in the antenna

The observation that nearly half of antennal lobe glomeruli receive projections from neurons co-expressing *Ir25a* and *Orco* suggested that there is extensive co-expression of IRs and ORs throughout the antenna. To determine if neurons co-express Orco and Ir25a protein we generated an antibody to Ir25a and conducted whole mount antennal immunostaining with this antibody and a previously characterized Orco antibody (Basrur et al., 2020) to label endogenous Orco and Ir25a proteins in wild-type mosquitoes (Figure 4A-D). We observed extensive co-expression of Orco and Ir25a (Figure 4A-D), confirming the co-expression of these distinct chemosensory genes seen using our QF2 and Split-QF2 driver lines. In addition to neurons that contain both Orco and Ir25a protein, we also observed neurons that express either Orco or Ir25a alone (Figure 4A-D), indicating that OR cells, IR cells, and mixed OR+IR cells exist. We validated the specificity of the Orco and Ir25a antibodies by performing whole-mount immunostaining on antennae from *Orco* and *Ir25a* mutants (Figure 4E-H). To confirm and extend these results, we performed RNA *in situ* hybridization on wild-type antennae with probes designed to target endogenous *Orco, Ir76b,* and *Ir25a* RNAs (Figure 4I-K). These experiments replicated patterns of co-expression observed in immunostained antennae (Figure 4A-D), with almost half of the *Orco* cells co-expressing *Ir25a*, and vice versa. In contrast we saw that few *Orco* cells co-express *Ir76b* by RNA *in situ* hybridization. Both the RNA *in situ* hybridization and immunostaining data indicated that widespread co-expression is not an artifact of the QF2 and split-QF2 driver lines.

**Figure 4.**
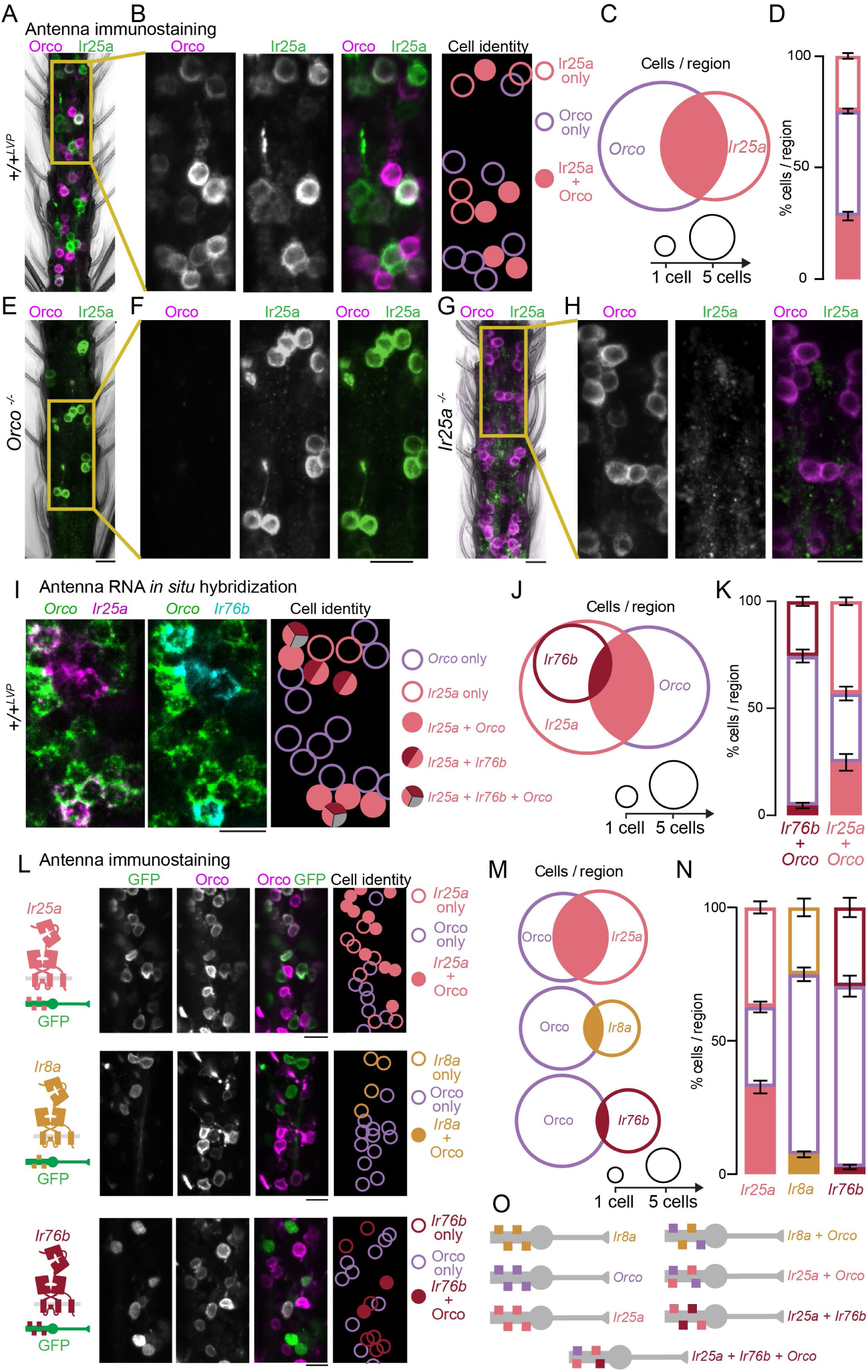
Extensive chemosensory co-receptor co-expression in the antenna. (**A**) Maximum-intensity projection of whole-mount Orco and Ir25a immunostaining in wild-type female antennae. (**B**) Enlarged view of the yellow rectangle in (A) with cartoon schematic indicating cell identity at the right. (**C,D**) Quantification of antennal cells in the indicated genotypes co-expressing Orco and Ir25a presented as Euler diagrams with area scaled to mean cells/region (C) and stacked bar plots (D). Data are presented as mean±SEM, n=7 antennal segments, 48-61 cells/region. (**E,G**) Maximum-intensity projection of whole-mount Orco and Ir25a immunostaining in *Orco^16/16^* mutant (**E**) and Ir25a^BamHI/BamHI^ mutant (**G**) female antennae. (**F,H**) Enlarged view of the yellow rectangles in (E,G). (**I**) RNA *in situ* hybridization in wild-type antennae with the indicated probes. (**J,K**) Quantification of wild-type antennal cells expressing the indicated genes as Euler diagrams with area scaled to mean cells/region (J), and stacked bar plots (K). Data are presented as mean±SEM, n=4 antennal segments, 45-63 cells/region. (**L**) Maximum-intensity projection of whole-mount Orco and GFP immunostaining in female antennae of the indicated genotypes with cartoon schematic indicating cell identity at the right. (**M,N**) Quantification of antennal cells in the indicated genotypes co-expressing Orco protein and GFP presented as Euler diagrams with area scaled to mean cells/region (M) and stacked bar plots (N). Data are presented as mean±SEM, n=6-8 antennal segments, 34-68 cells/region. (**O**) Cartoon schematic of olfactory sensory neuron populations identified in this figure. Scale bars: 10 µm.

To gain additional resolution on the degree of overlap between Orco and the three major IR family co-receptors, we carried out whole-mount antennal immunostaining with an antibody to the endogenous Orco protein and to GFP expressed from each sensory neuron QF2 driver. We confirmed extensive co-expression of Orco and *Ir25a* (Figure 4L-N) and found that substantially fewer cells co-express either Orco and *Ir8a* or Orco and *Ir76b*, even after accounting for fewer total *Ir76b* and *Ir8a* cells (Figure 4L-N). We also note that in addition to widespread co-receptor co-expression, some mosquito olfactory neurons express just one co-receptor (Figure 4O), highlighting the complexity in the rules that govern receptor co-expression in *Aedes aegypti* antennal olfactory sensory neurons.

### Single nucleus RNA sequencing reveals that many antennal neurons co-express multiple ligand-selective receptor subunits

Functional ORs and IRs are composed of a complex of co-receptor and ligand-selective receptor subunits. Because there are hundreds of ligand-selective OR and IR genes, it was not feasible to examine combinatorial co-expression of the full complement of receptors by RNA *in situ* hybridization or immunostaining. Instead, we developed a method for single nucleus RNA sequencing (snRNA-seq) in mosquito antennae based on previously described nucleus extraction protocols from *Drosophila melanogaster* antennae (Li et al., 2021; McLaughlin et al., 2021). We isolated antennae from female mosquitoes, extracted nuclei, performed droplet microfluidics to barcode reads from each cell, and performed droplet-based snRNA-seq using the 10x Genomics platform. For clarity of presentation, we use “cell” as the unit of analysis to refer to expression profiling of single nuclei. These experiments were carried out in two batches, Batch 1 at Rockefeller University and Batch 2 at Baylor College of Medicine (see Methods). We filtered for cells based on quality control parameters and combined data from two batches, to capture a total of 14,161 cells (Figure 5A, Figure S9A-G). Unsupervised clustering was used to categorize cells into broad subtypes, which revealed cells that express epithelial or glial markers (Figure 5B, Figure S9H). This analysis also yielded 19 neuron clusters based on expression of at least 3 of 4 neural markers (*CadN*, *brp*, *syt1*, and *elav*) in 50% or more of the cells within that cluster (Figure S9I). A total of 6,645 cells were classified as neurons. We then used unsupervised clustering on this population of neurons and identified 54 clusters of antennal neurons (Figure 5C).

**Figure 5:**
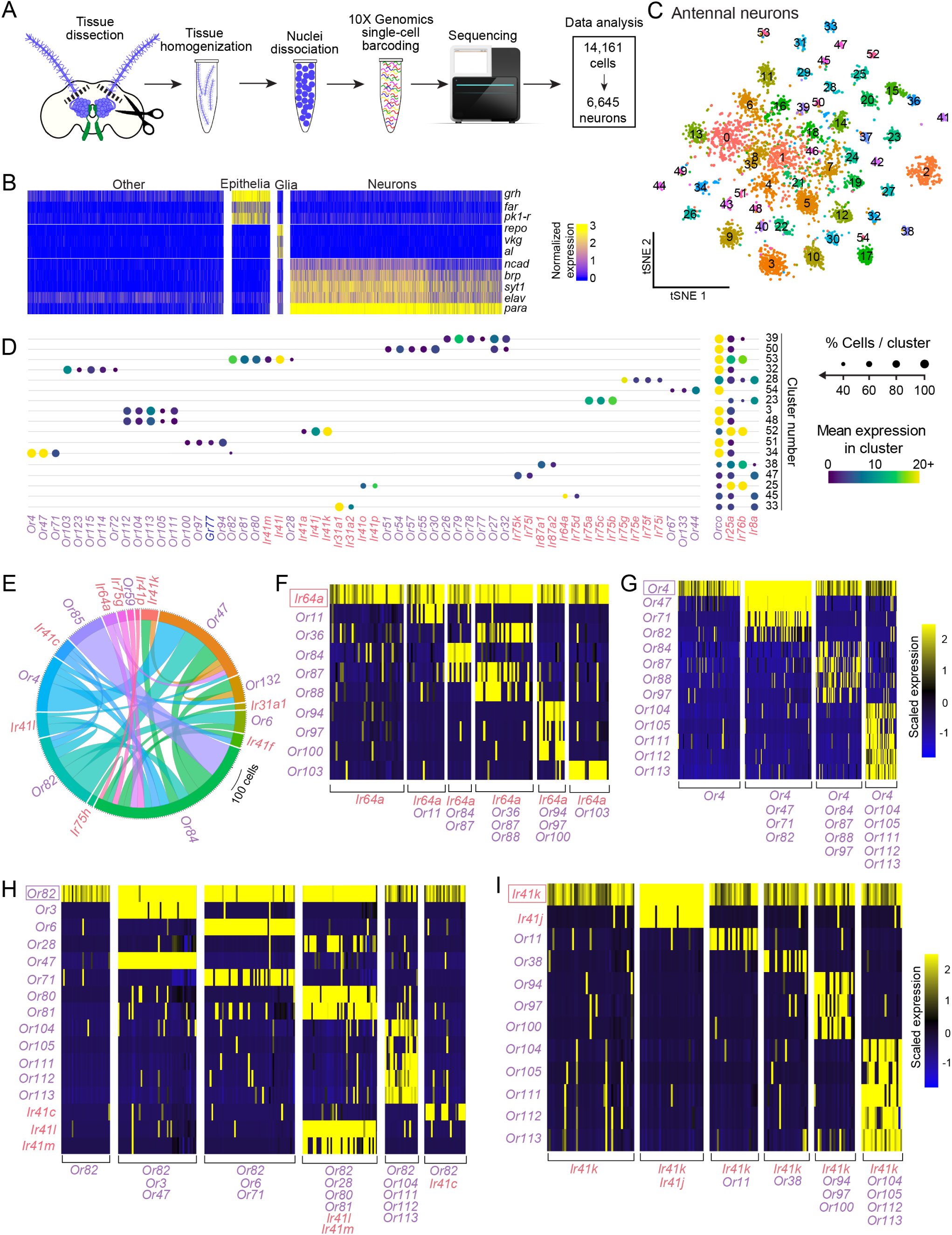
Antennal snRNA-seq reveals complex chemosensory receptor co-expression. (**A**) Schematic of female antenna snRNA-seq workflow. (**B**) Heat map of cells in the antenna grouped according to normalized expression [log(UMI of gene*10,000 / total UMI of cell +1)] of cell type marker. (**C**) t-distributed stochastic neighbor embedding (t-SNE) plot of antennal neurons annotated by cluster (See Figure S9I). (**D**) Dot plot summarizing chemosensory receptor expression in selected clusters (see Figure S10A for full dot plot). Circle size represents % of cells in each cluster that express a given gene above a normalized expression threshold of 1 UMI of gene*10,000/total UMI of cell. Scale indicates mean expression within a cluster. All circles representing a mean expression value greater than 20 have the same color. Circles for clusters with below 35% of cells expressing the indicated chemoreceptor gene are not included in plot (See Figure S10B). (**E**) Chord plot of co-expressed pairs of chemosensory receptors taken from within the 20 highest-expressed ligand-selective receptors. Genes depicted were above a normalized expression threshold of 1 log(UMI of gene*10,000/total UMI of cell +1) and all pairs shown were co-expressed in more than 20 nuclei. (**F-I**) Simplified heatmaps of selected cells, chemosensory receptors, and co-expression patterns using *Ir64a* (F), *Or4* (G), *Or82* (H), or (I) *Ir41k* to select cell types. Scaled expression: Z-score. Receptors are indicated in rows, and cells in columns. Visually-identified cell types are offset with brackets listing the chemosensory receptors expressed in that cell type. See also Figure S11.

To examine the distribution of chemosensory receptors, we averaged expression among cells within an entire cluster and saw cases where multiple receptors were co-expressed (Figure 5D, Figure S10A). Among mean expression levels in the cluster, highly-expressed chemosensory receptors generally belonged to only 1 cluster. Because highly-expressed ligand-selective receptors displayed a strong relationship to individual clusters, we hypothesized that more complex co-expression patterns could be obscured when looking at cluster-level expression patterns. For instance, if a lower-expressed receptor subunit was co-expressed with several different highly expressed receptor subunits, cells exhibiting these combinations might be distributed among several clusters and might not be apparent at this level of analysis. We therefore investigated co-expression within individual cells.

We first looked at the most highly-expressed receptor subunit pairs in a chord plot and saw several co-expression patterns that were not apparent in the cluster-based analyses (Figure 5E). By replotting the expression of individual cells within clusters using heatmaps, we observed many cases of cells co-clustering that expressed discrete combinations of chemosensory receptors. This indicates that clusters are not a faithful representation of all chemosensory receptor combinations with a given cell (Figure S10C). This prompted us to investigate other ways of categorizing cells besides clustering to look more broadly at receptor co-expression patterns.

To analyze the co-expression partners of a given receptor, we filtered our population of neurons for cells that express a receptor gene above a normalized expression threshold of 0.5 log(UMI of gene*10,000 / total UMI of cell +1) (Figure S11A). We then visualized co-expression using heatmaps. Because receptor expression has a bearing on clustering, we performed unsupervised clustering on these cells as a sorting mechanism to group cells by similarity to visualize patterns on heatmaps. Again, cells often grouped into clusters with clearly identifiable receptor expression patterns, including some that contained multiple receptor subunits (Figure S11B). Simplified heatmaps of groups of cells with distinct receptor co-expression patterns are illustrated in Figure 5F-I and S11C-F. This analysis revealed extensive co-expression that points to a far greater variety of cell types than previously anticipated. For instance, *Or82* marks at least 6 different cell types, some that appear to be *Or82*-specific and others that express an additional one, two, four, or five different ligand-selective receptors. Several of these OR-expressing cell types include one or more ligand-selective IR gene. *Ir64a* marks at least 6 different cell types that each expresses one or more ligand-selective OR genes (Figure 5F-I, S11C-F). These results reveal extensive and unexpected chemosensory receptor co-expression in the mosquito antenna.

### Coordinated co-expression of chemosensory receptors in the maxillary palp

We next examined receptor co-expression in the maxillary palp, a smaller and simpler olfactory organ than the antenna that detects important host cues including CO_2_ and 1-octen-3-ol, as well as other host odorants (Grant et al., 1995; Lu et al., 2007; McMeniman et al., 2014; Omer and Gillies, 1971). Each female *Aedes aegypti* maxillary palp contains approximately 35 capitate-peg sensilla that each house three chemosensory neurons (McIver, 1982). Based on prior work that examined the morphology and function of the maxillary palp, the neurons within each sensillum are termed the “A”, “B”, and “C” cells based on their size, from largest to smallest respectively (Figure 6A-C). The A cell responds to the important host cue CO_2_ and houses the Gr3 CO_2_ receptor. The B and C cells both express *Orco*. The B cell is believed to express *Or8*, which detects 1-octen-3-ol, while the C cell expresses *Or49*, which has a less well-defined odorant response profile (Figure 6A).

**Figure 6.**
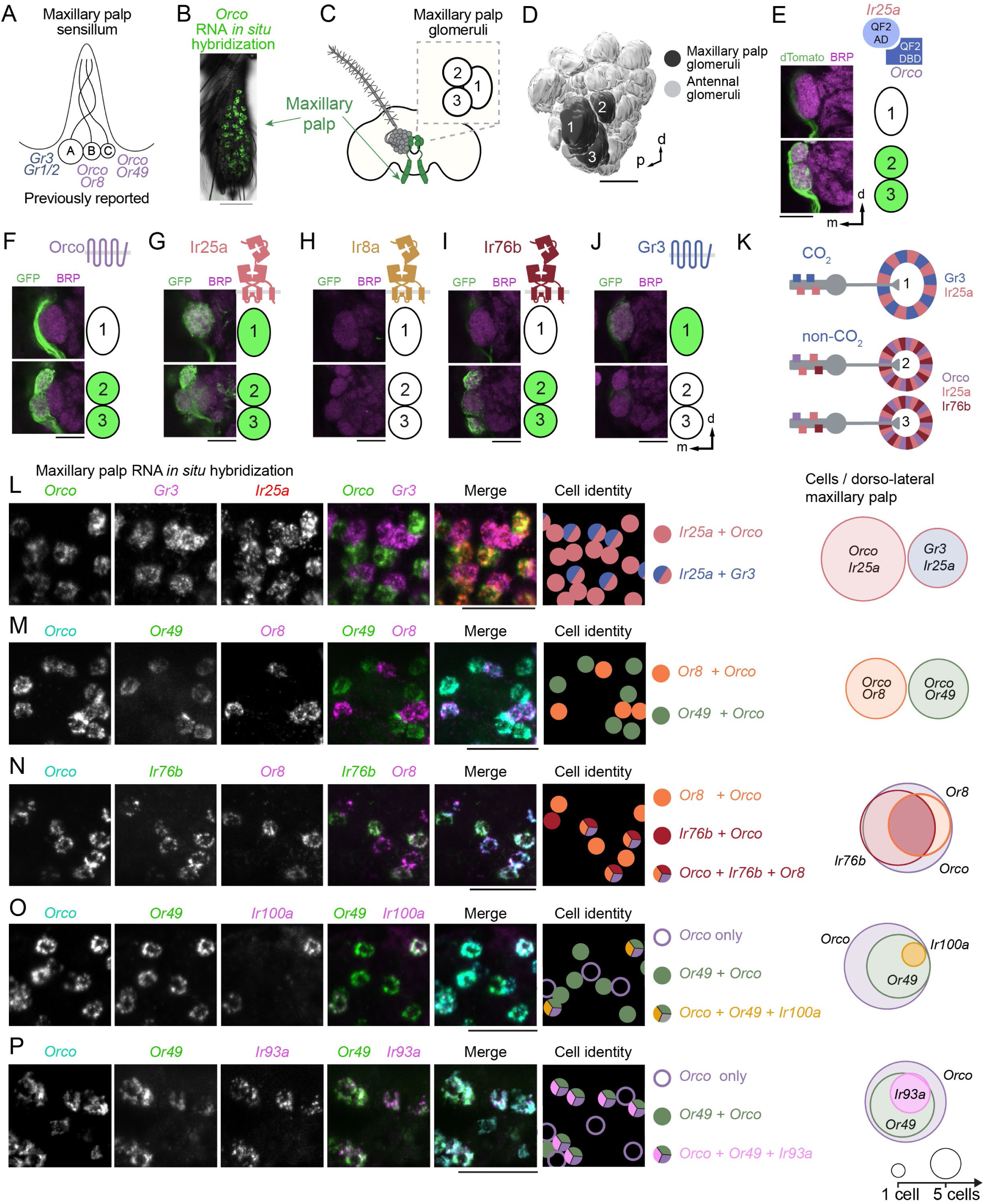
Coordinated co-expression of chemosensory receptors in the maxillary palp. (**A**) Schematic of a single capitate-peg sensillum in the maxillary palp, with A, B, and C cells and the previously reported chemosensory receptor expression. (**B**) Maxillary palp expression of *Orco* in the fourth segment of the maxillary palp revealed by whole-mount RNA *in situ* hybridization. Orientation: proximal up. (**C**) Cartoon indicating maxillary palp (green) and 3 glomeruli that are innervated by the maxillary palp. (**D**) 3-D antennal lobe reconstruction showing 3 glomeruli that are innervated by the maxillary palp. (**E-J**) Single confocal sections through the center of Glomerulus 1 (top) or Glomerulus 2 and Glomerulus 3 (bottom) in left antennal lobes of the indicated genotypes. Sections are taken from Z-stacks presented in Figure 2G (E) and Figure 1I (F-J). (**K**) Schematic of sensory neuron gene expression and glomerular convergence based on (E-J). (**L-P**) Whole-mount maxillary palp RNA *in situ* hybridization with the indicated probes, cartoon schematic indicating cell identity, and quantification of co-expression shown as Euler diagrams, area scaled to mean. n=5 maxillary palps, 26-65 cells/dorso-lateral maxillary palp. See also Figure S12. Scale bars: 50 µm (B), 25 µm (E-J, L-P). Orientation: d=dorsal, m=medial, p=posterior.

We hypothesized that these three cell types project to three glomeruli in the antennal lobe. To delineate the organization of maxillary palp projections in the brain, we used our QF2 and Split-QF2 driver lines to examine maxillary palp sensory innervation of antennal lobe glomeruli (Figure 6C-K). We discovered that Glomerulus 1, which is the largest glomerulus in the antennal lobe (Shankar and McMeniman, 2020), received input from *Gr3-*expressing sensory afferents. Glomerulus 1 was also innervated by *Ir25a*-expressing sensory neurons (Figure 6F-K). Glomerulus 2 and Glomerulus 3 received input from *Orco*-, *Ir25a-*, and *Ir76b*-expressing neurons (Figure 6F-K). Co-expression of *Orco* and *Ir25a* in neurons that project to these two glomeruli was confirmed using the Split-QF2 system. In *Orco-QF2-DBD*, *IR25a-QF2-AD* animals, Glomerulus 2 and Glomerulus 3 were labeled, but Glomerulus 1 was not (Figure 6E). These findings suggest that the A, B, and C cells express multiple co-receptors, spanning IR-OR and IR-GR classes.

To form functional odorant-gated IR or OR complexes, olfactory sensory neurons must express both co-receptors and ligand-selective receptors (Abuin et al., 2011; Benton et al., 2009; Larsson et al., 2004; Neuhaus et al., 2005). To simultaneously monitor the extent of co-expression of both co-receptors and ligand-selective receptors in the A, B, and C cells, we carried out multiplexed whole mount RNA *in situ* hybridization (Choi et al., 2018) in the maxillary palp (Figure 6L-P, Figure S12). The maxillary palp expresses many fewer chemosensory receptor genes than the antenna, with 18 receptors detected at the 1 TPM threshold in the maxillary palp compared to 138 in the antenna at the same threshold (Figure 1F,G), simplifying the task of selecting genes for expression analysis. We performed RNA *in situ* hybridization with probes for 10 of the 18 chemosensory co-receptors and ligand-selective receptors that were present in maxillary palp RNA-seq at a threshold of TPM>1. This technique visualized gene expression with sufficient sensitivity that even *Or71* and *Ir75g* (present at 1.93 and 1.67 TPM, respectively) were readily detected (Figure S12O,P).

We found no overlap in expression of *Orco* and *Gr3* in the maxillary palp, but *Ir25a* was expressed in all *Orco* and all *Gr3* cells (Figure 6L), consistent with our observation that *Ir25a-* expressing neurons project to all three antennal lobe glomeruli. Previous work in *Anopheles gambiae* suggested that *Orco*-expressing neurons in the maxillary palp can be evenly divided into two non-overlapping groups: an *Or8* population and an *Or49* population (Lu et al., 2007). It is widely thought that the same is true in *Aedes aegypti*. We show definitively that *Or8* and *Or49* are expressed in segregated populations of *Orco*-expressing neurons in *Aedes aegypti* (Figure 6M) and, when combined with the results of the previous experiment (Figure 6L), that these cells are also all *Ir25a*-positive. Additional RNA *in situ* hybridization experiments revealed that *Or8*- and *Or49-*expressing cells also often express *Ir76b,* with a bias towards expression in *Or8*-expressing cells (Figure 6N, Figure S12I-O, Data File 1). Taken together these data show that *Orco*-expressing olfactory sensory neurons co-express the co-receptor *Ir25a* and either of the ligand-selective subunits *Or49* or *Or8*, and often co-express the co-receptor *Ir76b* as well.

When we analyzed IR ligand-selective subunit expression, we found that *Ir100a* and *Ir93a* are selectively expressed in a subset of *Or49*-expressing neurons (Figure 6O,P, Figure S12L,M), suggesting that these cells can form functional OR and IR complexes with their respective co-receptors in the same neuron and that co-expression of ORs and IRs may be transcriptionally coordinated. *Or71* and *Or49* were found to be co-expressed, further supporting the idea that multiple ligand-selective ORs can be expressed in an olfactory sensory neuron in *Aedes aegypti* (Figure S12O). We also discovered that the ligand-selective receptor *Ir75g* was expressed in some but not all *Gr3*-expressing cells, which also express *Ir25a* (Figure S12P). Therefore, it is plausible that *Gr3* neurons can functionally express both GRs and IRs.

### Single nucleus RNA sequencing of maxillary palp reveals unanticipated neuronal complexity

As mentioned above, the current view in the field is that the *Aedes aegypti* maxillary palp has a simple organization in which all 35 capitate-peg sensilla are molecularly and functionally identical, each containing the same A, B, and C cells (Figure 6A). Our RNA *in situ* hybridization results called this model into question. To examine receptor co-expression in the maxillary palp in greater detail, we carried out snRNA-seq using similar tissue collection and analysis pipelines used for the antenna (Figure 7A, Figure S13, Figure S14A-C), yielding data from 2,298 cells. Using unsupervised clustering, we categorized these cells into epithelia, muscle, glia, and neurons (Figure 7B-C, Figure S13F). The neuron cluster comprised 630 cells that were further subdivided into four classes that showed remarkable correspondence to cell types previously described in the maxillary palp (Figure 7C-D, Figure S13G-H). Cluster 4 consists of putative mechanosensory neurons marked by expression of *nompC* and *hamlet*. Clusters 1, 2, and 3 were enriched for *Gr3*, *Or8*, or *Or49*, and likely correspond to A, B, and C cells, respectively (Figure 7D,F,J,K, Figure S14D-G).

**Figure 7:**
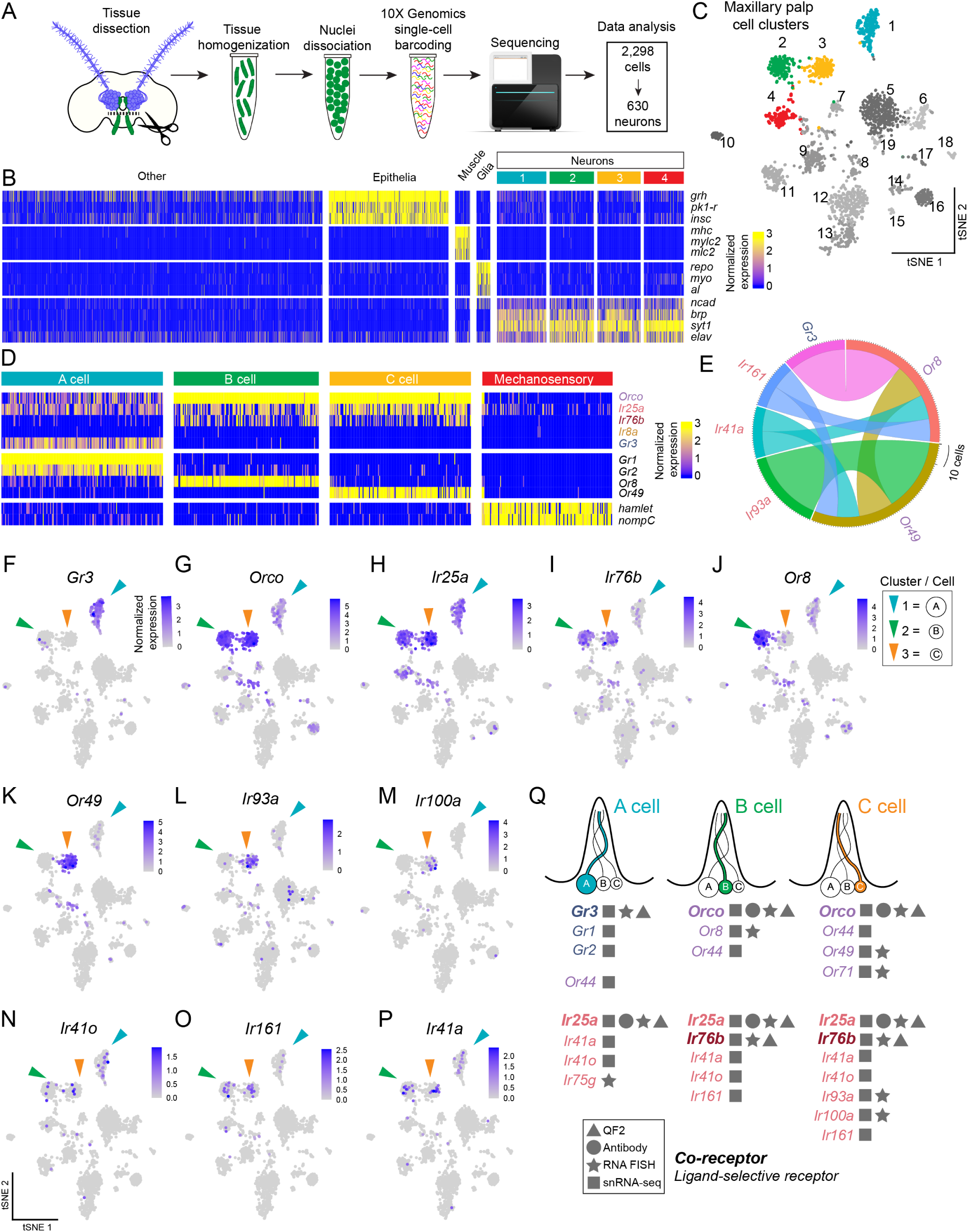
Maxillary palp snRNA-seq reveals unanticipated neuronal complexity. (**A**) Schematic of female maxillary palp snRNA-seq workflow. (**B**) Heat map of nuclei in the maxillary palp grouped according to cell type marker expression. (**C**) tSNE plot of maxillary palp nuclei. (**D**) Heat map of normalized expression of selected genes in 4 identified neuron clusters. See also Figure S13G-H. (**E**) Chord plot of co-expressed pairs of ligand-selective receptors that are present in more than 10 cells. To be considered positively expressed within a cell, gene must meet a normalized expression threshold of 1 log(UMI of gene*10,000/total UMI of cell+1). (**F-P**) Feature plots illustrating normalized expression [log(UMI of gene*10,000 / total UMI of cell +1)] of indicated genes visualized on tSNE plot (see Figure S14). (**Q**) Summary of chemosensory receptor expression in the maxillary palp based on all experimental data in this study (RNA FISH: fluorescent RNA *in situ* hybridization).

To investigate co-expression patterns of receptor genes within these clusters, we generated chord plots and found that *Or49* was co-expressed with *Ir93a* (Figure 7E), confirming our RNA *in situ* hybridization results. Two ligand-selective IR subunits, *Ir41a* and *Ir161*, were co-expressed in both the B cell and C cell. We used feature plots to further visualize the expression of individual receptor subunits within the clusters of maxillary palp neurons. A number of ligand-selective receptor genes were present in discrete clusters (Figure 7F-P, Figure S14H). Confirming our RNA *in situ* hybridization findings, both the *Gr3* cluster and *Orco* clusters also expressed the *Ir25a* co-receptor, and many *Ir76b*-expressing neurons were found in both the B and C cell clusters (Figure 7F-I, Figure S14D-E).

Consistent with RNA *in situ* hybridization data, *Ir93a* and *Ir100a* were expressed in the C cell cluster (Figure 7L-M). *Ir161* was expressed in both B and C cells (Figure 7O-P), and *Or44*, *Ir41a*, and *Ir41o* were found in all three chemosensory clusters (Figure 7N, Figure S14H). We did detect low levels of *Orco* expression in the *Gr3* cluster (Figure 7G) in these snRNA-seq experiments, an observation at odds with results from the three other methods used in the paper and one that we do not currently understand but could be due to differences in the sensitivity of detection methods. We also observed significant variability in expression of ligand-selective and co-receptor subunits across cells (Figure 7F-P), consistent with RNA *in situ* hybridization data (Figure 6L-P). Exploration of the effect of expression level on *in vivo* receptor function in individual neurons will be an interesting direction of future study.

A summary of maxillary palp chemosensory receptor gene expression based on all the data in this study is presented in Figure 7Q. This represents a significant departure from the current view of this sensory system (Figure 6A). Notably, these data suggest that many B and C cells have all the necessary ligand-selective and co-receptor subunits to form both functional OR and IR receptors.

### Receptor co-expression expands the functional responses of olfactory neurons

We next asked whether this extensive chemosensory receptor co-expression would allow maxillary palp neurons to respond to odorants detected by both ORs and IRs. We used single sensillum recording to measure odorant responses of the olfactory sensory neurons housed in maxillary palp-associated capitate-peg sensilla. This method of *in vivo* extracellular electrophysiology enables the simultaneous recordings of A, B, and C cells in response to odorant stimuli in an intact preparation of the female mosquito. Spike sorting is used to discriminate activity of these three neurons based on the amplitude and waveform of their responses. The physical size of the cell positively correlates with the amplitude of the response (Zhang et al., 2019). The largest A cell responds to CO_2_, while the smaller B cell responds to the host odorant 1-octen-3-ol (Bohbot and Dickens, 2009; Cook et al., 2011; Grant et al., 1995; Majeed et al., 2017 ; Syed and Leal, 2007). The smallest C cell has no consistent characteristic ligands in *Aedes aegypti*. We therefore focused our analysis on stimulus-evoked activity of the A cell and B cell while the mosquito was exposed to CO_2_ or odorants likely to stimulate either the OR or IR pathway (Figure 8A-E).

**Figure 8:**
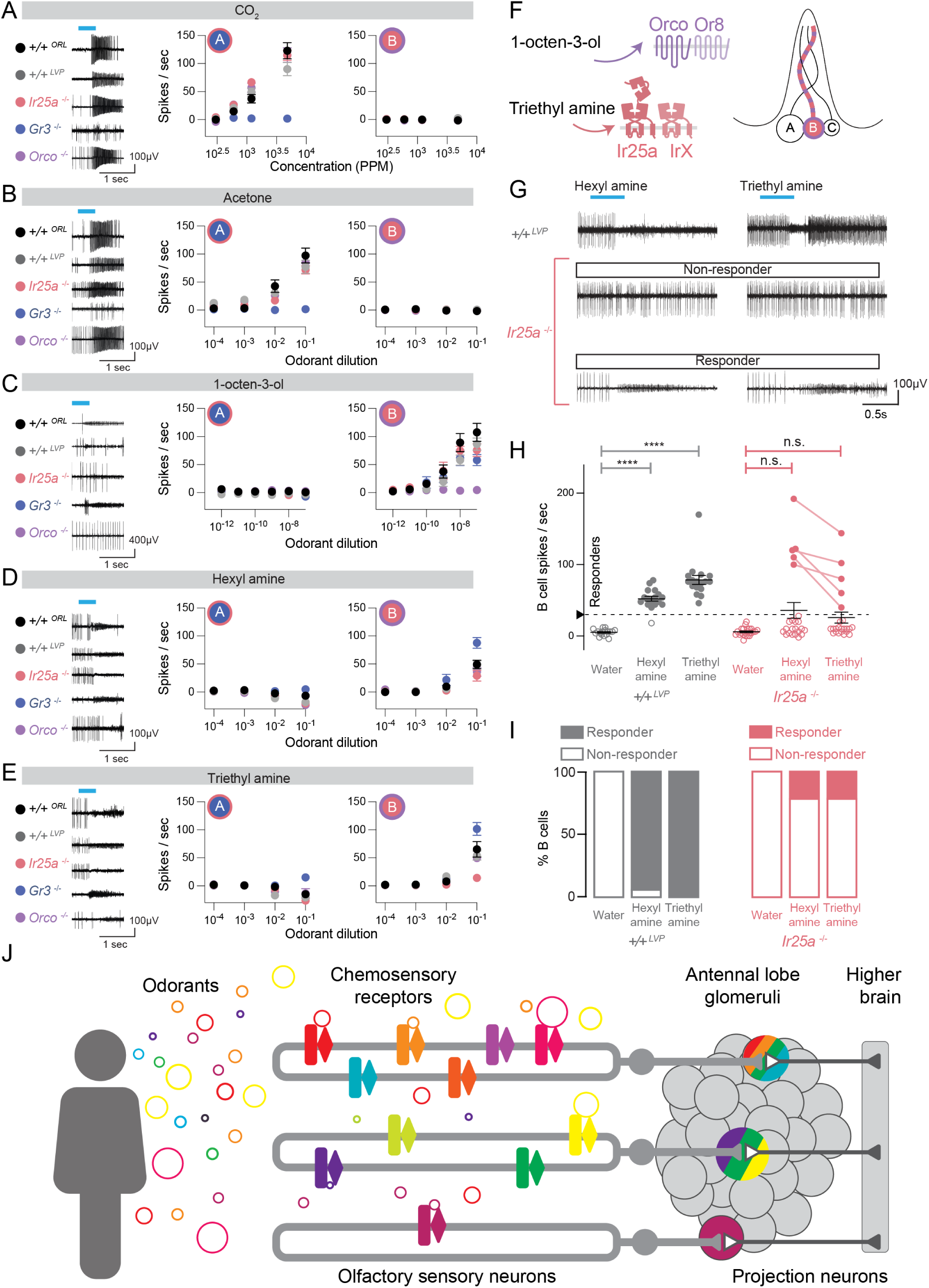
Functional consequences of chemosensory receptor co-expression. (**A-E**) Left, sample traces from maxillary palp single sensillum recordings in each indicated genotype for (A) CO_2_, (B) acetone, (C) 1-octen-3-ol, (D) hexyl amine, and (E) triethyl amine. Stimulus delivery window is indicated by the cyan bar. Middle and right, the number of spikes/sec in the A cell (middle) and B cell (right) for each indicated concentration of the stimulus. Data are presented as mean±SEM, n=4-16 recordings from separate sensilla. (**F**) Schematic of an individual sensillum containing 3 neurons, A, B, and C. Receptor odorant parings for the B neuron are schematized. The identity of the ligand-selective IrX subunit is unknown. (**G**) Sample traces for *+/+^LVP^* (top) and *Ir25a^BamHI/BamHI^*(bottom) with each indicated stimulus. Stimulus delivery window is indicated by the cyan bar. (**H,I**) Quantification of recordings from indicated genotypes shown as dot plots (H) and the stacked bar plots (I) showing the percent of total recordings from each genotype that responded to the stimulus (filled circles) and those that did not (open circles), with 30 spikes/sec defined as response threshold. Data are presented as mean±SEM, n=17 (*+/+^LVP^* ) and n=23 *Ir25a^BamHI/BamHI^* recordings from separate sensilla, n.s., not significant (p=0.1453 for hexyl amine and p=0.1642 for triethyl amine), ****p<0.0001, one-way ANOVA with Kruskal-Wallis test for multiple comparisons. (**J**) A revised model of chemosensory coding in *Aedes aegypti* based on this study.

To determine which family of receptors – GRs, ORs, or IRs – detects a given ligand we recorded odorant responses in wild-type mosquitoes as well as mosquitoes with mutations in *Gr3*, *Orco*, or *Ir25a*. Because the *Gr3* and *Orco* receptor mutants were generated in a different wild-type strain (*+/+^ORL^*) than the *Ir25a* mutant (*+/+^LVP^*) (De Obaldia et al., 2022), all analyses were conducted in two different wild-type background strains. Both wild-type strains showed similar odorant responses in all recordings (Figure 8A-E).

Consistent with previous findings (McMeniman et al., 2014), the A cell responded to CO_2_ in a dose-dependent manner but the B cell did not. The A cell response to CO_2_ was abolished in *Gr3* mutants (Figure 8A). CO_2_-sensing neurons in the maxillary palp respond to multiple odorants in *Aedes*, *Culex*, and *Anopheles* mosquitoes (Lu et al., 2007; Tauxe et al., 2013; Turner et al., 2011) and it has been proposed that *Gr3* is a broadly-tuned receptor that responds to many odorants. We examined the response to a recently identified CO_2_-neuron activator, acetone (Ghaninia et al., 2019), which like CO_2_ also activated the A cell but not the B cell (Figure 8B). The response to acetone was abolished in the *Gr3* mutant (Figure 8B), which suggests that the CO_2_ receptor can interact with non-CO_2_ ligands.

The host-emitted odorant 1-octen-3-ol has been shown to activate the B cell. Both *Aedes aegypti* and *Anopheles gambiae* Or8-Orco, which are expressed in the B cell, respond to 1-octen-3-ol when expressed in heterologous cells (Bohbot and Dickens, 2009; Lu et al., 2007). We found that firing of the B-cell, but not the A cell, increased in the presence of 1-octen-3-ol (Figure 8C). The B cell response to 1-octen-3-ol was abolished in the *Orco* mutant, but not in *Gr3* or *Ir25a* mutants (Figure 8C), consistent with the role of Or8-Orco in detecting this compound.

Volatile amines, including polyamines, have been proposed to be IR ligands in *Drosophila melanogaster* (Geier et al., 1999; Hussain et al., 2016; Min et al., 2013; Silbering et al., 2011). We therefore examined the response of maxillary palp neurons to two volatile amines, hexyl amine and triethyl amine. We found that both amines activated the B cell in wild type, *Gr3* mutants, and *Orco* mutants (Figure 8D-F). Average responses to hexyl amine and triethyl amine were strongly reduced but not abolished in the *Ir25a* mutant. When we scrutinized the raw data carefully, we noted that responses to both stimuli in *Ir25a* mutants fell into two clear types of neurons. The majority of *Ir25a* mutant neurons did not respond to these stimuli at all, but a few responded even more robustly than wild type (Data File 1).

To determine if there are two different functional types of B neurons, we generated a second independent dataset using these stimuli to examine responses in an additional 17 wild-type (*+/+^LVP^*) neurons and 23 additional *Ir25a* mutant neurons. The response to the water control stimulus never exceeded 30 spikes/sec firing frequency in either genotype, and we used this as a threshold to classify neurons as “responders” or “non-responders” (Figure 8H-I). We found that all *+/+^LVP^* neurons responded to triethyl amine, and 16 out of 17 *+/+^LVP^* neurons responded to hexyl amine (Figure 8H-I). Responses to both stimuli were significantly higher than the water control in wild-type (*+/+^LVP^*) neurons. In contrast, most neurons in the *Ir25a* mutant did not respond to either triethyl amine or hexyl amine (78.3% n = 23) and neither stimulus elicited significantly different responses from the water control when taking the entire population of 23 recorded neurons into account.

We noted that 5 out of 23 neurons (21.7%) showed strong responses to both amines that exceeded the corresponding response in*+/+^LVP^* neurons (Figure 8G-I). These neurons were considered outliers by a ROUT analysis (Q=1%), consistent with the classification system that we used to categorize neurons as responders or non-responders. Given our discovery of multiple additional IRs and ORs co-expressed along with *Or8* in the B cells (Figure 7Q), we speculate that there are at least two distinct types of B neurons, one that requires *Ir25a* to respond to amines and the other that does not. We hypothesize that this second type of B neuron expresses the *Ir76b* co-receptor, which could form functional amine receptors with one or more of the ligand-selective IRs expressed in the B cell. Our findings that the B cell responds to 1-octen-3-ol in an *Orco*-dependent manner and to triethyl amine and hexyl amine in an *Ir25a*-dependent manner is consistent with the hypothesis that ORs and IRs are functionally co-expressed in the same neurons and that co-expression enables these cells to respond to ligands that activate both classes of receptors.

## DISCUSSION

### Combinatorial chemosensory receptor co-expression in *Aedes aegypti*

The mismatch between the number of receptors in the *Aedes aegypti* genome and the number of glomeruli in the antennal lobe is resolved in part by co-expression of multiple odorant receptors in individual olfactory sensory neurons. We found that co-expression is extremely widespread, both between and within OR and IR receptor families, and that the number of receptors expressed in a neuron can vary substantially. While some neurons express only an individual co-receptor and ligand-selective receptor pair, others express “sets” of frequently co-expressed receptor subunits. We were surprised to find that a single receptor subunit could be co-expressed with completely different combinations of receptor subunits. The biological significance of this finding remains to be seen and the exact number of receptor groupings will require extensive additional study.

We found that many commonly co-expressed IRs and ORs belong to mosquito gene family expansions. The *Ir41* clade of IRs was among the most common IRs to be co-expressed with ORs. This clade is greatly expanded in *Aedes aegypti* relative to *Drosophila melanogaster*, with 16 members compared to only 3 orthologous genes, respectively (Matthews et al., 2018). The *Drosophila melanogaster* orthologues, *Ir41a*, *Ir76a*, and *Ir92a*, have all been shown to compose channels that respond to amines (Hussain et al., 2016; Min et al., 2013; Silbering et al., 2011). Volatile amines are enriched in human odor and are known to play an important role in the detection of humans by mosquitoes (Bernier et al., 2000; de Lacy Costello et al., 2014; De Obaldia et al., 2022). It is tempting to speculate that the expansion of the *Ir41* clade in *Aedes aegypti* enhances the ability of these mosquitoes to detect amines present in human odor, although this remains to be tested. Similarly, many of the commonly co-expressed ORs are also members of gene expansions in *Aedes aegypti* as well as *Anopheles gambiae* but have no direct orthologues in *Drosophila melanogaster* (Matthews et al., 2018). Because these mosquito olfactory specialists maintain a very different diet than *Drosophila melanogaster*, it will be fascinating to examine if the ligands for these receptors are enriched in human body odor.

Among the OR subunits found to be co-expressed with other ORs, *Or4* was notable. Subspecies of *Aedes aegypti, Aedes aegypti aegypti* and *Aedes aegypti formosus*, have evolved different host preferences, either for humans or non-human mammals respectively. *Or4* responds to the host odorant sulcatone and variation in its coding sequence corresponds strongly with human host preference (McBride et al., 2014; Rose et al., 2020). We observed co-expression of *Or4* in at least three molecularly distinct groups of sensory neurons that each contain 3, 4, or 5 non-overlapping ligand-selective OR subunits in addition to *Or4*. The ligand response profiles of these receptors, which all belong to the families of ORs that are expanded in *Aedes aegypti*, are unknown but we hypothesize that they respond to additional human host odorants. Our experiments used strains of mosquitoes that belong to the *Aedes aegypti aegypti* subspecies and show a preference for human over non-human animal odor (McBride et al., 2014; Rose et al., 2020). It will be interesting to see if co-expression of these OR subunits with *Or4* is altered in *Aedes aegypti formosus*, which prefer non-human animals.

### The possibility of neuronal co-convergence in antennal lobe glomeruli

The mismatch between the number of chemosensory receptors in the genome and the number of glomeruli in the antennal lobe originally pointed to two simple models: co-expression or co-convergence. In this study we presented extensive evidence for widespread co-expression in the *Aedes aegypti* olfactory system. However, our findings also point to the likelihood that co-convergence exists in this olfactory system as well. Previous work in *Drosophila melanogaster* and *Mus musculus* has shown that the identity of an olfactory neuron is defined by the chemosensory receptor it expresses. However, our snRNA-seq results identify a wide variety of cell types that cannot be defined by expression of a single chemosensory receptor in a given neuron. Rather *Aedes aegypti* chemosensory cell types are defined by the entire complement of odorant receptors they express. For example, we identified 7 types of *Ir64a* neurons and 4 types of *Or4* neurons each co-expressed with many different types of ORs and IRs. Given this revised view of olfactory sensory neuron types, the number of cell types in the antenna far exceeds the number of glomeruli in the antennal lobe. The most likely solution to this problem is co-convergence of neurons in the antennal lobe, although this remains to be tested directly.

What rules govern co-convergence in the mosquito? Do neurons that express the same dominant ligand-selective receptor subunit but also co-express different combinations of ligand-selective receptors project to the same glomerulus or separate glomeruli? These contrasting organizational principles would result in very different models of odor coding. We demonstrate that *Aedes aegypti* mosquitoes express many receptors in some neurons but only a single receptor in others. The presence of multiple receptors in a given neuron could serve as a mechanism to integrate odorant information in the primary sensory neuron itself rather than at the first synapse in the antennal lobe. Co-convergence of olfactory sensory neurons onto the antennal lobe could allow for the early integration of olfactory cues while still retaining discrete input channels that could be selectively modulated during changes in behavioral state, such as the suppression of host-seeking after a blood meal. Exploring the organization of this sensory system and the downstream circuitry will be essential, including the question how projection neurons encode olfactory information given such extreme diversity in sensory afferent types.

### Coordinated co-expression between *IR*, *OR*, and *GR* ligand-selective receptors

We identified co-expression of co-receptors and ligand-selective receptors belonging to distinct chemosensory families in single neurons in both the antenna and the maxillary palp. This co-expression poses a gene regulatory problem for an olfactory neuron. For ORs and IRs to form functional chemosensory receptors, at least one co-receptor and one ligand-selective receptor must be expressed in a cell. We have demonstrated that multiple ORs and IRs are expressed in specific receptor “sets”. Thus, the transcriptional landscape in *Aedes aegypti* olfactory neurons is not only permissive to co-expression, but ensures certain receptors are expressed with others.

How might this complex code of chemosensory receptor co-expression be regulated? In vertebrates, an elaborate epigenetic silencing mechanism ensures that each olfactory neuron stochastically expresses only a single allele of a single odorant receptor (Bashkirova and Lomvardas, 2019). In contrast, *Drosophila melanogaster* is thought to use a more conventional transcription factor code in which the specification of a neuron and the expression of its chemosensory receptor is tightly regulated (Jafari and Alenius, 2015; Li et al., 2016; Ray et al., 2008). Single-cell sequencing data generated from developing *Drosophila melanogaster* olfactory neurons demonstrate a complex regulatory landscape in which a set of transcription factors govern receptor expression, axon targeting, or both (Li et al., 2020). Two recent studies documented extensive co-expression of co-receptors in *Drosophila melanogaster*, calling into question the rules that regulate olfactory organization in this insect (McLaughlin et al., 2021; Task et al., 2021). It is yet to be determined if mosquito orthologues of these transcription factors have been co-opted to regulate the co-expression we observe or if this novel olfactory organization demands a distinct transcriptional mechanism.

There are other examples of receptor co-expression within a single class of chemosensory receptor. Polycistronic expression of multiple odorant receptors in *Anopheles gambiae* sensory neurons has been reported (Karner et al., 2015). This differs from co-expressed receptors in *Aedes aegypti*, which are often not closely associated genes within the genome and suggests that other mechanisms of gene regulation must account for the co-expression we observe. A recent study also identified neurons in *Anopheles gambiae* that co-express *Orco* and *Ir76b*, but the extent of OR-IR co-expression in this species remains to be seen (Ye et al., 2021). In *Drosophila melanogaster* there are rare cases of OR-OR or OR-IR co-expression, which have been thought the exceptions rather than the rule. *Or35a* is co-expressed with *Ir76b* (Silbering et al., 2011), and while these neurons respond to many odorants (Silbering et al., 2011; Yao et al., 2005), the functional role of co-expression remains unknown. In *Drosophila melanogaster*, *Or49a* and *Or85f* are co-expressed in a specific olfactory sensory neuron population where they play redundant roles in predator avoidance (Ebrahim et al., 2015). Recent work suggests that there may be more examples to be discovered (McLaughlin et al., 2021; Task et al., 2021).

Our findings are reminiscent of the nematode *Caenorhabditis elegans*, which copes with a very large number of chemosensory receptor genes and a very small number of sensory neurons by extensive receptor co-expression (Troemel et al., 1995; Vidal et al., 2018). *Aedes aegypti* mosquitoes have many more olfactory sensory neurons than *Caenorhabditis elegans* nematodes and the circuit organization also differs dramatically between the two. Another example of extensive olfactory receptor co-expression is seen in mice, where a subpopulation of olfactory sensory neurons each express multiple MS4a chemosensory receptors and project to the so-called necklace glomeruli that surround the main olfactory bulb (Greer et al., 2016). Interestingly these neurons respond to a number of cues that regulate innate behaviors, such as food odors and pheromones. Perhaps chemosensory receptor co-expression is more conducive to sensory systems that drive innate rather than learned behaviors.

### Maxillary palp chemosensory neurons go beyond a simple A, B, and C organization

The maxillary palp is a multi-modal sensory organ that responds to CO_2_ (Acree et al., 1968; Gillies, 1980; Grant et al., 1995), temperature (Roth, 1951), mechanical stimuli (Bohbot et al., 2014), attractive monomolecular odorants such as 1-octen-3-ol (Syed and Leal, 2007; Takken and Kline, 1989; Vythilingam et al., 1992), as well as blends of odorants extracted from human hosts (Tauxe et al., 2013). The prior view of the organization of the maxillary palp is that all volatile odorant-detecting capitate-peg sensilla house the sensory dendrites of three neurons that form identical repeating units: one large CO_2_-sensitive neuron that expresses *Gr3* (A cell), and two smaller neurons that express either Or8-*Orco* (B cell) or Or49-*Orco* (C cell) (Lu et al., 2007; McIver, 1972). It is difficult to reconcile the functional diversity of responses with this simple cellular organization. We demonstrate through multiplexed RNA *in situ* hybridization and snRNA-seq that the receptor composition of these neurons is far more complex, and they can be subdivided into many more than three cell types. Consistent with this idea, we found that B cells can be separated into different types based on their physiological response to volatile amines. This is revealed in *Ir25a* mutant animals, where the response to triethyl amine and hexyl amine is abolished in most B cells, but a subset of neurons retains their responses to this compound. We found that *Ir76b* is expressed in a subset of the *Or8*-expressing B cells, as are *Ir161*, *Ir41a*, and *Ir41o*. It is possible that these IRs mediate amine responses in the subset of *Ir25a* mutant neurons that retain amine responses. It will be interesting to examine this possibility and to explore the heterogeneity of odorant responses across all maxillary palp neuron types.

### Receptor co-expression as a possible mechanism for robust mosquito attraction to humans

We hypothesize that receptor co-expression is used broadly to detect redundant cues that are present in human odor, a blend that can vary from individual to individual and contains hundreds of different chemicals (Bernier et al., 1999; Bernier et al., 2000; De Obaldia et al., 2022). Both volatile amines and 1-octen-3-ol are emitted from human skin (Bernier et al., 2000; Cork and Park, 1996 ; de Lacy Costello et al., 2014). It is possible that receptor co-expression is used to form a highly redundant detection system for different cues that represent the same ecological target: humans. This motif has the benefit of limiting the number of neurons needed to detect varied odorants with the same meaning. However, in exchange it may sacrifice the ability to distinguish between cues detected by receptors expressed in the same sensory neurons. In summary, our study reveals unexpected complexity in the gene expression and functional organization of the mosquito olfactory system that may explain the persistence of mosquitoes in hunting humans. Future attempts to refine the design of repellents to ward off mosquitoes or attractant traps to lure them will have to have to reckon with the complexity of this system.

## ACKNOWLEDGMENTS

We thank Emily Dennis, Laura Duvall, Itzel Ishida, Philip Kidd, Erica Korb, Carolyn McBride, Christopher Potter, Darya Task, Zhilei Zhao, and members of the Vosshall Lab for comments on the manuscript; Gloria Gordon and Libby Mejia for expert mosquito rearing at Rockefeller and Mengistu Dawit Bulo for rearing strains at SLU; Javier Marquina-Solis for assistance with antennal lobe tracing; Priyanka Lakhiani and Julia Deere for participation in troubleshooting mosquito nuclei extraction; Christina Pyrgaki, Carlos Rico, Katarzyna Cialowicz, and Alison North at the Rockefeller Bio-Imaging Resource Center (RRID:SCR_017791) for assistance with confocal imaging; Helen Duan, Bin Zhang, and Connie Zhao at the Rockefeller Genomics Core for quality control for snRNA-seq samples and 10X Genomics sequencing; Daniel Gross, James Petrillo, and Peer Strogies at the Rockefeller Precision Instrumental Technologies (PIT) Resource Center for advice and fabrication of custom equipment; Olena Riabinina and Christopher Potter for advice and for providing Q-system reagents prior to publication; Carolyn McBride, Matthew DeGennaro, and members of the *Aedes* Toolkit Group for advice and discussion; Caroline Jiang for advice on statistical analysis; Nipun Basrur, Priya Rajasethupathy, Andrea Terceros, Harry Choi, and Molecular Instruments for advice on RNA *in situ* hybridization experiments; Andrea Terceros, Andras Sziraki, Junyue Cao, and 10X Genomics for advice on mosquito nuclei extraction and analysis; Rob A. Harrell II at the Insect Transgenesis Facility at the University of Maryland for embryo injections; Raphael Cohn, Gaby Maimon, Cory Root, Vanessa Ruta, and Ari Zolin for useful discussions; and Frances Weis-Garcia and the members of the MSKCC Antibody and Bioresource Core Facility for preparation of the nc82/Brp monoclonal antibody.

## AUTHOR CONTRIBUTIONS

M.A.Y. carried out all central tissue immunostaining. M.H. carried out all peripheral tissue immunostaining and RNA *in situ* hybridization experiments. B.J.M. provided chemosensory gene and transcript analysis. Z.G. cloned and isolated Split-QF2 lines with M.H. Z.N.G. cloned and isolated QF2 stop codon replacement lines with B.J.M. and M.A.Y. S.R. worked with M.H. to generate RNA *in situ* hybridization data. O.V.G. collected all tissue for snRNA-seq together with M.H. and M.A.Y. O.V.G. processed tissue for snRNA-seq experiments at Rockefeller. At Baylor, Y.Q. carried out sample preparation, flow cytometry, and 10X Genomics library preparation. T.-C.L. carried out snRNA-seq data analysis including read alignment, quality checking, and cell and gene filtering. O.V.G. carried out additional downstream analysis. T.-C.L. together with O.V.G. generated the figure panels for Figure 5 and Figure 7. H.L. supervised Y.Q and T.-C.L. and oversaw snRNA-seq experimental design and data analysis. The single sensillum recordings in Figure 8A-E were carried out by M.G. and those in Figure 8F-I were carried out by G.C.-V. R.I. supervised M.G. and G.C.-V. and analyzed all of the data in Figure 8 together with M.G. and G.C.-V. M.A.Y., M.H., and L.B.V. together conceived the study, designed the figures, and wrote the paper with input from all authors.

## Materials and Methods

### Human and animal ethics statement

Blood-feeding procedures and behavioral experiments with live hosts were approved and monitored by The Rockefeller University Institutional Animal Care and Use Committee (IACUC protocol 17018) and Institutional Review Board (IRB protocol LV-0652), respectively. Human volunteers gave their written informed consent to participate.

### Mosquito rearing and maintenance

*Aedes aegypti* wild-type laboratory strains (Liverpool and Orlando), CRISPR-Cas9 knock-in, and piggyBAC *QUAS* transgenic strains were maintained and reared at 25 – 28°C, 70-80% relative humidity with a photoperiod of 14 hr light: 10 hr dark as previously described (DeGennaro et al., 2013). Adult mosquitoes were provided constant access to 10% sucrose. For routine strain maintenance, animals were primarily blood-fed on live mice and occasionally on live human volunteers. Newly generated strains were blood-fed on human volunteers until they were established. All experiments except those in Figure 2J-T were conducted on adult female mosquitoes. Detailed genotypes used in each figure can be found in Data File 1.

### Generation of chemosensory receptor *QF2* and Split-*QF2* knock-in strains

*T2A*-*QF2* gene-sparing stop codon replacement lines were generated using the strategy outlined in Matthews et al. (Matthews et al., 2019). sgRNAs were placed as close to the stop codon as possible and donor constructs were designed to remove the stop codon and replace it with an in-frame cassette containing the *T2A* ribosomal skipping sequence and the *QF2* transcription factor or Split-*QF2* domains, comprising the *QF2* activation domain *QF2-AD*, or the *QF2* DNA-binding domain *QF2-DBD*. This strategy spares the function of the gene at the locus being targeted, expresses *QF2* or Split-*QF2* domains in the cells specified by enhancers at the locus. Insertions were marked by the *3xP3* enhancer expressing a fluorescent protein. To identify effective sgRNAs, 5 candidate sgRNAs per gene were first injected into separate pools of 500 Liverpool embryos and CRISPR-Cas9-mediated cut rate was evaluated as previously described (Kistler et al., 2015). Either a single sgRNA or 2 sgRNAs with the highest cut rates were then chosen to be injected with donor plasmids to target chemosensory gene loci using homology-directed repair. sgRNAs targeted the respective gene near the stop codon, target sequence with protospacer adjacent motif (PAM) underlined:

*Ir25a*: GTTTGTGTGCGTGTCCGTA <u>TGG</u>

*Ir76b*: GTATTACACTTATCTAAATA <u>TGG</u>

*Ir8a*: GTCACGCTTGTTGTACAGGG <u>CGG</u>, GAACAATTTGAACAAGGTCG <u>TGG</u>

*Gr3*: GTTAGTGATGCATAATATGA <u>CGG</u>

*Orco*: GTCACCTACTTCATGGTGT <u>TGG</u>

sgRNA DNA template was prepared by annealing oligonucleotides as described (Kistler et al., 2015). *In vitro* transcription was performed using HiScribe Quick T7 kit (NEB E2050S) following the manufacturer’s directions. Following transcription and DNAse treatment for 15 min at 37°C, sgRNA was purified using RNAse-free SPRI beads (Ampure RNAclean, Beckman-Coulter A63987), and eluted in Ultrapure water (Invitrogen, 10977–015).

Donor plasmids were constructed by Gibson assembly using the following fragments for QF2 lines:

1. *pUC19* digested with XbaI and BamHI
2. Left and right homology arms: *Gr3* (left: 1.9 kb, right: 1.6 kb), *Ir25a* (left: 1.8 kb, right: 1.6 kb), *Ir76b* (left: 1.2 kb, right: 2.2 kb), *Ir8a* (left: 1.7 kb, right: 1.7 kb), *Orco* (left: 1.2 kb, right: 1.3 kb) generated by polymerase chain reaction (PCR) using Liverpool genomic DNA as a template
3. A 2.6 kb fragment containing *T2A-QF2-SV40, 3xP3-dsRed*, PCR-amplified from a previously assembled vector (*ppk10779-T2A-QF2-SV40, 3xP3-dsRed*, Addgene accession #130667)

For Split-QF2 lines, donor plasmids were constructed by generating fragments using PCR from the indicated template with indicated primers in Data File 1 and assembled using NEBuilder HiFi DNA Assembly (NEB E5520S):

*Ir25a-T2A-QFAD::Zip+-SV40-3xP3-eYFP-SV40* was composed of:

1. Plasmid backbone with *Ir25* homology arms from *Ir25a-T2A-QF2* plasmid (6 kb)
2. *T2A-QFAD::Zip+-SV40* sequence from (Riabinina et al., 2019), fragment synthesized by Genewiz, sequence in Data File 1 (1.5 kb)
3. *3xP3-EYFP-SV40* from *pDSAY* (Addgene, #62291) (1.2 kb)

*Orco-T2A-Zip−::QFDBD-SV40-3xP3-dsRED-SV40* was composed of:

1. Plasmid backbone with *Orco* homology arms and *3xP3-dsRED-SV40* from *Orco-T2A-QF2* plasmid (6.3 kb)
2. *T2A-Zip−::QFDBD-SV40* synthesized by Genewiz, sequence in Data File 1 (1.5 kb)

For all *QF2* and Split-*QF2* constructs, the stop codon of the endogenous gene was removed and the PAM sequences corresponding to the sgRNAs used for injection were modified by PCR mutagenesis during Gibson assembly by introducing synonymous codon substitutions to protect the sequence from Cas9 cleavage while retaining the amino acid identity. Plasmids were isolated using an endotoxin-free plasmid midiprep kit (Macherey-Nagel) for QF2 lines and NucleoBond Xtra Midi Endotoxin-Free plasmid kit (Clontech 740420.50) for Split-QF2 lines and eluted in ultrapure water prior to injection. Donor plasmids are available at Addgene (accession numbers #162520-162526). Approximately 2,000 wild-type Liverpool strain *Aedes aegypti* embryos were injected with a mix containing recombinant Cas9 protein (PNA Bio, CP01) at 300 ng/µL, sgRNAs at 40 ng/µL and donor DNA plasmid (300 ng/µL for QF2 lines, 600 ng/µL for Split-QF2 lines) at the Insect Transformation Facility at the University of Maryland Institute for Bioscience & Biotechnology Research. Embryos were hatched and surviving G0 males and females were crossed to wild-type Liverpool mosquitoes and their G1 offspring were screened for fluorescence indicating positive stable germ line transformants. For QF2 lines, the fidelity of insertion was verified by PCR and Sanger sequencing. One representative line for each chemosensory receptor QF2 knock-in was selected for further study. QF2-driven expression patterns were examined by crossing to *QUAS-CD8:GFP-3xP3-ECFP* and/or *QUAS-dTomato-T2A-GCaMP6s-3xP3-ECFP*.

A technical problem arose in the construction of the *QUAS-dTomato-T2A-GCaMP6s-3xP3-ECFP* plasmid that caused only a single copy of dTomato to be introduced into the mosquito, rather than the brighter tandem dTomato or tdTomato that is more conventionally used. Nevertheless we found that dTomato is sufficiently bright for our experiments (Shaner et al., 2004).

All lines were outcrossed to wild-type Liverpool mosquitoes for at least 3 generations prior to being used in experiments. For Split-QF2 lines, a single family with the correct insertion was confirmed by PCR and Sanger sequencing for *Ir25a-QF2-AD* and *Orco-QF2-DBD*. To propagate these lines, a male founder was chosen to cross to wild-type Liverpool females. Animals were then back-crossed to Liverpool for at least 2 additional generations. To evaluate if the Split-QF2 system was functional in *Aedes aegypti*, *Ir25a-QF2-AD was crossed to QUAS-dTomato-T2A-GCaMP6s.* The resulting *Ir25a-QF2-AD, QUAS-dTomato-T2A-GCaMP6s* animals were then crossed to *Orco-QF2-DBD*. Expression of the dTomato reporter was observed in larval antennae and subsequently confirmed in adult antennae and brains.

### *QUAS* transgenic strains

*QUAS-CD8:GFP-3xP3-ECFP* and *QUAS-dTomato-T2A-GCaMP6s-3xP3-ECFP* transgenic strains were described previously (Matthews et al., 2019). Two independent insertions of the *QUAS-dTomato-T2A-GCaMP6s-3xP3-ECFP* reporter line (Jové et al., 2020; Matthews et al., 2019) were used in this study. These are located on different chromosomes and were used according to the crossing scheme needed for a given experiment. See Data File 1 for details.

### Chemosensory receptor mutant strains

The three chemosensory receptors mutant strains used in this study was previously described: *Ir25a^BamHI/BamHI^* (De Obaldia et al., 2022), *Gr3^4/4^* (McMeniman et al., 2014), *Orco^16/16^* (DeGennaro et al., 2013). *Gr3^4/4^* and *Orco^16/16^* were generated in the *Orlando* background (here referred to as *+/+^ORL^*) and the *Ir25a^BamHI/BamHI^* mutant was generated in the *Liverpool* background (here referred to as *+/+^LVP^*). To account for possible difference in genetic background the *+/+^ORL^* strain was used as the controls in all experiments where the *Gr3^4/4^* and *Orco^16/16^* mutants were used, and the *+/+^LVP^* strain was used as the control in experiments where the *Ir25a^BamHI/BamHI^*mutant was used.

### Transcript abundance estimates of *Aedes aegypti* OR, IR, and GR genes

Expression values for adult sugar-fed, non-blood-fed female sensory tissues were retrieved from the *Aedes aegypti* L5 genome GitHub repository (https://github.com/VosshallLab/AGWG-AaegL5) at this link: https://github.com/VosshallLab/AGWG-AaegL5/raw/master/AGWG%20AaegL5%20Chemoreceptor%20TPM.xlsx. These expression values reflect libraries from a previous transcriptome study (Matthews et al., 2016) that had been aligned to the *Aedes aegypti* genome (AaegL5) and chemosensory receptor geneset annotation reported in units of Transcripts Per Million (TPM) (Matthews et al., 2018). The number of genes from each of three gene families (ORs, IRs, and GRs) with expression values above the indicated threshold were plotted in Figure 1F,G and are available in Data File 1.

### Whole brain fixation and immunostaining

Dissection of adult brains and immunostaining was done as previously described (Matthews et al., 2019). 6-14 day-old mosquitoes were anesthetized on wet ice. Heads were carefully removed from the body by pinching at the neck with sharp forceps. Heads were placed in a 1.5 mL tube for fixation with 4% paraformaldehyde, 0.1 M Millonig’s Phosphate Buffer (pH 7.4), 0.25% Triton X-100, and nutated for 3 hr. Brains were then dissected out of the head capsule in ice-cold Ca^+2^-, Mg^+2^-free phosphate buffered saline (PBS, Lonza 17-517Q) and transferred to a 24-well plate. All subsequent steps were done on a low-speed orbital shaker. Brains were washed in PBS containing 0.25% Triton X-100 (PBT) at room temperature 6 times for 15 min. Brains were permeabilized with PBS, 4% Triton X-100, 2% normal goat serum (Jackson ImmunoResearch #005-000-121) for ∼48 hr (2 nights) at 4°C. Brains were rinsed once and then washed with PBT at room temperature 6 times for 15 min. Primary antibodies were diluted in PBS, 0.25% Triton X-100, 2% normal goat serum for ∼48 hr (2 nights) at 4°C. Brains were rinsed once then washed in PBT at room temperature 6 times for 15 min. Secondary antibodies were diluted in PBS, 0.25% Triton X-100, 2% normal goat serum for ∼48 hr (2 nights) at 4°C. Brains were rinsed once then washed in PBT at room temperature 6 times for 15 min. Brains were equilibrated overnight in Vectashield (Vector Laboratories H-1000) and were mounted in Vectashield. The following primary antibodies were used: anti-Brp/nc82 (mouse; 1:50, Developmental Studies Hybridoma Bank – see below) and/or anti-GFP (rabbit: 1:10,000; Life Technologies A-11122). The secondary antibodies used in all experiments except Figure S1 and Figure S6 were anti-mouse-Cy5 (1:250; Life Technologies A-10524) and anti-rabbit-Alexa Fluor 488 (1:500; Life Technologies A-11034). In Figure S1, the secondary antibody was anti-mouse-Alexa Fluor 488 (1:500; Life Technologies A-11001) and in Figure S6, the secondary antibodies were anti-mouse-Alexa Fluor 594 (1:500; Life Technologies A-11005) and anti-rabbit-Alexa Fluor 488 (1:500; Life Technologies A-11034).

### Purification of nc82/Brp monoclonal antibody

Hybridoma cells expressing monoclonal antibody nc82 (Antibody Registry ID: AB_2314866), which recognizes the *Drosophila melanogaster* Brp protein (Wagh et al., 2006) developed by Erich Buchner were obtained from the Developmental Studies Hybridoma Bank, created by the NICHD of the NIH and maintained at The University of Iowa, Department of Biology, Iowa City, IA 52242. Frances Weis-Garcia and the members of the MSKCC Antibody and Bioresource Core Facility subsequently used these hybridoma cells to purify this monoclonal antibody. The hybridoma was adapted to Gibco™ Hybridoma-SFM (Cat # 12045084) and 1% fetal bovine serum prescreened for ultra-low levels of bovine Ig. Antibody expression was confirmed and the adapted hybridoma was inoculated into the cell compartment of the Corning™ CELLine Disposable Bioreactor (Cat # 353137) in 15 ml of Hybridoma-SFM + 0.5% fetal bovine serum (production media) at 3 million viable cells / ml. The media compartment of the flask contained 350 ml of production media. The bioreactor was incubated at 37°C with 7% CO_2_ for 3 days, at which time the cells and media containing nc82 were harvested. 30 million viable cells from the harvest were re-inoculated back into the cell compartment in 30 ml fresh production media. The media in the media compartment was replaced the following day with 650 ml production media. Three days later, the media in the media compartment was replaced with 1,000 ml production media, with the next harvest 3 days later (7 days after the previous harvest). Cells were harvested weekly and fed bi-weekly until the desired amount of monoclonal antibody was reached. After the first harvest, each one contained about 3 mg of monoclonal antibody nc82/ml production media. The harvests to be purified were pooled, centrifuged at 12,855 x g for 15 min. 6.5 mg / run were loaded onto a Cytiva (formerly GE Life Sciences) 1 ml HiTrap Protein G HP antibody purification column (Cat # 29048581) at 1 ml / min. The column was then washed with 0.02 M Sodium Phosphate (pH 7.0) before the monoclonal antibody was eluted with 0.1 M Glycine-HCl (pH 2.7). One ml fractions were collected and immediately neutralized with 60 ml of 1.0 M Tris-HCl (pH 9.0). The harvest, flow through and fractions from the peak were run on an a 10% SDS-PAGE (Bio-Rad Cat # 345-0010) to confirm purity and determine which should be pooled. The pooled fractions of monoclonal antibody were dialyzed into PBS overnight using dialysis tubing (Spectrum™ 132544) with a 50 kDa MWCO. Another 10% SDS-PAGE was run, and the concentration determined using the absorbance at 280 using an extinction coefficient of 1.43.

### Generation of the IR25a polyclonal antibody

Rabbit polyclonal antibodies were raised against IR25a by Proteintech Group Inc. Antibodies were raised against a protein fusion of the 67 C-terminal amino acids of IR25a and glutathione S-transferase. cDNA corresponding to the C-terminal region was inserted into the expression vector PGEX-4T using primers TTTTGGATCCAAATACCGCAAGAACGTAAAG and TTTTCTCGAGTTAGAAACGAGATTTAAAGTTG and expressed in bacterial strain BL21. A purified 31 kDA fusion protein was used to immunize 2 rabbits. Serum was affinity purified to a final concentration of 450 µg/mL and tested by whole mount antenna immunostaining comparing *+/+^LVP^* to *IR25a^BamHI/BamHI^*. Antibodies from one of the two rabbits were found to selectively label *+/+^LVP^* antennae, and only this antibody was used in all further studies.

### Female Antennal lobe confocal imaging

All brains were imaged using a Zeiss Inverted LSM 880 laser scanning confocal microscope with a 25x / 0.8 NA immersion-corrected objective unless otherwise noted. Glycerol was used as the immersion medium to most closely match the refractive index of the mounting medium Vectashield. Antennal lobes in Figure 1, Figure 2, Figure 6, Figure S2-S8 were imaged at either 1024 x 1024 or 2048 x 2048 pixel resolution in X and Y with 0.5 µm Z-steps for a final voxel size of either 0.0615 x 0.0615 x 0.5 µm^3^ or 0.1230 x 0.1230 x 0.5 µm^3^. Both conditions oversampled relative to the objective resolution and no differences were noted between imaging conditions. The laser intensity and gain were adjusted along the Z-axis to account for a loss of intensity due to depth and care was taken to avoid saturation and ensure that the deepest glomeruli were visible for segmentation. We note that all confocal imaging was conducted in a manner that would maximize our ability to visualize the boundaries between glomeruli and to determine the presence or absence of a given fluorophore in each glomerulus, and was not intended as a quantitative measure of fluorescence intensity. *3xP3* was used as a promoter to express fluorescent proteins as markers for the knock-ins and *QUAS* transgenes used in this study, and care was taken to distinguish expression derived from the *3XP3* promoter from the expression of the QF2 driver and *QUAS* effector lines under investigation. *3xP3* drives expression in the optic lobes, as well as some cells in the dorsal brain. Neither area overlaps with the antennal lobes. As reported previously (Matthews et al., 2019), we saw no *3xP3*-driven expression in the antennal lobes in the reporter lines alone (data not shown). Representative antennal lobe images presented in the figures were cropped to remove *3xP3*-driven expression elsewhere in the brain.

### Male brain confocal imaging

All male brains (Figure 2M,O,Q,S,U) were imaged using a Zeiss Inverted LSM 880 laser scanning confocal microscope with a 25x / 0.8 NA immersion-corrected objective. Glycerol was used as the immersion medium to most closely match the refractive index of the mounting medium Vectashield. Brains were imaged at 1024 x 1024 pixel resolution in X and Y with 0.5 µm Z-steps for a final voxel size of 0.2372 x 0.2372 x 0.5 µm^3^. The laser intensity and gain were adjusted along the Z-axis to account for a loss of intensity due to depth and care was taken to avoid saturation and ensure that the deepest regions of the brain were visible. Confocal images of the brain were processed in ImageJ/FIJI (NIH).

### Female subesophageal zone confocal imaging

All female subesophageal zones (Figure 3B,D) were imaged using a Zeiss Inverted LSM 880 laser scanning confocal microscope with a 25x / 0.8 NA immersion-corrected objective. Glycerol was used as the immersion medium to most closely match the refractive index of the mounting medium Vectashield. Brains were imaged at 1024 x 1024 pixel resolution in X and Y with 0.5 µm Z-steps for a final voxel size of 0.2076 x 0.2076 x 0.5 µm^3^. The laser intensity and gain were adjusted along the Z-axis to account for a loss of intensity due to depth and care was taken to avoid saturation and ensure that the deepest regions of the subesophageal zone were visible. Confocal images of the subesophageal zone were processed in ImageJ/FIJI (NIH).

### Antennal lobe glomerulus quantification

Confocal images of the antennal lobes in Figure 1, Figure 2, Figure 6, Figure S2-S8 were processed in ImageJ/FIJI (NIH). The number of glomeruli was quantified as follows: a single region of interest (ROI) was manually drawn around each glomerulus at a section approximately central along the Z-axis. Every glomerulus was outlined and an ROI set was collected that contained the outlines of all glomeruli. Glomeruli were then separated into two groups, GFP-positive and GFP-negative glomeruli. A count of each was made to determine the number of glomeruli labeled by each line as well as the total number of glomeruli. The ROIs were flattened along the Z-axis to enable representation of the data in two dimensions in Figure 1, Figure 2, Figure S2-S5, Figure S7. The left antennal lobe in 3 brains was analyzed for each genotype in Figure 1 except for *Gr3*, for which the left antennal lobe was analyzed in 1 brain, and both left and right antennal lobes were analyzed in an additional 4 brains in Figure S6. Although we were able to recognize general regions of the antennal lobe, the interindividual variability made it impossible to identify most glomeruli by shape alone. We therefore have not attempted to name and number every glomerulus in *Aedes aegypti* as has been done in previous studies (Ignell et al., 2005; Shankar and McMeniman, 2020). As noted by Ito et al. (Ito et al., 2014), there is considerable confusion about the use of coordinate axes in the brains of animals in general and insects in particular. The glomeruli in the antennal lobe of *Aedes aegypti* were originally named by Ignell et al. (Ignell et al., 2005) using a set of coordinate axes that differ from those consistently used in *Drosophila melanogaster* (Couto et al., 2005; Fishilevich and Vosshall, 2005; Grabe et al., 2015; Laissue et al., 1999; Stocker et al., 1990). A recent study of the antennal lobe of *Aedes aegypti* renamed glomeruli to account for this discrepancy in coordinate axes (Shankar and McMeniman, 2020), and throughout this paper we use the same coordinate axes they have implemented. While Shankar and McMeniman renamed most antennal lobe regions and glomeruli, they chose not to rename the MD (Medio-Dorsal) cluster of glomeruli comprising MD1, MD2, and MD3 whose sensory input derives from the maxillary palp. We have observed in our study that the MD glomeruli are medial, but they are not notably dorsal, and therefore refer to them as Glomerulus 1, Glomerulus 2, and Glomerulus 3 in this paper for simplicity. While there is utility in naming glomeruli, we suspect that the *Aedes aegypti* mosquito antennal lobe atlas will be refined in the future with the advent of new genetic tools that will unambiguously allow the field to distinguish and name genetically identifiable glomeruli. We found that the size, shape, and number of antennal lobe glomeruli in *Aedes aegypti* was variable from animal to animal. It is possible that the boundaries between glomeruli are not easily distinguished by synaptic staining and that specific glomeruli will become identifiable once there are genetic tools available that label smaller populations of olfactory sensory neurons. The anatomical variability we see is consistent with both the original map that identified 50 glomeruli (Ignell et al., 2005), which divided glomeruli into 3 classes based on their variability in location, as well as a recent study that looked specifically at the size and shape of glomeruli across animals (Shankar and McMeniman, 2020) and revised the original map to a count of ∼80 glomeruli. Shankar and McMeniman named and numbered these glomeruli across animals, but they noted that they were only able to consistently identify 63 glomeruli. This is similar to the ∼65 glomeruli we observed in our work. While there is not yet a clear consensus on the exact number of antennal lobe glomeruli in *Aedes aegypti*, the number of chemosensory receptors expressed in the antenna and maxillary palp is at least twice as large as any of the estimates of glomerulus number. The variability in antennal lobe structure appears at first to contrast with *Drosophila melanogaster*, where each glomerulus can be clearly identified and named. However, we note that the antennal lobe map in *Drosophila melanogaster* has been refined with the advent of new genetic techniques, starting with 35 glomeruli in the original atlas (Stocker et al., 1990), then modified to 40 glomeruli (Laissue et al., 1999), and further refined in numerous studies (Couto et al., 2005; Fishilevich and Vosshall, 2005; Tanaka et al., 2012) including a recent count of 54 (Grabe et al., 2015) and 58 (Task et al., 2021) glomeruli. We have refrained from naming glomeruli in *Aedes aegypti* at this time because we believe that a more stereotyped arrangement will emerge as new genetic lines are generated that allow cell-type-specific labelling. A recent study in the mosquito *Anopheles gambiae* using mosquitoes that label *Orco*- expressing olfactory neurons also noted that the antennal lobe was variable between animals relative to *Drosophila melanogaster* (Riabinina et al., 2016). It is therefore possible that mosquito antennal lobes are more variable than Drosophilids (Grabe et al., 2015; Prieto-Godino et al., 2017). Variability in olfactory bulb structure is seen even in the mouse, *Mus musculus,* where the principles of olfactory organization were first established (Schaefer et al., 2001; Strotmann et al., 2000; Zou et al., 2009). The exact size and location of glomeruli can vary between animals more than initially appreciated and appears to be determined by both genetic factors and activity in olfactory sensory neurons during the early life of the animal. In *Drosophila melanogaster*, glomerulus size is highly genetically determined and correlates strongly with the number of olfactory sensory neurons that innervates each glomerulus (Grabe et al., 2015). Whether the variability in glomerulus size in the mosquito is due to activity-dependent changes in structure or other factors remains to be seen.

### Additional technical notes on expression and projection patterns of chemosensory receptor knock-in strains

#### Orco-QF2>QUAS-mCD8:GFP

We noted that the intensity of GFP varies between glomeruli in this driver line, with some bright and others comparably dim. We speculate that this is due to a combination of the variability in *Orco* expression levels in individual neurons and variability in the density of innervation in individual glomeruli. A large region of the anterior ventral antennal lobe was previously referred to as the Johnston’s organ center and was thought to comprise a single large glomerulus (Ignell et al., 2005). In other insect species, Johnston’s organ mediates detection of auditory cues. Consistent with a recent study (Shankar and McMeniman, 2020), we segmented this region into multiple glomeruli based on anatomical boundaries revealed with Brp immunofluorescence. Glomeruli in this region are innervated by *Orco*-expressing neurons, calling into doubt the original report that these glomeruli process auditory stimuli and suggesting instead that they serve an olfactory function. In support of this hypothesis, the analogous area of the *Anopheles coluzzii* antennal lobe has been shown to receive projections from *Orco*-expressing olfactory sensory neurons (Riabinina et al., 2016). We also observed GFP projections into the subesophageal zone in *Orco-QF2>QUAS-mCD8:GFP* animals, which appear to derive from expression in the proboscis, the primary taste organ in insects. This is consistent with similar expression in *Anopheles coluzzii* (Riabinina et al., 2016) and functional data in *Anopheles gambiae* showing that olfactory responses are detected in this gustatory organ (Kwon et al., 2006).

#### Ir25a-QF2>QUAS-mCD8:GFP

The intensity of GFP projections varies between glomeruli in this driver line, with some bright and other comparably dim, as noted for *Orco-QF2*. The brightest glomeruli are primarily medial and anterior. We see the dimmest innervation in the area previously described as Johnston’s organ center as well as in the central antennal lobe. Labeling was also seen in other areas of the brain, most notably the subesophageal zone and anterior mechanosensory motor center.

#### Ir8a-QF2>QUAS-mCD8:GFP

Depending on the brain being analyzed there were either 2 or 3 medial glomeruli labelled in this line. In the cases where there were 3 medial glomeruli, this third medial glomerulus was innervated by a few large-diameter axons. These were larger and sparser than the smaller axons that densely innervated most other glomeruli in this line. We also note that there are 2-3 cell bodies that express GFP located in the cell body rind lateral to the antennal lobe (rALl). We are unable to definitively describe where these cells project without genetic reagents that selectively label these cells, but they appear to send bilateral processes that cross the midline within what appears to be the saddle to innervate the anterior mechanosensory motor center outside the antennal lobe. All naming is in accordance with the new insect brain nomenclature presented in Ito et al. (Ito et al., 2014).

#### Ir76b-QF2>QUAS-mCD8:GFP

In addition to projections to the antennal lobe, this line shows innervation of the subesophageal zone of the brain.

#### Gr3-QF2>QUAS-mCD8:GFP

All antennal lobes in this line show innervation of a single glomerulus (also referred to as ”MD1” and here referred to as “Glomerulus 1”; (Ignell et al., 2005; Shankar and McMeniman, 2020). In several brains, we saw a second small medial glomerulus that derives its innervation from the antenna and is in a small medial cluster of landmark glomeruli midway down the anterior-posterior axis closest to the center of the brain. Innervation appears to come from only a few axons. This low and variable reporter expression is consistent with the low level of expression of *Gr3* in the antennal transcriptome (Matthews et al., 2016). Because this line only shows innervation of these 1-2 glomeruli, we analyzed all glomeruli only in the single brain in Figure 1I, and additionally analyzed 8 more antennal lobes in 4 brains for the presence or absence of labelling in these two glomeruli. We analyzed both left and right antennal lobes from 4 brains and found that in 3 of the 4 brains there was a second glomerulus in one or both antennal lobes (Figure S6). The presence of the second glomerulus was not specific within a single animal as we found all variations of presence and absence of this glomerulus across both antennal lobes in these 4 animals. In some *Gr3-QF2>QUAS-mCD8:GFP* animals, we detected a small number of processes that extended beyond the antennal lobe and into the higher brain, although the exact termination site varied. We never saw CO_2_-evoked activity in the variable second glomerulus or these projections outside the antennal lobe. Images in Figure S6 were taken as described above with the following changes: Secondary antibodies used were anti-mouse-Alexa Fluor 594 (1:500; Life Technologies A-11005) and anti-Rabbit-Alexa Fluor 488 (1:500; Life Technologies A-11034). Images were taken using a Zeiss Inverted LSM 880 laser scanning confocal microscope with a Plan-Apochromat 40x/1.4 Oil DIC objective. Images were taken at 1024 x 1024 in XY to generate images with a final voxel size of 0.1384 x 0.1384 x 0.5 µm^3^. Images were scored as containing GFP in one or two glomeruli.

### Additional technical notes on expression and projection patterns of Split-QF2 strains

All antennal lobe immunostaining in Figure 2, Figure 6, Figure S7, Figure S8 was carried out as described above with slight modifications to utilize the *15xQUAS-dTomato-T2A-GCaMP6s* effector line. The same primary antibodies were used because of the structural similarity between GCaMP6s and GFP. Intrinsic dTomato was detected without antibody amplification, as it retained fluorescence after fixation and staining. Brp (Cy5), dTomato, and GCaMP6s (Alexa Fluor 488) were imaged as three separate confocal channels as described above. Glomeruli labelled by dTomato completely overlapped with those labelled by GCaMP6s immunofluorescence, so both channels were used during the quantification of positive and negative glomeruli. dTomato labeling was used to generate sample images. There was no staining in the antennal lobes of the individual split effector lines crossed to *15xQUAS-dTomato-T2A-GCaMP6s* (n=3 per genotype) (Figure 2, Figure S7).

### Antennal lobe anterograde dye fill

For images in Figure S1, mosquitoes were anesthetized on wet ice until immobile and then transferred to a cold dissection dish. A single antenna or maxillary palp was loaded with Texas-red conjugated dextran (Molecular Probes D3328) diluted 10 mg in 100 μL external saline (103 mM NaCl, 3 mM KCl, 5 mM 2-[Tris(hydroxymethyl)methyl]-2-aminoethanesulfonic acid (TES), 1.5 mM CaCl_2_, 4 mM MgCl_2_, 26 mM NaHCO_3_, 1 mM NaH_2_PO_4_, 10 mM trehalose, 10 mM glucose, pH 7.3, osmolality adjusted to 275 mOsm/kg). To load the dye a small drop (approximately 0.5-1 µL) of dye was placed onto the surface of the dish and the animal was moved such that the intended cut-site on a single antenna or maxillary palp was placed in the drop of dye. The antenna or maxillary palp was then removed with sharp forceps and a fine scalpel (F.S.T 10315-12) while it was submerged in the dye. Care was taken to remove the maxillary palp proximal to the fourth segment, to include all the capitate-peg sensilla, and to remove the antenna near the base but to leave the antennal pedicel completely intact. The animal remained immobile on ice with the antenna or maxillary palp submerged and the dye was loaded for 2-5 min. After this time the animal was placed in a small soup cup with access to 10% sucrose and returned to standard rearing conditions overnight to give the dye time to diffuse throughout the neurons and fill the length of the axon. The next morning dissection of adult brains and immunostaining was carried out as described above.

### Antennal lobe 3-D reconstructions

In an attempt to develop a map of the *Aedes aegypti* antennal lobe, 3 brains from the *+/+^LVP^* strain were immunolabeled with Brp to identify the boundaries between antennal lobe glomeruli. The left antennal lobe in each brain was independently reconstructed from confocal sections taken with a Plan-Apochromat 63x/1.40NA oil immersion objective, at 1024 x 1024 pixel resolution in X and Y with 0.5 µm Z-steps for a final voxel size of either 0.1318 x 0.1318 x 0.5 µm^3^ using the software Imaris (Bitplane). Although the area previously termed Johnston’s organ center was considered a single glomerulus in a previous study (Ignell et al., 2005), we noted anatomical boundaries in this region, suggesting that it contains multiple glomeruli. This observation is consistent with recently published work (Shankar and McMeniman, 2020) and this area was segmented by an individual researcher to generate the final reconstructions. Two of these are shown in Figure S1. Each glomerulus was manually segmented into an individual surface using Surpass View. We were consistently able to identify the three glomeruli innervated by the maxillary palp, previously termed MD1, MD2 and MD3 (Ignell et al., 2005) which we refer to in this study as Glomerulus 1, Glomerulus 2, and Glomerulus 3 (Figure 1, Figure 6). The overall structure of the antennal lobe varied considerably from animal to animal and although we were able to identify certain regions and certain landmark glomeruli including those that are targeted by the maxillary palp, we were unable to assign an unambiguous identity to every glomerulus, as is possible in *Drosophila melanogaster* (Couto et al., 2005; Fishilevich and Vosshall, 2005). This variability makes it essentially impossible to identify a given glomerulus between animals and we therefore have decided to avoid referring to glomeruli by previous naming schemes, including MD1, MD2, MD3. An authoritative atlas of the *Aedes aegypti* antennal lobe awaits genetic reagents that label subpopulations of sensory neurons that will permit the field to refer to glomeruli by their molecular identity.

### Antennal whole mount immunofluorescence

Whole-mount immunostaining of adult antennae was performed as described (Riabinina et al., 2016) with modifications. 7-11 day-old Liverpool mosquitoes were immobilized on ice, decapitated and heads and placed in 1 mL ZnFA fixative solution (0.25% ZnCl_2_, 2% paraformaldehyde, 135 mM NaCl, 1.2% sucrose and 0.03% Triton X-100) for 20–24 h at room temperature in the dark. Next, the heads were washed three times for 30 min each with HBS buffer (150 mM NaCl, 5 mM KCl, 25 mM sucrose, 10 mM HEPES, 5 mM CaCl_2_ and 0.03% Triton X-100). Antennae were carefully removed in HBS on ice and placed in 400 μL HBS in 0.5 mL Eppendorf tubes. After a brief wash in HBS, the tissue was incubated in 400 μL 80% methanol/20% dimethyl sulfoxide (DMSO) solution for 1 hr at room temperature, washed for 5 min in 400 μL 0.1 M Tris pH 7.4, 0.03% Triton X-100 solution and incubated in 400 μL blocking solution (PBS, 5% normal goat serum (Jackson 005-000-121), 1% DMSO and 0.3% Triton X-100) for at least 3 hr at room temperature or overnight at 4°C. Next, the tissue was placed in a 0.5 mL Eppendorf tubes containing 400 μL blocking solution with primary antibodies [rabbit anti-Orco EC2 (Larsson et al., 2004), 1:50, Vosshall lab; chicken anti-GFP,1:200, Aves GFP-1020] and submerged and held in a water bath sonicator (Branson m1800) for 30 sec at the high setting. Next, the tubes were placed on a rotator for 2 days at 4°C in the dark, after which the sonication procedure was repeated. The tubes were placed on a rotator for 2 additional days (for a total of 4 days) at 4°C in the dark. Next, the tissue was washed 5X 30 min each at room temperature in PBS, 1% DMSO and 0.3% Triton X-100. Secondary antibodies (anti-rabbit Alexa Fluor 555 Plus, 1:200, Thermo Fisher A-32732, anti-chicken Alexa Fluor 488, 1:200, Thermo Fisher A-11039) and nuclear dye (TO-PRO-3 Iodide, 1:400, Thermo Fisher T3605) were added to the blocking solution, and tubes were sonicated as described above and incubated for 4 days at 4°C in the dark with the sonication repeated after 2 days of incubation. The tissue was then washed 5X 30 min at room temperature in PBS, 1% DMSO and 0.3% Triton X-100, rinsed in PBS and mounted in Slow Fade Diamond for confocal imaging.

### Antennal whole-mount immunostaining with Ir25a antibody

This protocol was performed as previously described (Basrur et al., 2020) with modifications. Six- to 11-day-old female mosquitoes were anesthetized on wet ice, decapitated, and placed in 1.5 mL 5 U/mL chitinase (Sigma C6137) and 100 U/mL chymotrypsin (Sigma CHY5S) in 119 mM NaCl, 48 mM KCl, 2 mM CaCl2, 2 mM MgCl2, 25 mM HEPES, 1% DMSO buffer on ice. Heads were incubated on a ThermoMixer (Eppendorf 5382000023) at 37°C for 5 min, followed by 55 min in a rotating hybridization oven at 37°C. Heads were then rinsed once and fixed in 4% paraformaldehyde, 1X Ca+2, Mg+2 free PBS, and 0.03% Triton X-100 for 24 hr at 4°C on a rotator. All subsequent 4°C steps used a nutator, and room temperature steps used a rotator. Heads were washed for 30 min at room temperature at least three times in 1X PBS with 0.03% Triton X-100 (0.03% PBT). Antennae were then dissected into 0.5-mL microfuge tubes and dehydrated in 80% methanol/20% DMSO for 1 hr at room temperature. Antennae were washed in 0.03% PBT for 30 min at room temperature, and blocked/permeabilized in 1X PBS, 1% DMSO (Sigma 472301), 5% normal goat serum, 4% Triton X-100 for 24 hr at 4°C. Antennae were washed for 30 min at least five times with 0.03% PBT, 1% DMSO, 5% normal goat serum at 4°C, and then moved to primary antibody in 1X PBS, 1% DMSO, 5% normal goat serum, 0.03% Triton X-100 for 72 hr at 4°C. Primary antibodies used were mouse anti-*Apocrypta bakeri* Orco monoclonal antibody #15B2 (1:50 dilution, gift of Joel Butterwick and Vanessa Ruta), and rabbit anti-Ir25a (1:50 dilution). Orco monoclonal antibody and Ir25a polyclonal antibody specificities were verified in *Aedes aegypti* by staining *orco* mutant and *Ir25a* mutant antennae, respectively (Figure 4E-H). Antennae were washed for 30 min at least five times with 0.03% PBT, 1% DMSO at room temperature, and then washed overnight in the same solution. Antennae were then moved to secondary antibody (1:200) in 1X PBS, 1% DMSO, 5% normal goat serum, 0.03% Triton X-100 for 72 hr at 4°C. Secondary antibodies used were goat anti-mouse Alexa Fluor 488 (Thermo A-11001) and goat anti-rabbit Alexa Fluor 555 Plus (Thermo A32732). Antennae were washed for 30 min at least five times with 0.03% PBT, 1% DMSO at room temperature, and then washed overnight in the same solution. Antennae were rinsed in 1X PBS, rinsed three times in Slowfade Diamond (Thermo S36972), and mounted in Slowfade Diamond.

### Whole mount antennal and maxillary palp RNA *in situ* hybridization

RNA was detected in whole mount antenna and maxillary palp using the hybridization chain reaction (HCR) technique as previously described (Choi et al., 2018) with modifications. Probes, amplifiers, Probe Hybridization Buffer, Amplification Buffer, and Probe Wash Buffer were purchased from Molecular Instruments. Full list of probe lot numbers can be found in Data File 1. 5-8 day-old Liverpool mosquitoes were anesthetized on wet ice, manually decapitated with forceps, and heads with antennae and the proboscis were digested in a chitinase-chymotrypsin solution (119 mM NaCl, 48 mM KCl, 2 mM CaCl_2_, 2 mM MgCl_2_, 25 mM HEPES, 5 U/mL chitinase (Sigma-Aldrich C6137-50UN), 100 U/mL alpha-chymotrypsin (Sigma-Aldrich CHY5S-10VL), 2% DMSO) (Manning and Doe, 2017) at 37°C for 30 min (antennae) or 1 hr (maxillary palps) in a Fisher Isotemp oven and subsequently fixed in 4% paraformaldehyde, 1X PBS, 0.03% Triton X-100 on a rotator at 4°C overnight. Heads were washed 4 times on ice for 10 min each in 0.1% PBS-Tween-20. Antennae or maxillary palps were dissected in 0.1% PBS-Tween-20 on ice and dehydrated with a graded series of methanol/0.1% PBS-Tween: 25% methanol in 0.1% PBS-Tween-20 for 10 min on ice, 50% methanol in 0.1% PBS-Tween-20 for 10 min on ice, 75% methanol in 0.1% PBS-Tween-20 for 10 min on ice, and two washes of 100% methanol for 10 min on ice. Tissues were incubated overnight in 100% methanol at -20°C and were subsequently rehydrated with a series of graded methanol/0.1% PBS-Tween-20: 75% methanol in 0.1% PBS-Tween-20 for 10 min on ice, 50% methanol in 0.1% PBS-Tween-20 for 10 min on ice, 25% methanol in 0.1% PBS-Tween-20 for 10 min on ice, and two washes of 0.1% PBS-Tween-20 for 10 min each on ice. Tissue was digested in 20 µg/mL Proteinase-K (Thermo Fisher AM2548) in 0.1% PBS-Tween for 30 min at room temperature and washed twice with 0.1% PBS-Tween-20 for 10 min each at room temperature. Tissue was fixed in 4% paraformaldehyde in 0.1% PBS-Tween-20 for 20 min at room temperature and washed 3 times for 10 min each in 0.1% PBS-Tween-20 at room temperature. Tissue was incubated in Probe Hybridization Buffer at room temperature for 5 min and then in 37°C pre-warmed Probe Hybridization Buffer rotating in a hybridization oven for 30 min. 8 pmol of each probe set was prepared in 37°C pre-warmed Probe Hybridization Buffer and tissue was incubated in probe solution at 37°C in a hybridization oven for 2 nights. Tissues were washed in 37°C pre-warmed Probe Wash Buffer 5 times for 10 min each at 37°C. Tissues were washed twice in 5X SSC 0.1% Tween-20 at room temperature for 10 min each. Tissues were pre-amplified in room temperature Amplification Buffer for 10 min. 18 pmol hairpins were separately prepared by heating 6 µL of 3 µM stock of hairpins H1 and H2 at 95°C for 90 sec on an Eppendorf Mastercycler and allowing to cool to room temperature in a dark drawer for 30 min. Hairpins were resuspended in 100 µL amplification buffer and tissues were incubated in this hairpin solution in the dark on a rotator at room temperature overnight. Tissues were washed 5 times for 10 min each in 5X SSC 0.1% Tween-20 and mounted in SlowFade Diamond (Thermo Fisher S36972) on glass slides with coverslips for confocal imaging.

### Whole mount antennal, maxillary palp, and proboscis dTomato visualization

7-14 day-old *Ir25a-QF2*, *Orco-QF2*, *Ir25a-QF2AD*, *Orco-QFDBD*, and *Ir25a-QF2AD Orco-QFDBD>15XQUAS-dTomato-T2A-GCaMP6s* mosquitoes were anesthetized on wet ice, manually decapitated with forceps and heads with antennae, proboscises, and maxillary palps were immediately fixed in 1 mL 4% paraformaldehyde, 1X PBS, 0.03% Triton X-100, on a rotator in the dark at 4°C overnight. Heads were washed 3X 30 min each in 1X PBS, 0.03% Triton X-100 at room temperature, then antennae, proboscises, and maxillary palps were carefully removed and placed in 1X PBS, 0.03% Triton X-100. Next, antennae, proboscises, and maxillary palps were placed in a solution of 1X PBS, 0.03% Triton X-100, 1% DMSO, and a 1:400 dilution of TO-PRO-3 (Thermo Fisher T3605) for 24 hr at 4°C in the dark. Antennae, proboscises, and maxillary palps were then washed 5X 30 min each in 1X PBS, 0.03% Triton X-100 at room temperature in the dark, washed once with 1X PBS, transferred to a well of SlowFade diamond to remove excess PBS, and mounted in SlowFade Diamond for confocal imaging.

### Antennal and maxillary palp confocal imaging and cell quantification

Images of peripheral tissues were acquired with a Zeiss Axio Observer Z1 Inverted LSM 880 NLO laser scanning confocal microscope (Zeiss) with a 25x/0.8 NA or 63x/1.4 NA immersion-corrected objective at a resolution of 3096 x 3096 pixels or 2048 x 2048 pixels. When comparing dTomato fluorescence across genotypes, image acquisition parameters were kept consistent. When necessary, tiled images were stitched with 20% overlap. We note that all confocal imaging was conducted in a manner that would maximize our ability to visualize the presence or absence of each fluorophore and was not intended as a quantitative measure of fluorescence intensity. Confocal images were processed in ImageJ (NIH). Because the antenna is a cylindrical structure, when whole antennal segments are mounted on a slide and imaged on a confocal microscope, signal can be easily detected from the region closest to the coverslip and confocal objective, but signal is weaker when imaging the side further from the coverslip and objective. For the purposes of consistent quantification, we only quantified cell numbers from the region closest to the coverslip (red rectangle in Figure S12A). For quantifying expression in the maxillary palp, only the dorso-lateral region of the 4^th^ maxillary palp segment was analyzed. (yellow rectangle in Figure S12B). Quantification of co-expression in antennae and maxillary palps was done in ImageJ (NIH) using the Cell Counter plugin. Cells in each channel were manually marked independently of the signal in the other channels. After cells in each channel are marked, and markers were then merged. Cells that were labeled with multiple markers (co-expressing cells) were then marked with a third marker (Figure S12C-H). Cell counts were then imported into Microsoft Excel and R for analysis.

### Antenna Dissection for snRNA-seq

Approximately 100-250 female *+/+^LVP^* mosquitoes aged 6-8 days post-eclosion were anesthetized on wet ice for 10 min. Mosquitoes were then placed in a 70 µm cell strainer (Falcon 08-771-1). The cell strainer containing the anesthetized mosquitoes was placed in a 60 mm Petri dish (Corning 430166), and ice-cold molecular-grade 100% ethanol was gently poured into the cell strainer for 5 sec. The cell strainer with ethanol-rinsed mosquitoes was then transferred to a new 60 mm Petri dish and ice-cold Schneider’s Medium (Gibco 21720024) was poured into the cell strainer to rinse. Approximately 20 mL of ice-cold Schneider’s medium was poured into a 100 mm Petri dish (Corning 430293) on wet ice or reusable ice pack (Cooler Shock, mid-size freeze pack). Schneider’s Medium-rinsed mosquitoes were transferred from the cell strainer to the 100 mm Petri dish. A new 70 µm cell strainer (pluriSelect 43-10070) with walls trimmed with a sterile razor blade to a height of 0.5 – 0.75 cm was placed into the same 100 mm Petri dish. The antennae were then removed using forceps and placed into the cell strainer. Antennae were rinsed approximately every 10 min by agitating the cell strainer and pipetting fresh ice-cold Schneider’s Medium into the cell strainer. Dissection of each sample was limited to 90 min to ensure nuclei integrity, and when 90 min elapsed or all mosquitoes dissected, antennae were transferred into a DNA LoBind 1.5 mL tube (Eppendorf 022431021) pre-wet with Schneider’s Medium. The cell strainer with antennae was inverted with forceps into the tube and approximately 300 µL ice-cold Schneider’s Medium was pipetted into the cell strainer to release antennae into the Eppendorf tube. The sample was then flash-frozen in liquid nitrogen and stored at -70°C until ready for nuclei extraction. A total of approximately 1000-1500 antennae were collected for each snRNA-seq batch, collected across four dissection sessions. Two batches of female antennae were processed for the snRNA-seq data presented in this paper. All tissue was collected at Rockefeller University. Batch 1 was processed at Rockefeller University (including nuclei extraction, 10X Genomics run, library preparation and sequencing), and Batch 2 was shipped on dry ice and processed at Baylor College of Medicine.

### Batch 1 (Rockefeller antenna sample) nuclei extraction

Nuclei extraction of mosquito antennae was performed as previously described (McLaughlin et al., 2021) with modifications. Dissected antennae were thawed on wet ice, and all subsequent steps were performed on wet ice unless otherwise noted. Once samples were thawed, antennae were centrifuged in a benchtop microcentrifuge for 5-10 sec, Schneider’s Medium was removed and replaced with 100 µL of homogenization buffer (McLaughlin et al., 2021). Antennae from multiple dissection sessions were combined into a single DNA LoBind 1.5 mL tube using a low-retention repel polymer technology 200 µL filter tip (TipOne 11821830), with ∼1 mm from the distal end trimmed using a sterilized and RNAse away-treated (Thermo Fisher 7000TS1) razor blade. With no more than 500 µL buffer present in the tube, tissue was ground for 30-60 sec with a pellet pestle motor (Kimble 749540-0000) and RNase-free pestle (Kimble 749521-0590). The volume of buffer was brought up to 1000 µL with additional ice-cold homogenization buffer. Next, a 1 mL Dounce tissue grinder and pestle set (Wheaton 357538) that had been autoclaved at 121°C for 4 hr the previous day was pre-wetted with homogenization buffer. Using a low-retention (repel polymer technology) 1000 µL filter tip (TipOne 11821830), samples were transferred into the Dounce homogenizer. Nuclei were released by homogenizing with 20 strokes of the loose pestle, and 40 strokes of the tight pestle. Next, a low-retention 1000 µL tip was used to remove ∼500 µL of the suspension. The suspension was filtered through a 40 µm Flowmi filter (Bel-Art H13680-0040) into a pre-wet 20 µm PluriStrainer (pluriSelect 43-10020-40) in a 1.5 mL LoBind Eppendorf tube. The second ∼500 µL antennae nuclei suspension was then filtered the same way into the same Eppendorf tube. The suspension was then divided equally into two 1.5 mL LoBind Eppendorf tubes and centrifuged for 10 min at 500xG at 4°C. The supernatant was gently discarded without disturbing the pellet. Next, pellets were resuspended in 100 µL 1X PBS, 1% bovine serum albumin, 10 µL/mL RNAse inhibitor (Roche RNAINH-RO) by pipetting 5 times with a low-retention 1000 µL tip, combined and pipetted to resuspend and break up cell clumps 15 more times. The suspension was then filtered three times by running it through a Flowmi filter into a 10 µm strainer (pluriSelect 43-10010-40) in a 1.5 mL LoBind Eppendorf tube. To ensure nuclei were not clumping, 10 µL of the suspension was removed and stained with acridine orange and propidium iodide (Logos Biosystems, LGBD10012). The concentration of nuclei was determined by counting cells on a Luna FX7 automated cell counter (Logos Biosystems L70001).

### Batch 1 (Rockefeller antenna sample): 10X Genomics, library preparation and sequencing

Single cell 3’ expression Libraries were generated using Chromium Single Cell 3’ Library & Gel Bead kit Version 3.1 (10X Genomics PN1000269). Standard protocols from 10X Genomics were followed to generate the dual index libraries. Due to the small nucleus size (4-5 µm in diameter), 17 cycles were used for cDNA amplification and 13 cycles for index PCR. The quality and quantity of the libraries were assessed on Agilent TapeStation, the library was sequenced on Illumina NovaSeq 6000 sequencer using 100 cycle SP flowcell and 800 million paired reads were generated (read 1 = 28 bp, read 2 = 90 bp).

### Batch 2 (Baylor antenna sample): nuclei extraction

Nuclei extraction from mosquito antennae were performed as previously described (Li et al., 2021) with modifications. Fresh homogenization buffer (Li et al., 2021) was prepared and kept on ice. Samples were thawed from -80°C on wet ice, spun down in 100 µL Schneider’s Medium using a bench top spinner, and as much medium as possible was discarded. Antennae from multiple dissection sessions were combined into a single 1.5 mL Eppendorf tube using a low-retention 200 µL filter tip (Rainin 30389240) with ∼1 mm from the distal end trimmed using a sterilized and RNAse away-treated (Thermo Fisher 7000TS1) razor blade (VWR 10835-965) and 100 µL Homogenization buffer was added. The sample was ground with a pestle motor (Kimble 6HAZ6) for 30 – 60 sec on wet ice. 900 µL homogenization buffer was added, and 1000 µL homogenized sample was transferred into the 1 mL Dounce tissue grinder set (Wheaton 357538) that had been autoclaved at 200°C for >5 hr or overnight a day in advance. Nuclei were released by 20 strokes with a loose Dounce pestle and 40 with a tight Dounce pestle on ice, taking care to avoid bubbles. 1000 µL of the sample was filtered through a 5 mL cell strainer (35 µm), and then filtered using 40 µm Flowmi (BelArt, H13680-0040) into 1.5 mL EP tube, centrifuged for 10 min at 1000xG at 4°C. The supernatant was discarded with care not to disturb the pellet. The nuclei were resuspended using 500 µL 1xPBS/0.5%BSA with RNase inhibitor (9.5 mL 1x PBS, 0.5 mL 10% BSA, 50 µL RNasin Plusby) pipetting at least 20 times to completely re-suspend the nuclei. Sample were filtered using 40 µm Flowmi into a new 5 mL fluorescence-activated cell sorting (FACS) tube and kept on wet ice.

### Batch 2 (Baylor antenna sample): FACS sorting, 10X Genomics, library preparation, sequencing

FACS sorting was done using a BD FACSAria III Cell Sorter to collect nuclei. Nuclei were stained with Hoechst-33342 (1:1000; >5 min). Hoechst-positive nuclei were collected into 1.5 mL Eppendorf tube with 500 µL 1x PBS with 0.5% BSA as the receiving buffer (RNase inhibitor added). For each 10X Genomics run, all nuclei were collected. Approximately 15,000 nuclei were collected from the antennae. Nuclei were spun for 10 min at 1000XG at 4°C, and then resuspended using 43.2 µL 1x PBS with 0.5% BSA (RNase inhibitor added). Since the yield of nuclei was low, all nuclei were loaded onto a 10X Genomics controller. 10X Genomics sequencing libraries were prepared following the standard protocol from 10X Genomics 3’ v3.1 kit with following settings. All PCR reactions were performed using the Biorad C1000 Touch Thermal cycler with 96-Deep Well Reaction Module. 13 cycles were used for cDNA amplification and 16 cycles were used for sample index PCR. As per 10X Genomics protocol, 1:10 dilutions of amplified cDNA and final libraries were evaluated on Agilent 4200 TapeStation. Single-cell RNA libraries were sequenced on Illumina NovaSeq 6000 sequencer with minimum sequencing depth of 50,000 reads/cell using the read lengths 28bp Read1, 8bp i7 Index, 91bp Read2.

### Maxillary palp dissection, nuclei extraction, FACS Sorting, 10X Genomics, library preparation, and sequencing

Maxillary palp dissections were conducted as described for the antenna. A total of 2,908 total maxillary palps were collected across twenty dissection sessions at Rockefeller. These samples were shipped on dry ice and processed at Baylor College of Medicine. Nuclei extraction and FACS was performed at Baylor as described for the Batch 2 antenna sample with approximately 7,000 nuclei collected. 10X Genomics, library preparation, and sequencing was done as described above for the Batch 2 antenna sample.

### snRNA-seq analysis: cell identification, ambient RNA removal, batch combination, and neuron classification

The *Aedes aegypti* genome (AaegL5.0, GCF_002204515.2 on NCBI) was indexed using Cell Ranger (version6.0.2). FASTQ files generated from 10X Genomics 3’ gene expression libraries were mapped to the indexed genomes and gene counts in each cell were calculated by CellRanger (version 6.0.2). Intron signals were included by specifying the --include-introns parameter for cellranger count.

DecontX from the celda package (version 1.8.1) was chosen for removing the ambient RNAs that are produced during the nuclei isolation. The raw and filtered reads generated from Cell Ranger were compared by DecontX to obtain decontaminated reads (Figure S9A, Figure S13A). The decontaminated reads were rounded by the R base::round function and the decontaminated matrices were generated by the DropletUtils package (version 1.12.3). Decontaminated expression matrices were loaded into the Seurat package (version 4.0.5) and multiplets were identified by DoubletFinder (version 2.0.3). The pK with maximum AUC was chosen for DoubletFinder. The multiplet numbers were estimated by the multiplet rate table on the 10X Genomics website. DoubletFinder-defined multiplets were excluded for the downstream analysis (Figure S9B, Figure S13B). Cells with extreme gene numbers or abundant mitochondria transcripts were removed using Seurat. We excluded nuclei expressing fewer than 400 or greater than 4000 genes. Nuclei with more than 5% of mitochondrial transcripts were excluded. Genes expressed in fewer than 3 nuclei were removed (Figures S9C-E, Figure S13C-E).

Expression matrices of remaining nuclei were loaded into the Seurat package and processed by Seurat (version 4.0.5). The analyses applied default parameters of Seurat unless specified. Expression matrices were normalized using NormalizeData() function. Highly variable genes were selected using FindVariableFeatures(). The data were scaled using the ScaleData() function with the vars.to.regress = c(’nCount_RNA’) parameter to regress out the effect of the total counts. The scaled data were dimensionally reduced using the RunPCA() function. t-distributed stochastic neighbor embedding (t-SNE) was used for visualizing the non-linear dimensionality reduction with 1 to 50 dimensions. Nuclei were clustered using the Louvain algorithm (Figure S9G, Figure 7C).

We performed two independent snRNA-seq experiments on the antenna to collect a large number of nuclei for our analysis. The two batches of antenna snRNA-seq data were merged and split using merge() and SplitObject() functions in Seurat. Split objects were normalized and selected for highly variable genes independently. To reduce the batch effects from two samples, we first selected genes for integrating two batches using the SelectIntegrationFeatures() function in Seurat (Figure S9F). Two batches were then integrated using the FindIntegrationAnchors() and IntegrateData() functions. Batch-corrected samples were then analyzed following the procedures described in the previous section from scaling to clustering to identify cluster-specific genes.

To classify cells as neurons, we first identified genes that are orthologous to the neuronal marker genes used in *Drosophila melanogaster* using pBLAST. Four mosquito genes, LOC5565901, LOC5570204, LOC5564848, and LOC5570381, are orthologous to the *Drosophila melanogaster* neural markers, *syt1*, *elav*, *CadN*, and *brp*, respectively. We saw that expression largely overlapped with the olfactory sensory neuron co-receptors *Orco*, *Ir25a*, *Ir76b*, *Ir8a*, and *Gr3*, consistent with the idea that these are neuronal markers. We defined neural clusters based on the expression of *syt1*, *elav*, *CadN*, and *brp*, and clusters expressing at least three neuronal markers in more than 50% of cells in the corresponding cluster were defined as neural clusters (Figure S9I, Figure S13H-I). These neural clusters were then examined for ligand-selective receptor and co-receptor expression.

### Antenna heat map

The normalized expressions of genes in all nuclei were utilized to plot heatmaps using the ComplexHeatmap package in R. Epithelia-, glia-, and neuron-enriched genes in the *Drosophila melanogaster* antenna were considered as references of the corresponding marker genes in *Aedes aegypti*.

#### Antenna tSNE plot

To generate tSNE plots in Figure 5C of all antennal nuclei and antennal neurons, expression matrices were first log-normalized, selected for highly variable genes, and scaled. Scaled data were applied to the RunTSNE() with 1 to 50 dimensions. All antenna nuclei and antennal neurons were clustered using the Louvain algorithm with resolutions 0.5 and 3 respectively (Figure S9G, Figure 5C).

#### Antenna dot plot

The dot plot of cluster-enriched chemosensory receptors in Figure 5D was based on the DotPlot() function in Seurat and customized using the ggplot2 package. The mean normalized expression and expression percentage of each chemosensory receptor were extracted by the DotPlot() function. Chemosensory receptors expressed in more than 35% of nuclei in the corresponding cluster with mean expression values (UMI of gene*10,000 / total UMI of cell +1) larger than 1 were considered cluster-dominant chemosensory receptors (Figure S10A-B). The expression percentages of all dominant chemosensory receptors were scaled and clustered. For visualizing differences lower-expressed ligand-selective receptor subunits, circles representing a mean expression value greater than 20 have the same color. The expression percentage and mean expression of each chemosensory receptor were plotted using the geom_point() function in ggplot2. The hclust() function was used to cluster genes.

#### Antenna chord plot

The chord plot of co-expressed chemosensory receptor in Figure 5E was generated using the chorddiag package in R. Normalized expressions of the top 20 expressed chemosensory receptor were examined for the co-expression in the antenna neuron population. Receptors that express more than 1 normalization value were considered as positively expressed. Each expressed chemosensory receptor was iteratively compared to the expression of the remaining 19 chemosensory receptor in the corresponding nuclei. If more than 20 nuclei expressed two chemosensory receptors simultaneously, these two receptors were considered as co-expressed chemosensory receptors and visualized using the chorddiag package.

#### Antenna scatter plot

The co-expression scatter plot in Figure S11F was based on the normalization expression from each single nuclei for a pair of chemosensory receptors. The normalization values were plotted using the geom_point() function of the ggplot2 package in R.

#### Antenna simplified co-expression heatmap

Heatmaps were generated for the purposes of visualizing examples co-expression patterns from a large dataset of 6,645 neurons and 231 chemosensory receptors (106 ORs, 73 IRs, and 52 GRs). For each chemosensory receptor, cells expressing the given gene above a normalized expression level threshold of 0.5 log(UMI*10,000+1) were identified and subsetted from the neural population (Figure S11A-B). Background noise of chemosensory receptor expression made it difficult to identify robust patterns of co-expression. Therefore, for the purpose of sorting cells to visualize expression patterns, we reclustered chemosensory receptor-expressing cell population using FindNeighbors(), RunPCA() with 1 to 50 dimensions and FindClusters() with a resolution of 3. This unsupervised clustering grouped cells based on their whole transcriptome, not solely based on chemosensory receptor expression. In many cases, cells in a cluster exhibited the same co-expression patterns. In heatmaps illustrating all the chemosensory receptors using scaled expression levels, these expression patterns were visible by eye (Figure S11B). Clusters exhibiting representative patterns were selected from the population, and heatmaps were re-generated for simplified co-expression heatmaps in Figure 5F-I and Figure S11C-F. For example, for analysis of *Ir41k* co-expression patterns: 412 cells were selected from the population with a normalized expression threshold above 0.5 log(UMI*10,000+1), and clustered using 50 principle components and a cluster resolution of 3 (Figure S11B). 6 clusters were identified (2, 3, 4, 6, 7, 8) as examples to illustrate co-expression patterns. In the simplified heatmap for *Ir41k* only, cells were reclustered using 9 principal components and a 1.9 clustering resolution (Figure 5F). In all other simplified heatmaps, the same clusters were used as identified in the larger co-expression heatmap.

### Maxillary palp: tSNE, heatmap, chord plot, expression feature plot

The tSNE of maxillary palp nuclei in Figure 7C was generated similarly to the antenna tSNE, with cluster resolution adjusted to 2.5. The heatmap of all maxillary palp nuclei in Figure 7B,D was plotted as described for the antenna heatmap. The maxillary palp chord plot in Figure 7E was processed similarly to the antenna one. The top 20 most highly expressed chemosensory receptors except for *Gr1* and *Gr2*, together with *Gr3*, were examined for the co-expression at single-nuclei resolution. Co-expressed pairs of receptors found in more than 10 cells were included. Maxillary palp gene expression feature plots were made using the command FeaturePlot on normalized expression values.

### Mosquito preparation for single-sensillum recordings

Female mosquitoes from two wild-type and 3 mutant strains (*+/+^ORL^*, *+/+^LVP^*, *Ir25a^BamHI/BamHI^*, *Gr3^4/4^* and *Orco^16/16^*) five to seven days post-emergence, were anesthetized on wet ice for 1-2 min. An individual mosquito was then glued onto a piece of double-sided sticky tape on a microscope slide (76 × 26 mm) and secured by a piece of tape covering the thorax and abdomen. The maxillary palps were immobilized using a short segment of human hair placed over the basal part of the maxillary palps. The sensilla of the maxillary palps were subsequently visualized using an Olympus light microscope (BX51WI; LRI Instrument AB, Lund, Sweden) at 750×. A continuous humidified stream of synthetic air (Strandmöllen AB, Ljungby, Sweden) was passed over the maxillary palp (2 L min^−1^) via a glass tube (7 mm i.d.), terminating 10 mm from the maxillary palps, to avoid desiccation.

### Single-sensillum recordings from maxillary palp capitate peg sensilla

Electrophysiological recordings from capitate peg sensilla were made and analyzed according to previously described protocols (Ghaninia et al., 2019; Majeed et al., 2016). In brief, two tungsten microelectrodes, electrolytically sharpened in 10% KNO_2_ solution, were used as reference and recording electrodes. The reference electrode was inserted into the eye and the recording electrode was positioned at the base or shaft of the sensillum using a piezo motorized micromanipulator (Märzhäuser Wetzlar GmbH & Co. KG, Wetzlar Germany) until electrical contact was established. Extracellular signals from the olfactory sensory neurons housed in the capitate peg sensilla were amplified and recorded using a high-impedance probe (universal single ended probe) and a USB-acquisition controller (IDAC-4) (Ockenfels Syntech GmbH, Buchenbach, Germany). Extracellular spikes were differentiated based on amplitude as A, B, and C, according to standard nomenclature (Ghaninia et al., 2019; Majeed et al., 2016), and manually counted using Autospike 3.7 (Ockenfels Syntech GmbH). The response to odorant stimuli were analyzed by subtracting the number of spikes 0.5 sec post-stimulus from the number of spikes 0.5 sec pre-stimulus, and the outcome was multiplied by two to obtain a spike/sec measurement. In cases where the neuronal response was high enough to result in pinching of the spike train (>150 spikes/sec), the number of spikes post-stimulus were counted for the first 100 msec and then multiplied by 5, as the inter-spike frequency is constant once the neuron is activated maximally. Neurons were classified as responders or non-responders based upon whether their odorant response was above or below a 30 spikes/sec threshold, respectively.

### Odorant stimulus delivery for single-sensillum recordings

Odorants used in Figure 8 were selected for the highest purity available (>98%): R-(-)-1-octen-3-ol (PubChem CID: 6992244, Penta Manufacturing 15-18900); acetone (PubChem CID: 180 Sigma A4206); hexyl amine (PubChem CID: 8102, Sigma 219703); triethyl amine (PubChem CID: 1146, Sigma T0886). All odorants were diluted into a large stock solution that was used throughout each entire experiment to avoid variability in concentrations. Serial decadic dilutions of acetone, hexyl amine, and triethyl amine were made in MilliQ ultrapure water (18 megaohm resistance) and 1-octen-3-ol was diluted in paraffin oil (EMD Millipore #PX0045-3). Aliquots of 10 µL of each compound and dilution was pipetted onto a piece of filter paper (5 × 20 mm) placed inside a Pasteur pipette. Similar volumes of MilliQ ultrapure water and paraffin oil were used as controls. Stimulus cartridges were used within 5 min after loading, and only used once. For dose-response analysis using CO_2_, gas cylinders containing metered amounts of CO_2_ (300, 600, 1200, 4800 ppm) and oxygen (20%), balanced by nitrogen (Strandmöllen AB, Ljungby, Sweden) were used as previously described (Ghaninia et al. 2019). Odorants were introduced by passing a 0.5 sec air puff through the Pasteur pipette using a stimulus controller (Ockenfels Syntech GmbH) into the airstream passing over the maxillary palps through a hole in the glass tube, 10 cm upstream from the preparation.

### Statistical analysis

All statistical analyses were performed using Prism (GraphPad), Excel (Microsoft) or R version 3.6.3 (R Development CoreTeam, 2017). Data are shown as mean±SEM unless otherwise noted. Details of statistical methods are reported in the figure legends.

## DATA AND RESOURCE AVAILABILITY

Supplementary Figures S1-S14 accompany the paper. Raw data are provided in Data File 1, and additional raw data, plots and analysis, and custom scripts are available at https://github.com/VosshallLab/Younger_Herre_Vosshall2020. Additional raw data and analysis of the snRNA-seq data are on Github at https://github.com/VosshallLab/Younger_Herre_Vosshall2020. These include processed data (Seurat files) and scripts, in addition to descriptive statistics and analysis, including a pseudo-bulk table of gene counts and cell number distribution, violin plots, feature plots, and broad co-expression heatmaps. snRNA-seq sequencing reads are freely available from the Gene Expression Omnibus (accession number GEO: GSE192978). Additional files related to snRNA-seq data analysis and visualization are available at Zenodo (antenna: https://zenodo.org/record/5818543#.YdX7VWjMIuU; maxillary palp: https://zenodo.org/record/5818952#.YdX7KWjMIuU, Plasmids are available from Addgene (accession #162520-162526).

---

**Figure S1.**
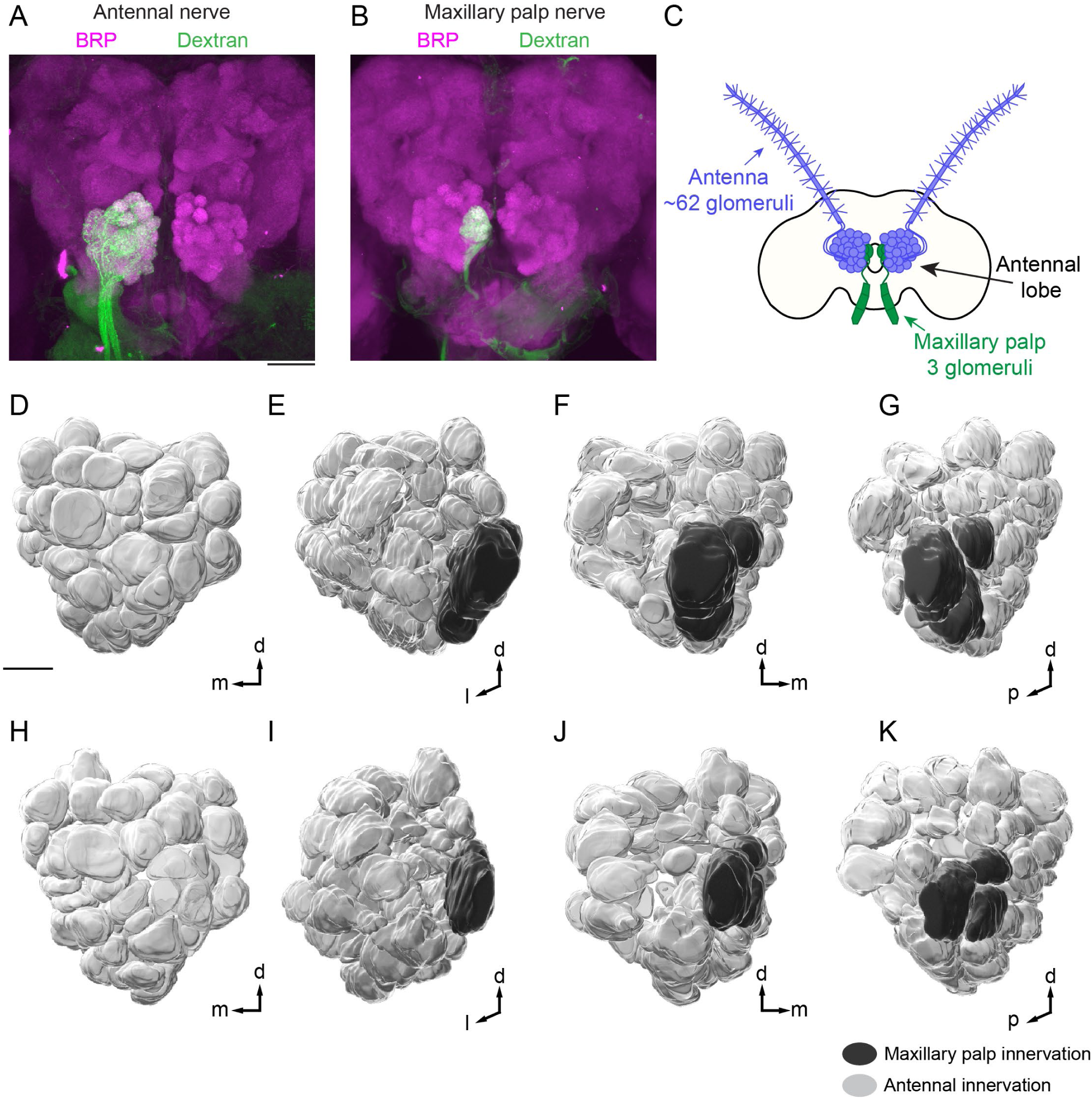
Organization of *Aedes aegypti* antennal lobe glomeruli (Related to Figure 1) (**A-B**) Maximum-intensity projections of confocal Z-stacks of a brain after anterograde dye fill of a single ipsilateral antenna (A) or ipsilateral maxillary palp (B) using a dextran-conjugated fluorophore (green) with immunofluorescent labeling of Brp (synaptic marker, magenta). (**C**) Approximate number of antennal lobe glomeruli per brain hemisphere innervated by the indicated sensory structure, derived from quantification of the left antennal lobe in 12 brains presented in Figure 1I-J, Figure S2-S5. (**D-K**) 3-D reconstruction of a single left antennal lobe with 61 (D-G) or 66 (H-K) glomeruli shown at 4 different angles. Glomeruli are colored according to innervation by the indicated sensory appendage. Panel (G) is reprinted in Figure 5B. Scale bars: 50 µm (A-B), 20 µm (D-K). Orientation: d=dorsal, m=medial, p=posterior.

**Figure S2.**
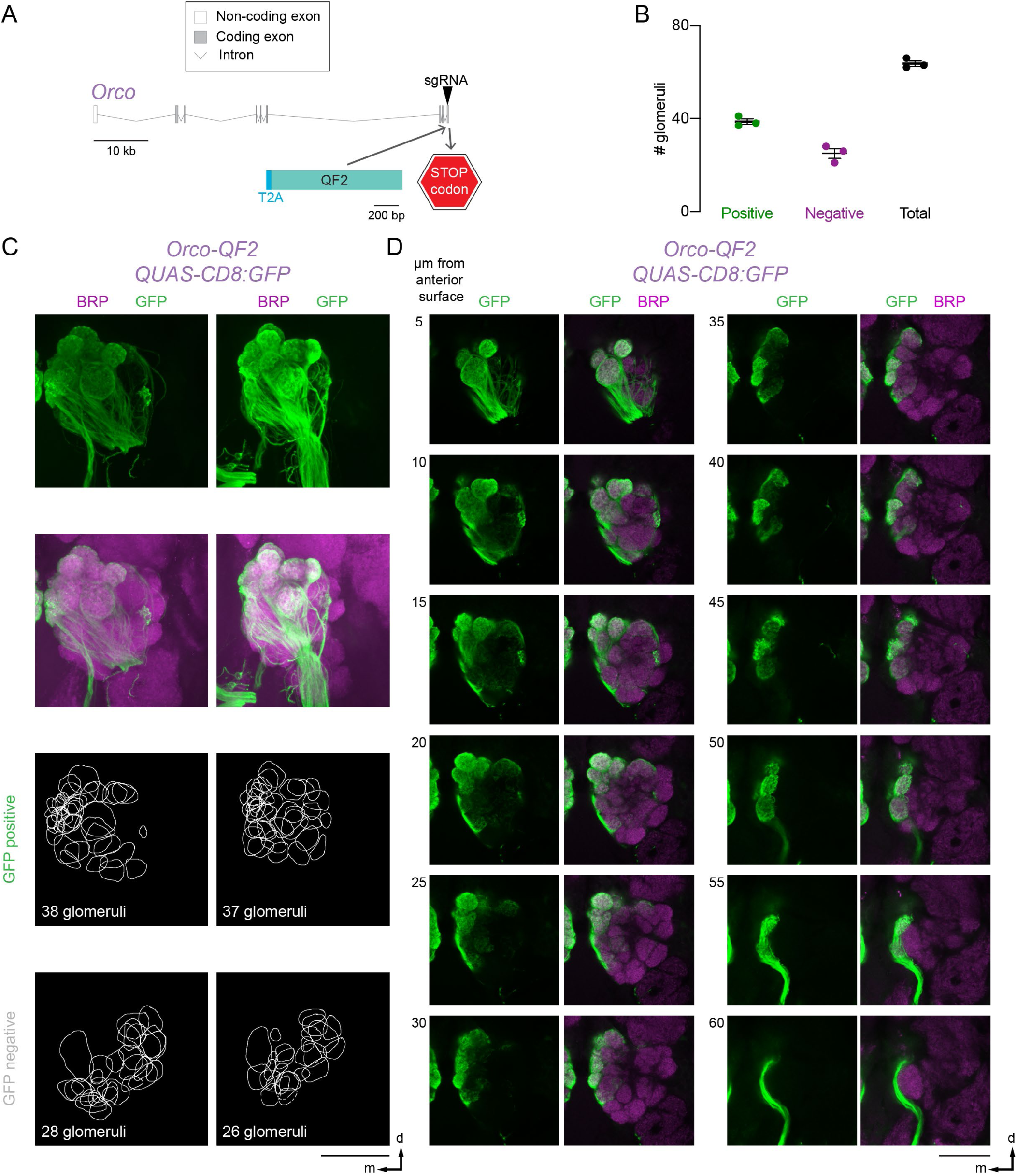
Projections of *Orco-QF2*-expressing neurons in the antennal lobe (Related to Figure 1) **(A)** *Orco* locus with exons (grey boxes), introns (grey lines) and CRISPR-Cas9 gRNA site (arrowhead) used to insert T2A-QF2 (light blue). (**B**) Quantification of the number of glomeruli that are GFP positive (green), GFP negative (magenta), and total number of glomeruli (black). Analysis based on brains in (C-D) and Figure 1I,J. (**C**) Maximum-intensity projections of confocal Z-stacks of left antennal lobes from two different brains of the indicated genotype with immunofluorescent labeling of GFP (green) and Brp (synaptic marker, magenta) (top) and 2-D representation of the boundary of each glomerulus that is GFP positive and GFP negative (bottom). (**D**) Single confocal sections taken from the maximum-intensity projection confocal Z-stack of the left antennal lobe shown in Figure 1I with immunofluorescent labeling of GFP (green) and Brp (synaptic marker, magenta). A single plane is shown every 5 µm in Z to capture each glomerulus. Scale bar (C-D): 50 µm. Orientation: d=dorsal, m=medial.

**Figure S3:**
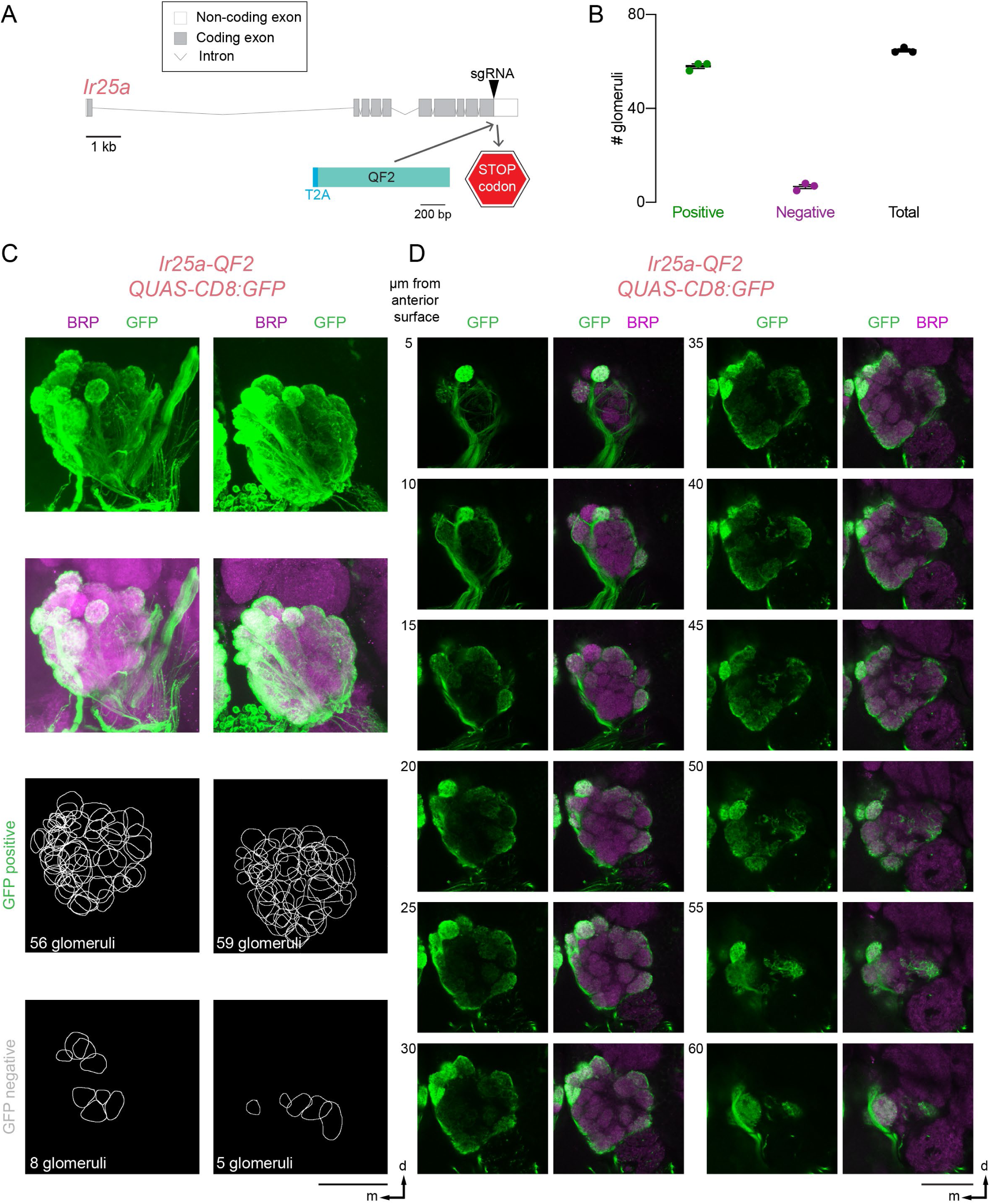
Projections of *Ir25a-QF2*-expressing neurons in the antennal lobe (Related to Figure 1) (**A**) *Ir25a* locus with exons (grey boxes), introns (grey lines) and CRISPR-Cas9 gRNA site (arrowhead) used to insert T2A-QF2 (light blue). (**B**) Quantification of the number of glomeruli that are GFP positive (green), GFP negative (magenta), and total number of glomeruli (black). Analysis based on brains in (C-D) and Figure 1I,J. (**C**) Maximum-intensity projections of confocal Z-stacks of left antennal lobes from two different brains of the indicated genotype with immunofluorescent labeling of GFP (green) and Brp (synaptic marker, magenta) (top) and 2-D representation of the boundary of each glomerulus that is GFP positive and GFP negative (bottom). (**D**) Single confocal sections taken from the maximum-intensity projection confocal Z-stack of the left antennal lobe shown in Figure 1I with immunofluorescent labeling of GFP (green) and Brp (synaptic marker, magenta). A single plane is shown every 5 µm in Z to capture each glomerulus. Scale bar (C-D): 50 µm. Orientation: d=dorsal, m=medial.

**Figure S4:**
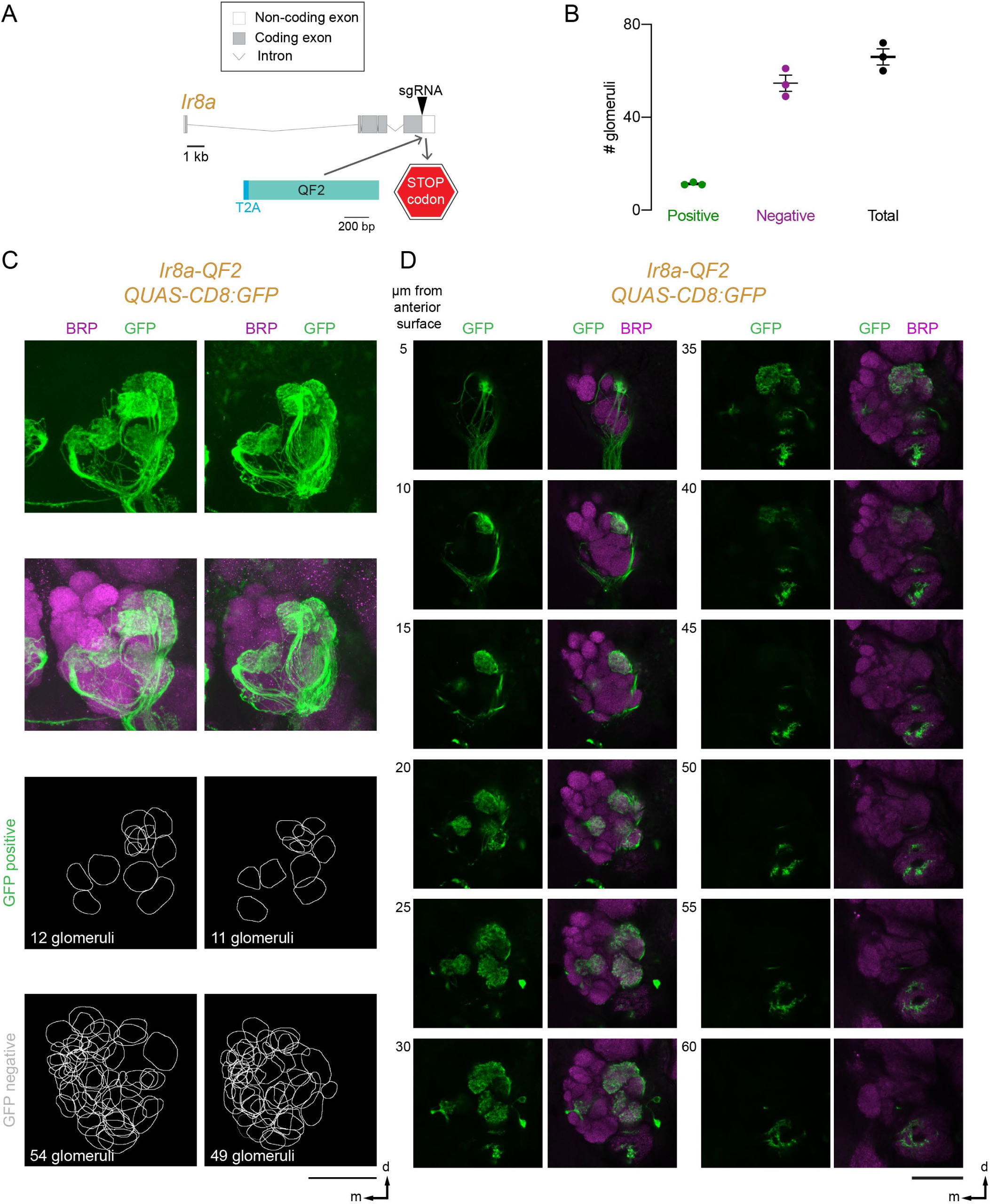
Projections of *Ir8a-QF2*-expressing neurons in the antennal lobe (Related to Figure 1) (**A**) *Ir8a* locus with exons (grey boxes), introns (grey lines) and CRISPR-Cas9 gRNA site (arrowhead) used to insert T2A-QF2 (light blue). (**B**) Quantification of the number of glomeruli that are GFP positive (green), GFP negative (magenta), and total number of glomeruli (black). Analysis based on brains in (C-D) and Figure 1I,J. (**C**) Maximum-intensity projections of confocal Z-stacks of left antennal lobes from two different brains of the indicated genotype with immunofluorescent labeling of GFP (green) and Brp (synaptic marker, magenta) (top) and 2-D representation of the boundary of each glomerulus that is GFP positive and GFP negative (bottom). (**D**) Single confocal sections taken from the maximum-intensity projection confocal Z-stack of the left antennal lobe shown in Figure 1I with immunofluorescent labeling of GFP (green) and Brp (synaptic marker, magenta). A single plane is shown every 5 µm in Z to capture each glomerulus. Scale bar (C-D): 50 µm. Orientation: d=dorsal, m=medial.

**Figure S5:**
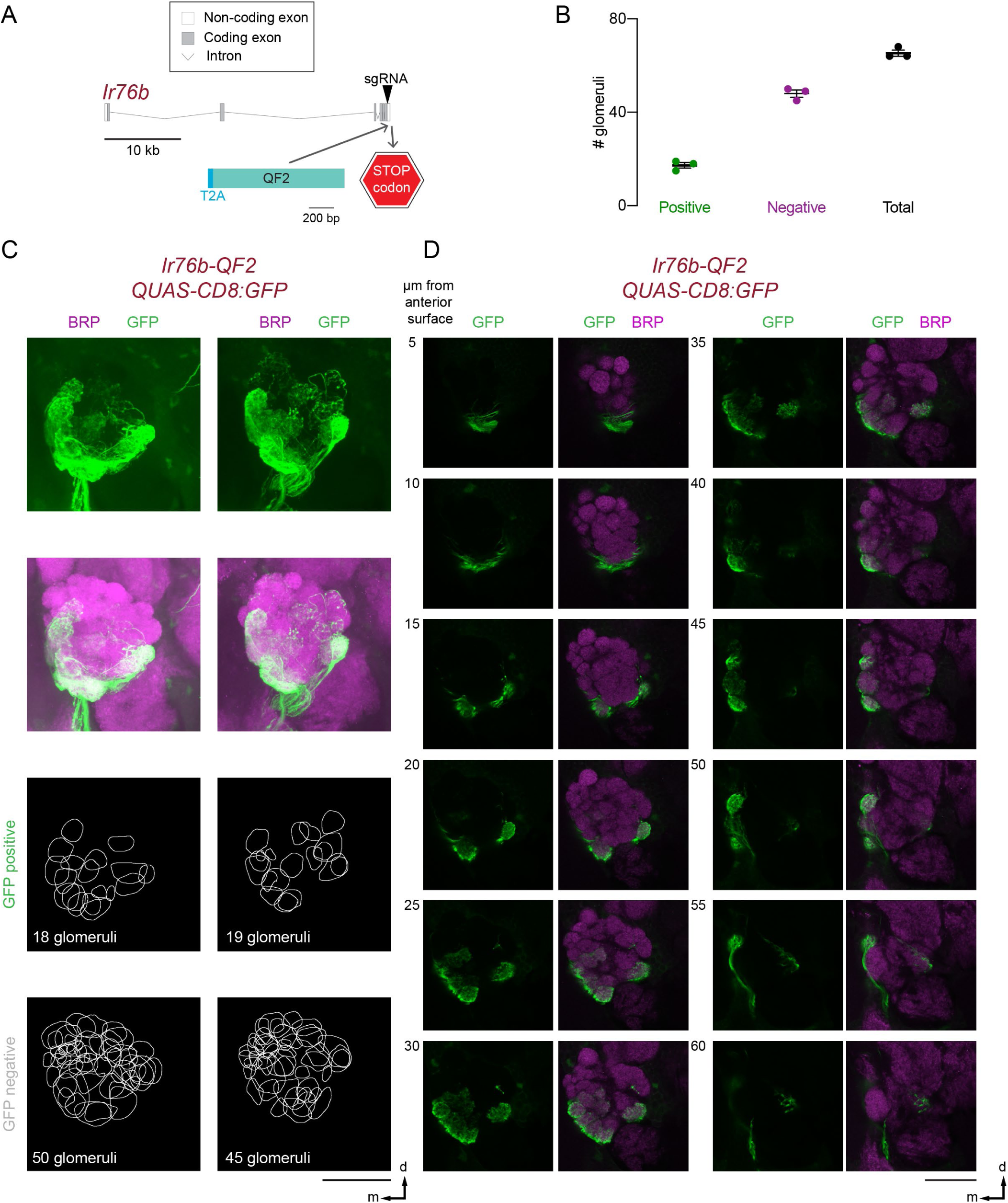
Projections of *Ir76b-QF2*-expressing neurons in the antennal lobe (Related to Figure 1) (**A**) *Ir76b* locus with exons (grey boxes), introns (grey lines) and CRISPR-Cas9 gRNA site (arrowhead) used to insert T2A-QF2 (light blue). (**B**) Quantification of the number of glomeruli that are GFP positive (green), GFP negative (magenta), and total number of glomeruli (black). Analysis based on brains in (C-D) and Figure 1I,J. (**C**) Maximum-intensity projections of confocal Z-stacks of left antennal lobes from two different brains of the indicated genotype with immunofluorescent labeling of GFP (green) and Brp (synaptic marker, magenta) (top) and 2-D representation of the boundary of each glomerulus that is GFP positive and GFP negative (bottom). (**D**) Single confocal sections taken from the maximum-intensity projection confocal Z-stack of the left antennal lobe shown in Figure 1I with immunofluorescent labeling of GFP (green) and Brp (synaptic marker, magenta). A single plane is shown every 5 µm in Z to capture each glomerulus. Scale bar (C-D): 50 µm. Orientation: d=dorsal, m=medial.

**Figure S6.**
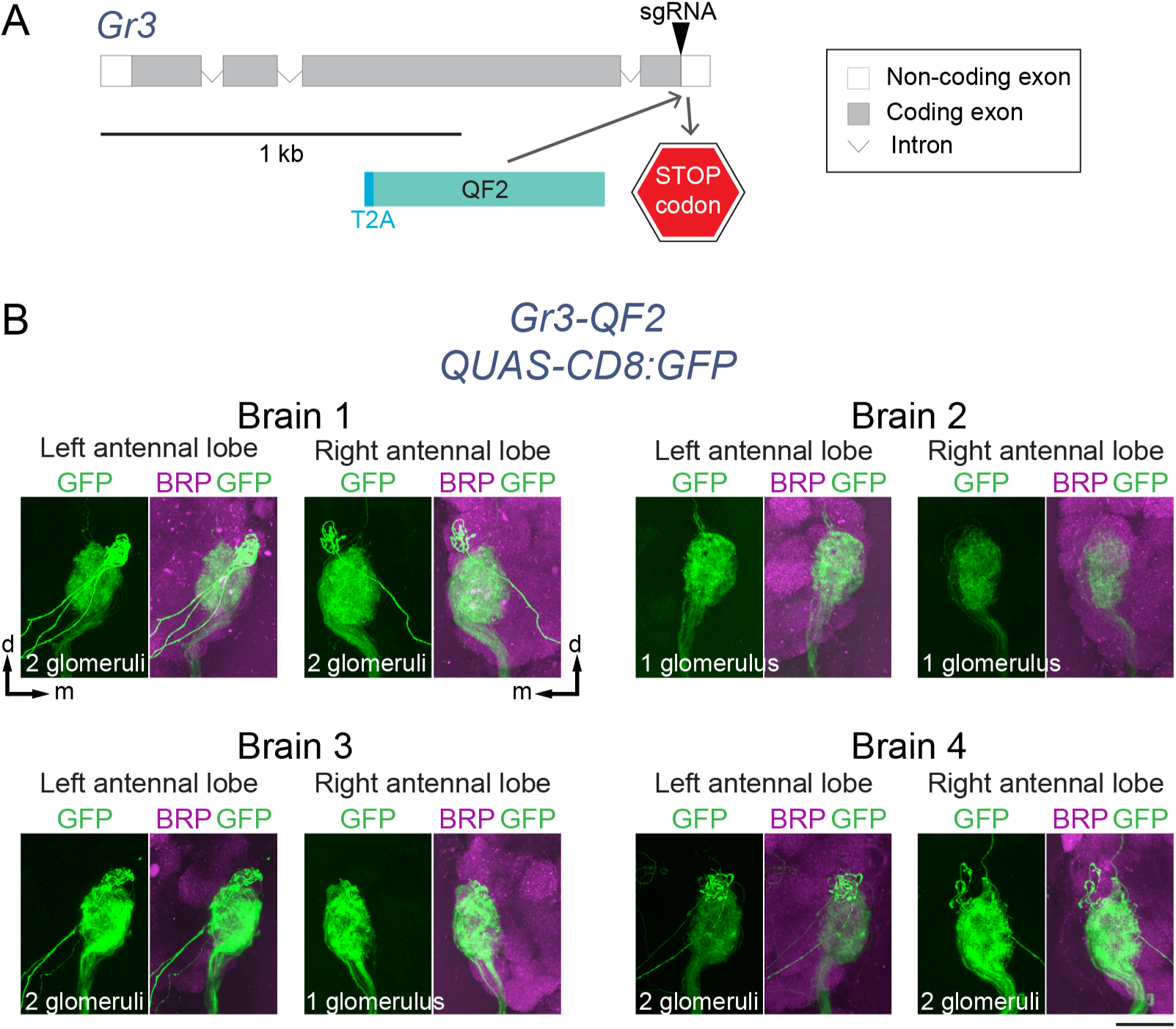
Projections of *Gr3-QF2*-expressing neurons in the antennal lobe (Related to Figure 1) (**A**) *Gr3* locus with exons (grey boxes), introns (grey lines) and CRISPR-Cas9 gRNA site (arrowhead) used to insert T2A-QF2 (light blue). (**B**) Maximum-intensity projection confocal Z-stack through the medial antennal lobes of 4 brains with immunofluorescent labeling of GFP (green) and Brp (synaptic marker, magenta). Scale bar: 25 µm. Orientation: d=dorsal, m=medial.

**Figure S7.**
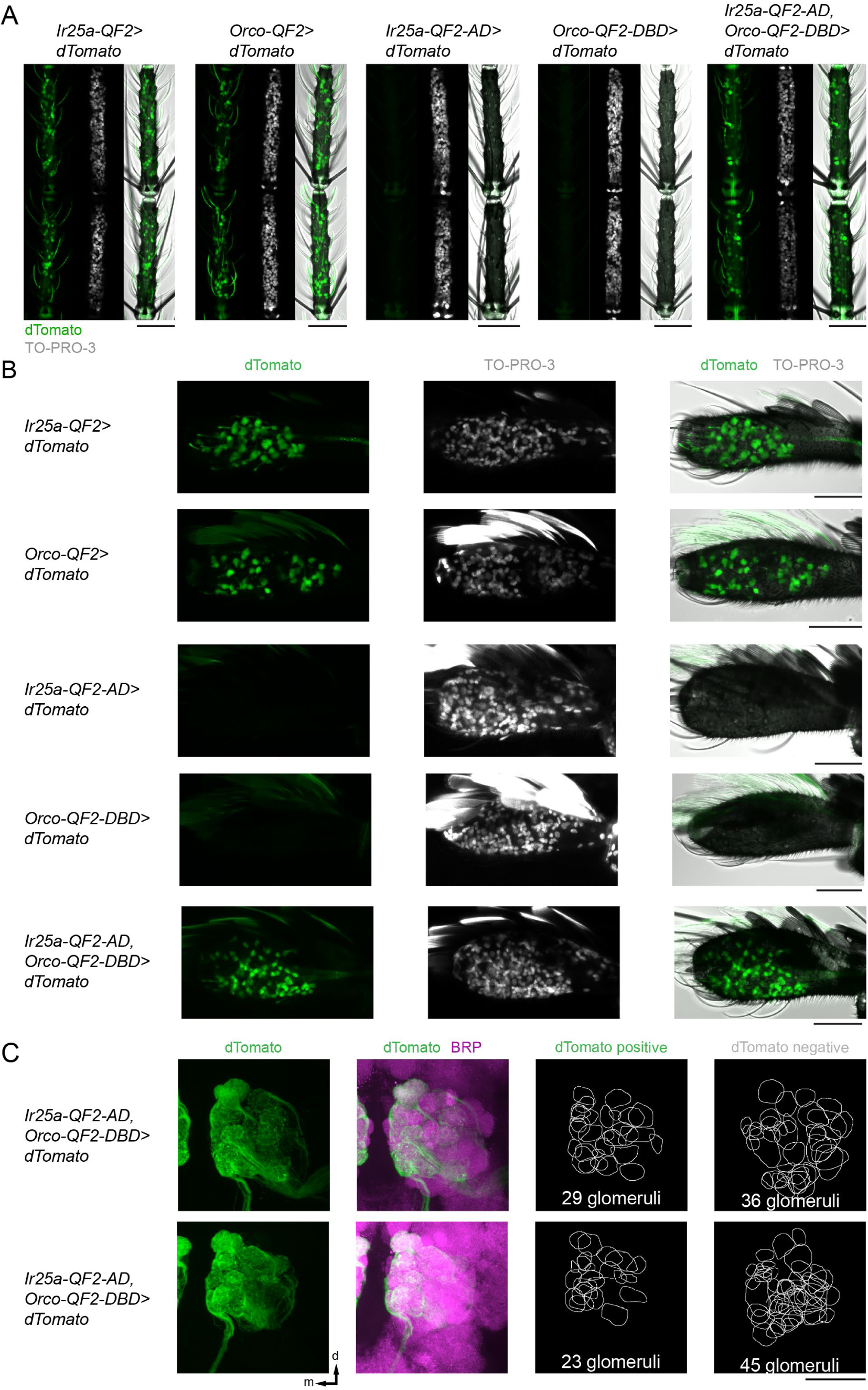
Specificity of Split-QF2 reagents (Related to Figure 2) (**A-B**) Maximum-intensity projections of confocal Z-stacks of antennae (A) and maxillary palps **(B)** of the indicated genotypes showing intrinsic dTomato fluorescence and stained with the nuclear dye TO-PRO-3, with transmitted light overlay. (**C**) Maximum-intensity projections of confocal Z-stacks of left antennal lobes from two different brains of the indicated genotype with immunofluorescent labeling of dTomato (green) and Brp (synaptic marker, magenta) (top) and 2-D representation of the boundary of each glomerulus that is GFP positive and GFP negative (bottom). Scale bars: 50 µm. Orientation (C,D): d=dorsal, m=medial.

**Figure S8.**
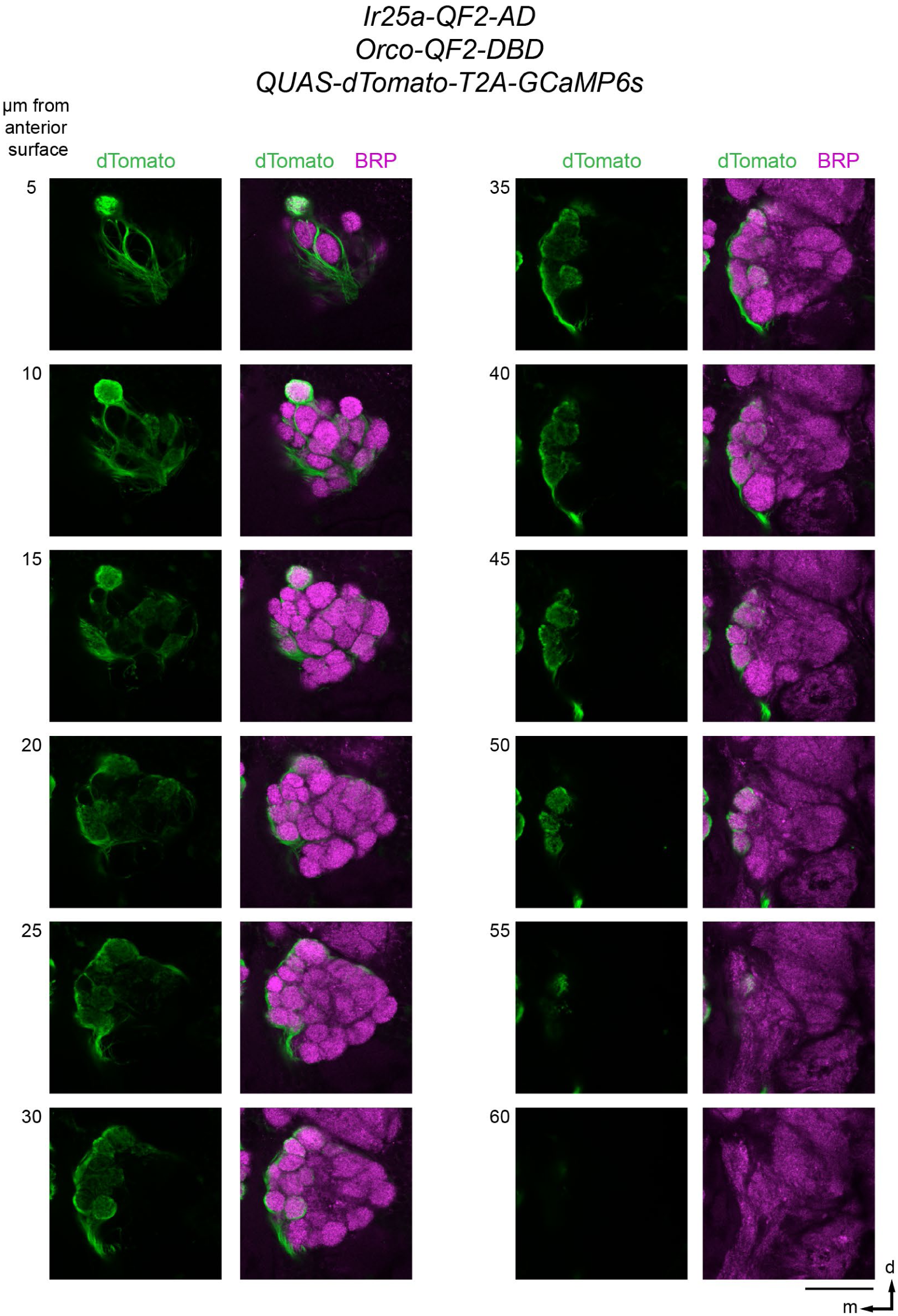
Projections of *Orco-QF2-DBD*; *Ir25a-QF2-AD*-expressing neurons in a single antennal lobe (Related to Figure 2) Single confocal sections taken from the maximum-intensity projection confocal Z-stack of the left antennal lobe shown in Figure 2G with immunofluorescent labeling of dTomato (green) and Brp (synaptic marker, magenta). A single plane is shown every 5 µm in Z to capture each glomerulus. Scale bar: 50 µm. Orientation: d=dorsal, m=medial.

**Figure S9:**
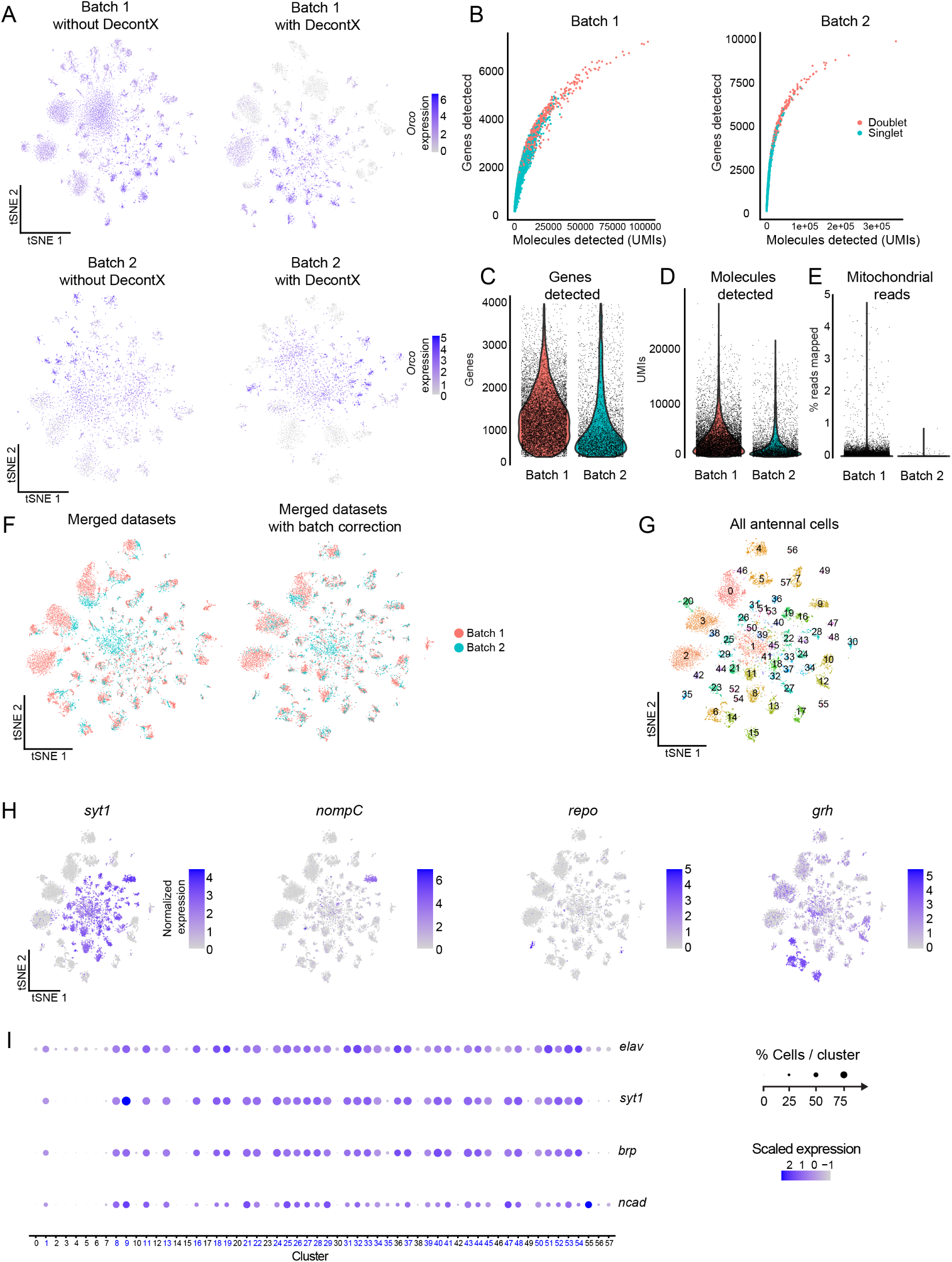
Antennal snRNA-seq ambient RNA removal, filtering, batch correction and neuron-identification (Related to Figure 5) (**A**) Ambient RNA removal using DecontX was used independently on data from snRNA-seq experiments processed at Rockefeller (Batch 1) and Baylor (Batch 2), illustrated using normalized expression of *Orco* mapped onto t-distributed stochastic neighbor embedding (t-SNE) plots. Normalized Expression: log(UMI of gene*10,000 / total UMI of cell +1). (**B**) Identification of multiplets for removal using DoubletFinder. Pearson Correlation coefficient of genes and counts was 0.89 for Batch 1 and 0.82 for Batch 2. (**C-E**) Sample properties and distributions after filtering with DecontX. Nuclei that were retained expressed between 400 and 40000 genes (C) and fewer than 5% mitochondrial transcripts (E). Nuclei were not additionally filtered on UMIs after multiplet removal (D). (**F**) Independently collected snRNA-seq experiments were merged and batch effects reduced. (**G**) Antennal cell clusters after batch effect reduction, visualized using t-SNE. (**H**) Normalized expression [log(UMI of gene*10,000 / total UMI of cell +1)] mapped onto t-SNE plots for *syt1* as a marker for neurons*, nompC* for mechanosensory cells*, repo* for glial cells, and *grh* for epithelial cells. (**I**) Dot plot of neural markers used to identify neuron clusters. Clusters were identified as neurons if over 50% of cells within a cluster expressed 3 out of 4 neural markers (*elav*, *syt1*, *brp*, *ncad*). The 19 neuron clusters that were identified are labeled in blue. Mean scaled expression in cluster: Z-score.

**Figure S10:**
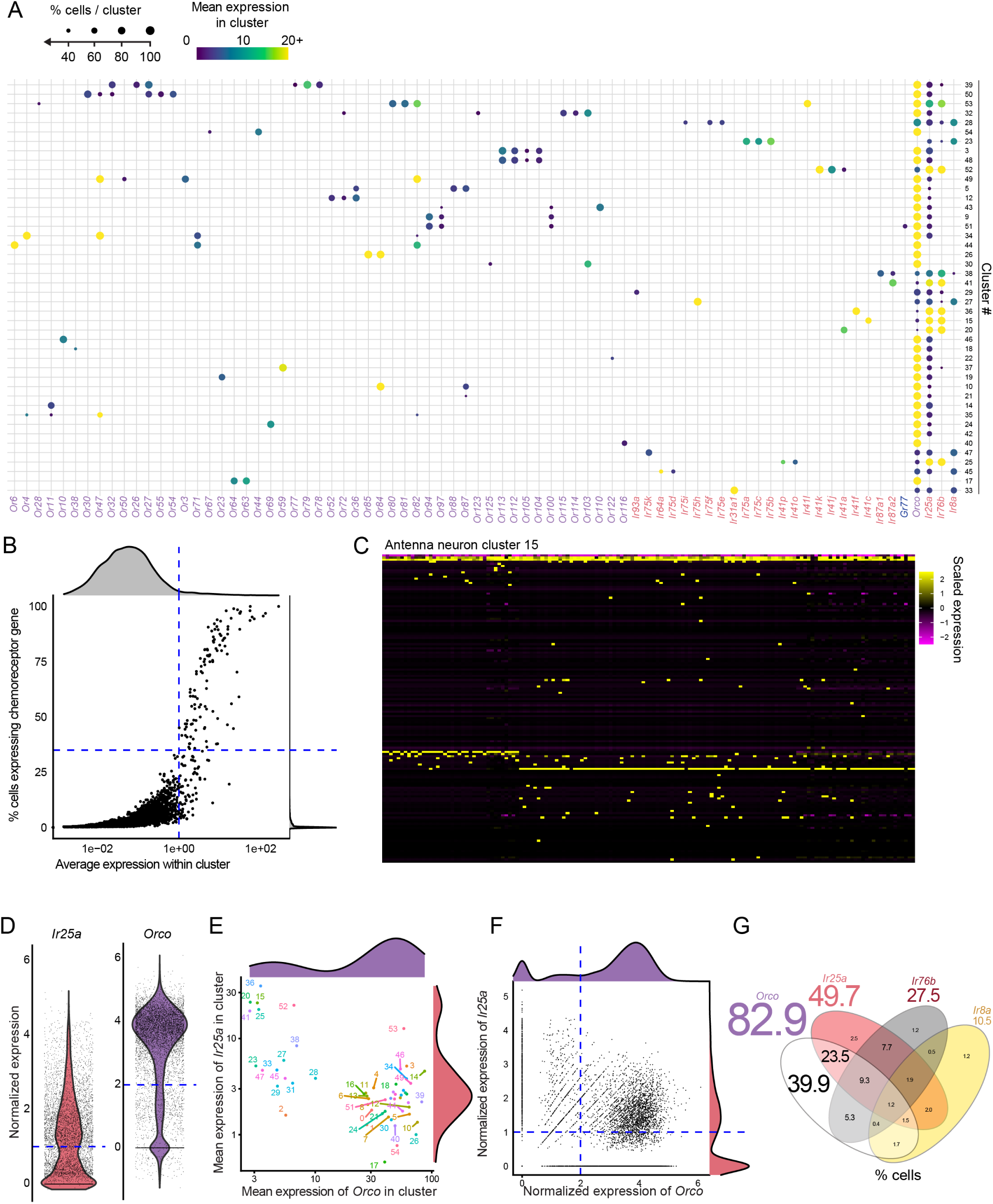
Antennal snRNA-seq cluster chemosensory receptor expression analysis and co-receptor analysis (Related to Figure 5) (**A**) Dot plot summarizing chemosensory receptor expression in neuron clusters. The circle size represents the % of cells in each cluster that express a given gene above criteria illustrated in (B). (**B**) Distribution of chemosensory receptor genes within clusters used for thresholding in the dot plot in (A). Points denote expression patterns of individual chemosensory receptor genes listed on x-axis of (B) for each cluster. Lines indicate mean normalized expression level of 1 within cells of the cluster and 35% of cells expressing a chemosensory receptor gene in a cluster. Genes in upper and lower right hand-quadrant was included for depiction in the dot plot (B). (**C**) Example co-expression heatmap of 148 cells within antenna neuron cluster 15, demonstrating distinct combinations of chemosensory receptor expression in groups of cells within one cluster. Scaled expression: Z-score. (**D-G**) Co-expression of chemosensory co-receptors. To determine co-expression, a normalized expression threshold of log(UMI of gene*10,000 / total UMI of cell +1) was used for *Ir25a, Ir76b, Ir8a.* Due to higher expression levels, a threshold of 2 log(UMI of gene*10,000 / total UMI of cell +1) was used for *Orco* (D). Scatter plots of *Orco* and *Ir25a* expression within neuron clusters (E) and individual cells (F). Venn diagram depicting percent of neurons co-expressing different combinations of co-receptors according to normalized expression [log(UMI of gene*10,000 / total UMI of cell +1)]. Larger numbers outside of the Venn diagram indicate total percent of neurons expressing indicated co-receptor (G).

**Figure S11:**
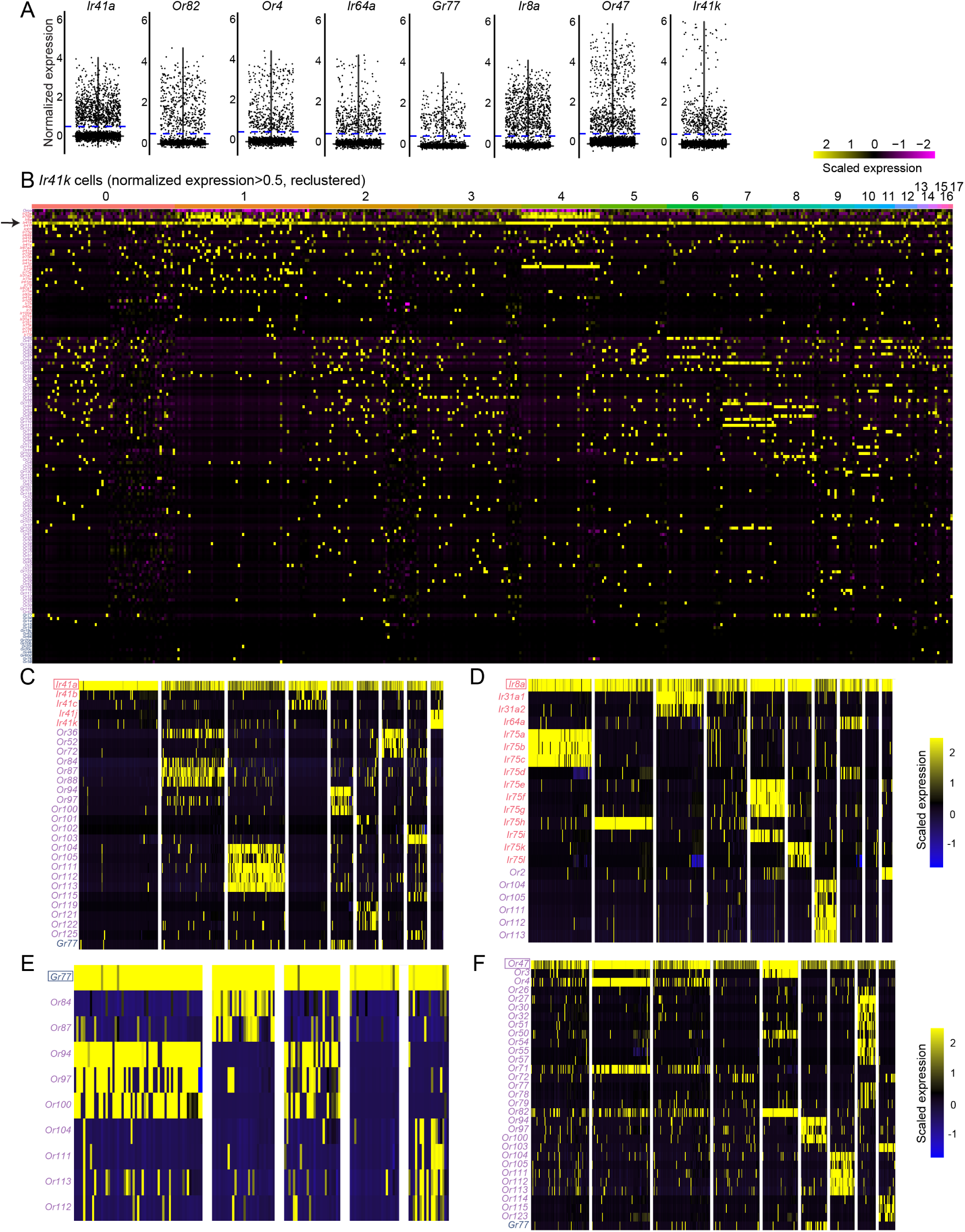
Antennal snRNA-seq cluster chemosensory receptor expression analysis and co-receptor analysis (Related to Figure 5) (**A**) Violin plots illustrating the expression distribution of selected genes used for co-expression simplified heatmaps in Figure 5F-I and this figure. Normalized Expression: log(UMI of gene*10,000 / total UMI of cell +1), (C-F). A normalized expression threshold of 0.5 log(UMI of gene*10,000 / total UMI of cell +1) used to identify cells expressing a given chemosensory receptor (indicated by dotted blue line). (**B**) Example co-expression heatmap of 412 cells with *Ir41k* as filtered in (A). Columns represent individual cells, sorted by clustering. Rows represent chemosensory receptor genes, ordered first by chemosensory receptor family, then mean expression level in cluster 0. Cells from clusters 2, 3, 4, 6, 7, and 8 were selected as illustrating examples of co-expression for simplified heatmap visualization in Figure 5F. Genes were selected manually based on expression level and number of cells exhibiting a given co-expression pattern. Scaled expression: Z-score. (**C-F**) Simplified heatmaps illustrating co-expression patterns for (C) *Ir41a,* (D) *Ir8a,* (E) *Gr77*, and (F) *Or47* using selected cells and visually-identified genes via clustering similar to (C). Scaled expression: Z-score.

**Figure S12.**
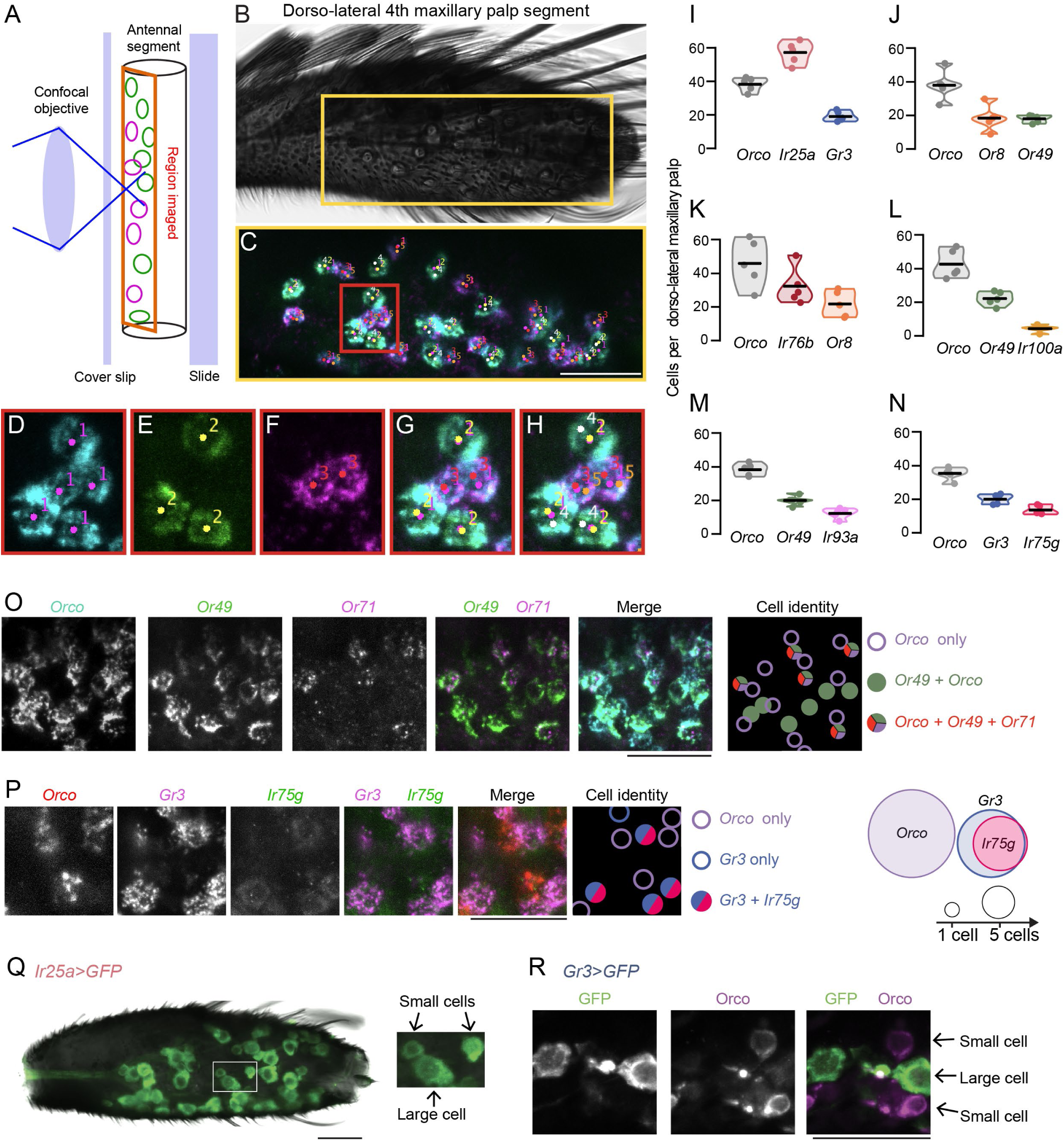
Quantification of maxillary palp cell populations (Related to Figure 6) (**A-H**) Workflow for cell quantification. Schematic of antennal region imaged on a confocal microscope (A) and image of maxillary palp with imaged area indicated with the yellow square (B). Whole-mount maxillary palp RNA *in situ* hybridization, yellow region from (C). Cells are manually marked independently as *Orco*+, *Or49*+, or *Or8*+ (red inset from B) using FIJI Cell Counter (D-F) and markers from each channel are merged (G). Cells with markers 1 and 2 are then scored as *Orco*+*Or49*+ with marker 4, and cells with markers 1 and 3 are then scored as *Orco*+*Or8*+ with marker 5 (H). Counts from each marker for each image are exported into Excel and R for further analysis. (**I-N**) Total cell counts from whole mount maxillary palp RNA *in situ* hybridization in Figure 6. Mean with range, n=5. (**O-P**) RNA *in situ* hybridization of whole-mount maxillary palp with the indicated probe and cell identity schematic. (**Q-R**) Whole-mount maxillary palp immunostaining showing *Ir25a* expression in “small” and “large” cells (Q) and *Gr3* expression in “large” cells and Orco protein in “small” cells (R). Scale bars: 25 µm except (C): 50 µm. Orientation (B, P): proximal left.

**Figure S13:**
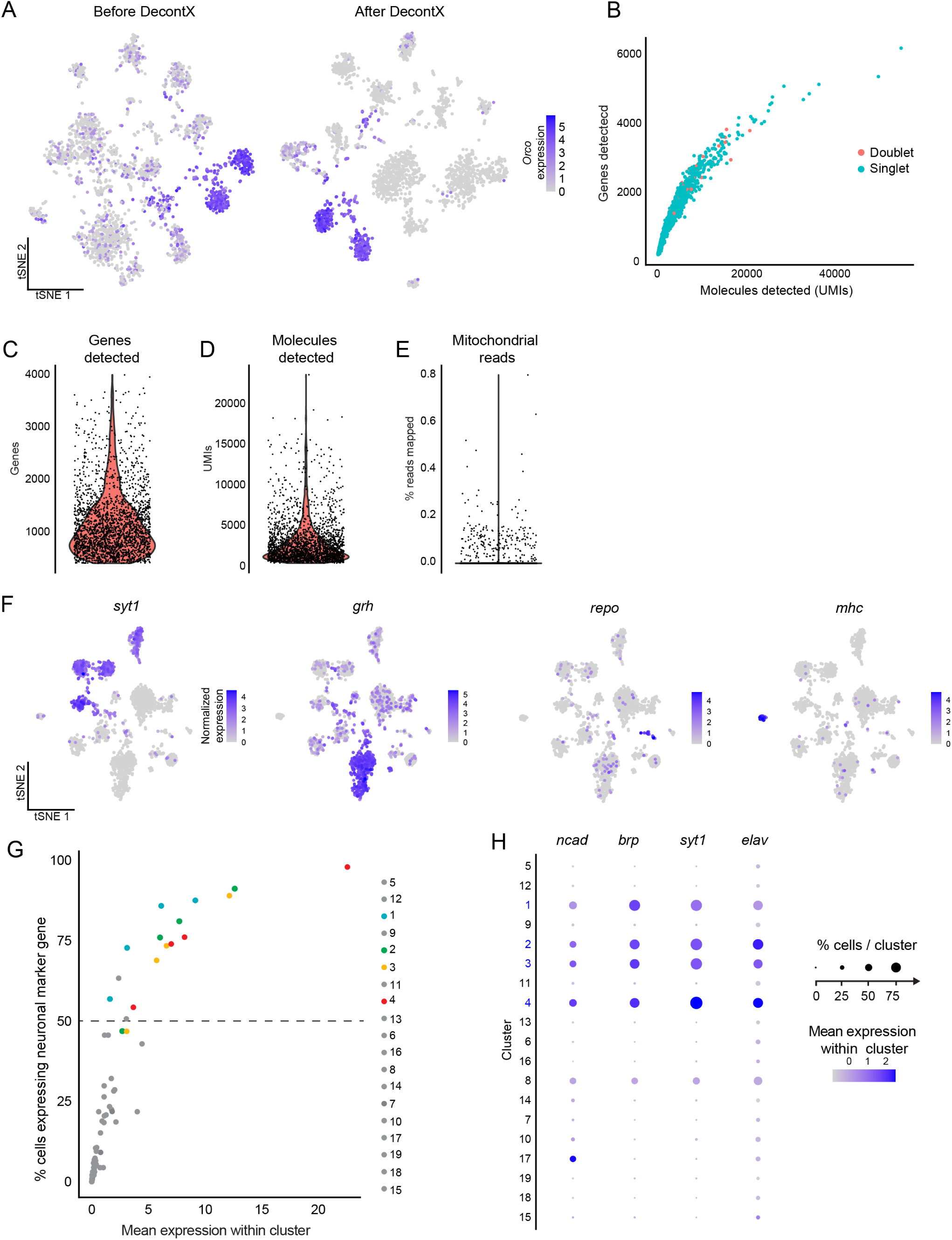
Maxillary palp snRNA-seq ambient RNA removal, filtering, and neuron-identification (Related to Figure 7) (**A**) Ambient RNA removal using DecontX, illustrated using normalized expression of *Orco* mapped onto t-distributed stochastic neighbor embedding (t-SNE) plots for maxillary palp snRNA-seq experiment. Normalized Expression: log(UMI of gene*10,000 / total UMI of cell +1). (**B**) Identification of multiplets for removal using DoubletFinder. Pearson Correlation coefficient of genes and counts was 0.93. (**C-E**) Sample properties and distributions after filtering. Nuclei were retained that expressed between 400 and 40000 genes (C) and fewer than 5% mitochondrial transcripts (E). Nuclei were not additionally filtered on UMIs after multiplet removal (D). (**F**) Normalized expression [log(UMI of gene*10,000 / total UMI of cell +1)] mapped onto t-SNE plots for *syt1* as a marker for neurons*, grh* for epithelial cells, *repo* for glial cells, and *mhc* for muscle cells. (**G**) Distribution of neural marker genes (*ncad, brp, syt1*, and *elav*) within clusters. Points denote expression patterns of individual neural marker genes for each cluster. Line indicates the threshold used to identify neuron clusters, with 50% of cells within a cluster expressing a 3 out of 4 defined neural markers. Mean expression in cluster: UMI of gene*10,000 / total UMI of cell +1. (**H**) Dot plot of neural markers used to identify neuron clusters. Clusters of identified neurons are identified as clusters 1, 2, 3, and 4. Mean scaled expression in cluster: Z-score.

**Figure S14:**
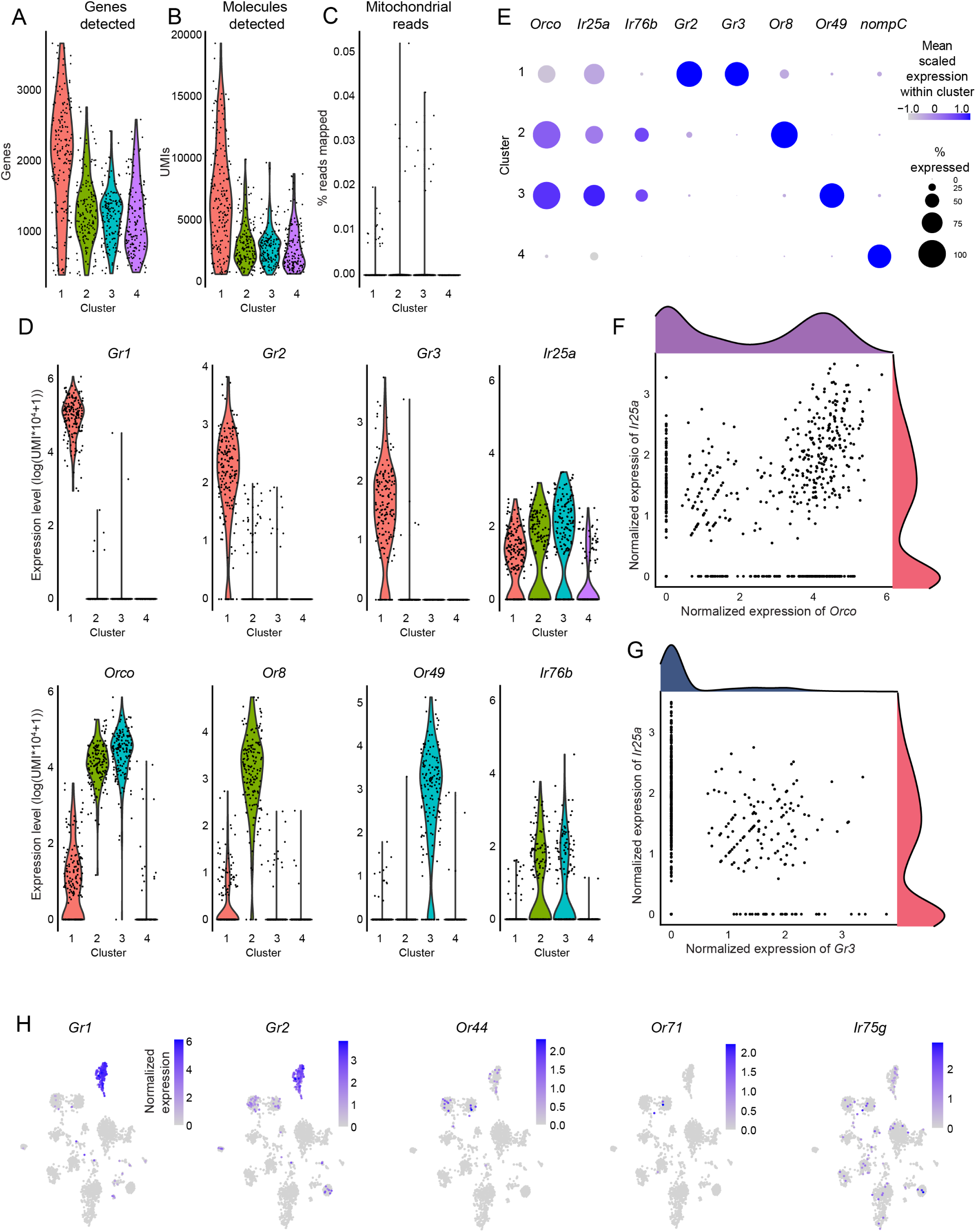
Maxillary palp snRNA-seq cluster chemosensory receptor expression analysis and co-receptor analysis on four identified neuron cluster populations (Related to Figure 7) (**A**) Number of genes detected per cell in neuron clusters defined in Figure S13G-H. (**B**) Number of transcripts detected per cell in neuron clusters. (**C**) Percent mitochondrial reads per cell in neuron clusters. (**D**) Distribution of normalized expression levels of the indicated genes in cells within neuronal clusters. Normalized Expression: log(UMI of gene*10,000 / total UMI of cell +1). (**E**) Dot plot illustrating mean scaled expression (Z-score) and cells expressing a given gene. (**F-G**) Scatter plot depicting expression levels within individual neuron-identified cells of *Orco* and *Ir25a* (F) and *Gr3* and *Ir25a* (G). Normalized Expression: log(UMI of gene*10,000 / total UMI of cell +1). (**H**) Feature plots showing normalized expression mapped onto t-SNE plots for the indicated genes. Normalized Expression: log(UMI of gene*10,000 / total UMI of cell +1).

